# Mechanistic and Antigenic Boundaries of *Henipavirus* and *Parahenipavirus* Glycoproteins

**DOI:** 10.1101/2024.12.11.627382

**Authors:** Aaron J. May, Muralikrishna Lella, Jared Lindenberger, Alex Berkman, Ujjwal Kumar, Moumita Dutta, Maggie Barr, Rob Parks, Xiaozhi Lu, Madison Berry, Amanda Newman, Xiao Huang, Arpita Mrigwani, Kijun Song, Victor Ilevbare, Salam Sammour, Chan Soo Park, Radha Devkota Adhikari, Priyanka Devkota, Katarzyna Janowska, Yanshun Liu, Garrett Scapellato, Taylor N. Spence, Katayoun Mansouri, Kevin Wiehe, Robert J Edwards, Kevin O. Saunders, Barton F. Haynes, Priyamvada Acharya

**Author notes:** Correspondence (A.M.), (P.A.). These authors contributed equally (Muralikrishna Lella, Jared Lindenberger, Alex Berkman).

## Abstract

Henipaviruses, a genus within the *Paramyxoviridae* family, include the highly virulent Nipah and Hendra viruses that cause reoccurring outbreaks of deadly disease. Recent discoveries of several new *Paramyxoviridae* species, including the zoonotic Langya virus, have revealed much higher antigenic diversity than currently characterized and prompted the reorganization of these viruses into the *Henipavirus* and *Parahenipavirus* genera. Here, to explore the limits of structural and antigenic variation in both genera, collectively referred to here as HNVs, we constructed an expanded, antigenically diverse panel of HNV fusion and attachment glycoproteins from 56 unique HNV strains that better reflects global HNV diversity. We expressed and purified the fusion protein ectodomains and the attachment protein head domains and studied their biochemical, biophysical and structural properties. We performed immunization experiments in mice leading to the elicitation of antibodies reactive to multiple HNV fusion proteins. Cryo-electron microscopy structures of diverse fusion proteins elucidated molecular determinants of differential pre-fusion state metastability and higher order contacts. A crystal structure of the Gamak virus attachment head domain revealed an additional domain appended to the conserved 6-bladed, β-propeller fold. Taken together, these studies expand the known structural and antigenic limits of the HNVs, reveal new cross-reactive epitopes within both genera and provide foundational data for the development of broadly reactive countermeasures.

## Introduction

Henipaviruses (HNVs) are a genus of single-stranded RNA (ssRNA) viruses that includes the highly virulent Nipah virus (NiV) responsible for reoccurring outbreaks of deadly disease. The *Henipavirus* genus is a part of the *Paramyxoviridae* family, and therefore related to other notable human pathogens such as the parainfluenza viruses, measles, and mumps ^1^. The combined possibility within the family for both rapid transmission, as with measles, and high lethality, as with NiV, further highlights pandemic risk and necessitates urgent research to establish preparedness that will enable a rapid and effective response to a future emergent threat. For HNVs specifically, their identification in diverse animal reservoirs in different geographical locations, their high risk of zoonotic transmission, and current lack of approved vaccines or therapies to treat HNV infection in humans highlights the high risk and pandemic potential of this genus.

HNVs have been studied since the discovery of Hendra (HeV) and Nipah viruses in 1994 and 1998, respectively ^2–4^. Additional members of the genus have been identified since then, but it was not until 2022 with the discovery of Langya virus (LayV) that another species with pathogenicity in humans was confirmed ^5^. Like HeV and NiV, LayV was of zoonotic origin. However, unlike HeV, NiV, and most of the other known HNVs at the time, LayV was found to have likely originated from a shrew reservoir, rather than from fruit bats. Several concurrent studies identified more shrew-borne HNVs over a wide geographic range, indicating the existence of a largely uncharacterized clade within the Henipavirus genus ^6–8^. Among these studies were some untargeted surveillance efforts, revealing new HNV sequences among dozens of other sequences across multiple virus families ^9^. While the newly discovered shrew-borne species had initially been classified as members of the *Henipavirus* genus, they have recently been reclassified into the *Parahenipavirus* genus ^10^. In this study, we use the abbreviation HNV to refer collectively to species of both genera.

The HNV attachment (G) and fusion (F) surface glycoproteins together facilitate virus entry into host cells, with G responsible for receptor binding and F mediating viral and host membrane fusion. During this process, both G and F undergo important conformational changes, including destabilization of the prefusion conformation of F by G. The specifics of these conformational steps, including how G destabilizes the pre-fusion F conformation or “triggers” its pre-fusion to post-fusion transition remain elusive. Although a class I fusion protein ^11^, the HNV-F protein, along with all *Paramyxoviridae* F proteins, differs from the canonical class I fusion mechanism in that there is a separation of the receptor attachment function into a separate protein ^12^. Whereas other viral fusion proteins proceed through the fusion conversion process by repositioning or removal of their attachment subunits, HNV-F proteins retain inherent metastability through F protein architecture alone.

In addition to their role in virus entry, F and G are the sole targets for neutralizing antibodies on the surface of HNVs, and thus are important from the perspective of preventative and therapeutic countermeasure development ^13^. The antigenicity of the F and G proteins, outside those of the more well-studied Nipah and Hendra viruses, and the possibility of cross-reactivity across the genus remains largely unexplored. While there are several known monoclonal antibodies (mAbs) that are able to neutralize both NiV and HeV ^14–18^, so far none of these have been found to have any reactivity with LayV ^19,20^. Of the few mAbs known to bind to LayV, some have been noted to be cross-reactive with Mojiang virus (MojV), but not to NiV or HeV ^20^. Whether there are any broadly reactive anti-HNV antibodies or where the limits of cross-reactivity are within the genera considering newly-discovered species remains unknown. To address these genus-wide mechanistic and antigenic questions, we assembled a curated set of HNV G and F sequences that samples the diversity of the genus as broadly as possible. We purified and characterized these proteins biochemically, biophysically, and structurally. By vaccinating mice with diverse HNV F proteins we elicited and isolated new antibodies with broad reactivity within the *Parahenipavirus* genus. We identified species with F proteins that demonstrate heightened pre-fusion stability; by translating structural features from these proteins, we stabilized the pre-fusion conformation of F proteins across the HNV genera. For the G protein, we characterized the binding of a G head domain panel to Ephrin B2 and B3, the known receptors for NiV and HeV, and to a panel of known anti-NiV antibodies. Additionally, we visualized a novel domain of a *Parahenivpavirus* G protein. Taken together, our findings in this study begin to distinguish antigenic groupings within recently discovered HNVs, clarify receptor binding differences in newly discovered species and antigenic relationships within the HNV genera, visualize new structural features in the F and G proteins, and provide insights for translateable pre-fusion stabilization techniques.

### Identification and Classification of HNV Species

To begin categorizing this diverse set of HNV strains, we identified as many unique HNV G and F ectodomains as could be found and assessed the phylogeny of these sequences. We used NCBI BLAST ^21^ to search for unique HNV-like sequences related to the input sequence, NC_002728 ^22^. We also carried out an extensive literature search to include other sequences that may not have been discoverable through BLAST, such as Gamak virus ^6^. Next, we narrowed the list to include only sequences from mostly complete genomes, leaving out those where only partial DNA sequences of G or F proteins were available. Almost all the strains that were selected had genome lengths close to the canonical 18.2 kbp for HNVs ^23^, with some of the species selected having slightly longer genomes reported. No sequence in our panel had a deposited genome shorter than 11.2 kbp. The several sequences that were shorter than the rest were either Nipah-Malaysia or Hendra strains that we included since they have been well-characterized in the literature.

Next, Clustal Omega ^24^ was used to align the G and F amino acid sequences. To generate the soluble ectodomain constructs, we truncated the sequences, using trends in the sequence alignments to decide on exact truncation points between species. All HNV-F ectodomains were of similar length, allowing all to be truncated at the C-terminal end of the ectodomain at the site equivalent to Nipah virus residue 488, consistent with previous F ectodomain constructs ^19^ **(Supplemental Data 1, Supplemental Table 1)**. For the G proteins, the domain boundaries did not align as consistently across species as they do with the overall less variable F protein, though the N-terminal portion of the sequence up to the end of the transmembrane domain was relatively consistent across all species, allowing for the truncation position for the ectodomain to be set at the analogous site to Nipah virus residue 71. While the full G ectodomain and its elusive fusion promotion mechanism is an important area for research, in this study we focus on the G head domain that harbors the receptor binding site and is targeted by a myriad of neutralizing antibodies ^14,16,25,26^. Therefore, while our sequence and phylogenetic analysis considers the whole genome or full G amino acid sequence, only the head domains of G proteins were purified. For these head domains, sequence variability resulted in the transition between the neck domain and head domain being less clear from sequence (**Supplemental Data 2)**. In classical *Henipavirus* sequences, there are frequent glycine and proline residues leading up to a conserved cysteine in the head domain. The Parahenipaviruses also contain a proline-rich region before the head domain, with as many as five consecutive prolines in some sequences. This proline-rich region is often followed in Parahenipaviruses by many aspartic and glutamic acid residues. Ultimately, for each strain we sought to select the site analogous to Nipah virus residue 177 as a suitable start point based on sequence analysis and available structural data for Nipah and Langya G proteins ^14,27^ **(Supplemental Data 2, Supplemental Table 1)**. After removing redundant sequences where differences were noted only in the signal sequence, transmembrane domains, cytosolic domains, or other regions not included in the final F ectodomain or G head domain constructs, the panel was assembled into a phylogenetic tree based on complete genomes (**Figure 1**). From this list, a subset of unique F ectodomains and G head domains was purified that would broadly sample the diversity of species in the phylogenetic tree, resulting in 35 purified F ectodomains and 32 purified G head domains. A listing of the sequence source for each purified protein can be found in **Supplemental Table 1** and exact amino acid sequences in **Supplemental Data 1-2.**

**Figure 1:**
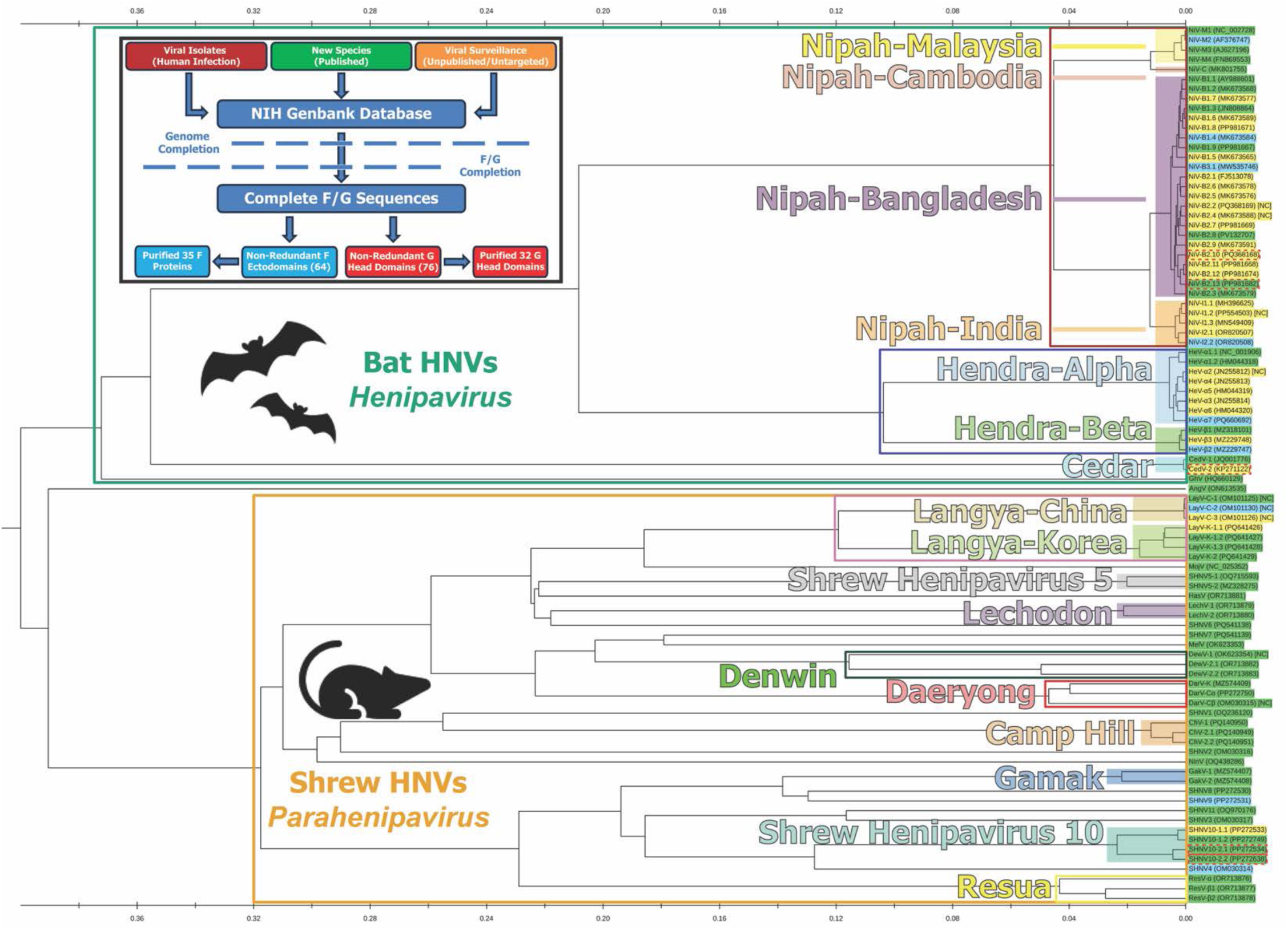
Selection and Classification of Henipavirus Species. **Upper Left:** A flowchart that describes the origin of sequences used and the filtering mechanisms, genome and F/G sequence completion, used to assemble a list of non-redundant F and G proteins. **Tree:** A phylogenetic tree based on full-genome nucleotide sequences of the selected strains. Horizontal branch lengths are to scale to represent sequence divergence. Hollow boxes are drawn to represent broad groupings within the tree such as genus, with Classical/Bat Henipaviruses (green) and Shrew/Parahenipaviruses (orange), as well as potential species groupings such as Nipah viruses, (dark red) and Hendra viruses (dark blue). Solid highlight boxes are drawn to represent more narrow groupings of specific species or subspecies groupings such as Nipah-Malaysia (yellow), Nipah-Bangladesh (purple), and Langya-China (tan). The tips of each branch are labelled with the strain names and associated gen bank accession number. Each tip is colored based on which unique glycoproteins are available from that strain: unique F ectodomain only, blue, unique G ectodomain only, yellow, no unique G head domain, dashed red border, both unique F and G ectodomain, green.

The International Committee on Taxonomy of Viruses (ICTV) has recently updated terminology related to the Henipaviruses, establishing the *Henipavirus* and *Parahenipavirus* genera, and specifying select species within these groups ^10^. Our effort to compare the properties of dozens of unique F and G sequences required finer granularity than the species level, and as a result, for this study we adopted an abbreviation and numbering scheme to aid in distinguishing specific sequences. The basis for this terminology is expanded upon in **Supplemental Note 1** and the corresponding accession code and current ICTV name for each sequence is listed in **Supplemental Table 1.**

For F proteins, the sequence identity relative to NiV-M1-F is high for both NiV and HeV strains, with all NiV sequences over 98% and HeV strains over 97% (**Supplemental Figure 1**). The sequence identity of most non-NiV/HeV strains to NiV-M1-F is ∼40%, with sequence similarity around 55-60%, revealing that the F protein is relatively well conserved across genera. A phylogenetic tree of HNV-F proteins based on amino acid sequences was overall similar to the whole genome tree with several small differences. Specifically, the distinction between NiV-C-F and NiV-M strains is more apparent in the F-based phylogenetic tree, the F proteins of SHNV3 and SHNV11 appear to be very closely related, and AngV-F appeared distinct from all other HNV strains, rather than part of the bat clade as seen through full-genome phylogeny (**Supplemental Figure 2, Top**).

Compared to F proteins, HNV-G proteins demonstrate much greater sequence variability. Sequence identity values are only ∼78% between NiV and HeV, and drop to below 20% for *Parahenipavirus* G proteins, compared to NiV-M1-G (**Supplemental Figure 1B**). The sequence similarity scores are also rather low, with most Parahenipavirus strains at around 35% similarity to NiV (**Supplemental Figure 1B**). When considering amino acid-based phylogeny, the relationships between strains remain similar as we had observed for the F ectodomains (**Supplemental Figure 2, Top**). One notable difference is that SHNV3-G and SHNV11-G, as with their F counterparts, appear more closely related than their full genome would suggest (**Supplemental Figure 2, Bottom**). Overall, the limited sequence similarity observed for the G head domains suggests that there could be greater antigenic, structural, and mechanistic differences between different HNV-G proteins than among F proteins.

### Preparation of Diverse HNV-F Ectodomains

We expressed 35 HNV-F ectodomains constructs that encoded residues 1 to the equivalent of 488 in NiV, followed by a foldon trimerization domain ^28^ and a C-terminal Twin Strep tag. For all constructs, the putative natural cleavage site that allows processing from the F_0_ precursor to the fusion-competent F_1_/F_2_ complex was retained. Following expression of the protein through transient transfection in HEK 293F cells and purification using their C-terminal Strep affinity tags followed by size exclusion chromatography (SEC), we obtained variable yields of F proteins, with some constructs expressing over 2 mg/L and others less than 0.1 mg/L (**Supplemental Figure 1**). For eight F proteins, the yields were too low to perform any downstream studies. We found that yields could vary dramatically between strains that only differed by a few amino acids. For example, NiV-B1.1-F and NiV-B1.2-F differ by only three residues in the ectodomain, but while NiV-B1.2-F produced ∼1 mg/L, NiV-B1.1-F had no yield across two purification attempts.

HNV-F ectodomains typically eluted from SEC with three peaks, one that eluted immediately after the column void volume, a broad “middle peak,” and the “main peak” that corresponds to the molecular weight of an F trimer eluting last (**Figure 2A, Supplemental Figure 3**), with the ratio of the peaks differing from strain to strain (**Supplemental Figure 1**). For some strains, a shoulder was observed for the middle peak, suggesting multiple sized components eluting within this peak. The SDS-PAGE profile of the middle peak fractions showed a major band at the molecular weight expected for the trimeric ectodomain unit, indicating that the presence of the higher molecular weight peak was not a result of any covalent linkage or differential glycosylation that would change the molecular weight enough to be distinguishable on SDS-PAGE (**Supplemental Figure 4**). This conclusion is supported by mass photometry analysis, where both the middle and main peaks, individually or mixed together, generated an essentially identical molecular weight profile, with each being dominated by the expected molecular weight of the trimer, and both showing a very small presence of species ∼2-3 times the mass of the trimer (**Figure 2A, Insert)**). SDS-PAGE also indicated that all F proteins are being expressed in their F_0_ state, without any proteolytic processing to F_1_/F_2_ **(Supplemental Figure 4).** Negative stain electron microscopy (NSEM) of each middle or main peak reveals comparable sets of two-dimensional class averages. The typical pre- and postfusion shapes identified previously ^19^ can be seen throughout the panel for both middle and main peak components (**Supplemental Figure 5**). Taken together, these data suggest that the middle peak is composed of transient, non-covalently associated F ectodomain trimers, which are reminiscent of previous studies of *Paramyxoviridae* F proteins where oligomeric states were reported for the F protein ^25,29,30^. In summary, though the F protein expression yields varied greatly, similar profiles were observed through SEC, SDS-PAGE, and NSEM analysis across all HNV species tested.

**Figure 2:**
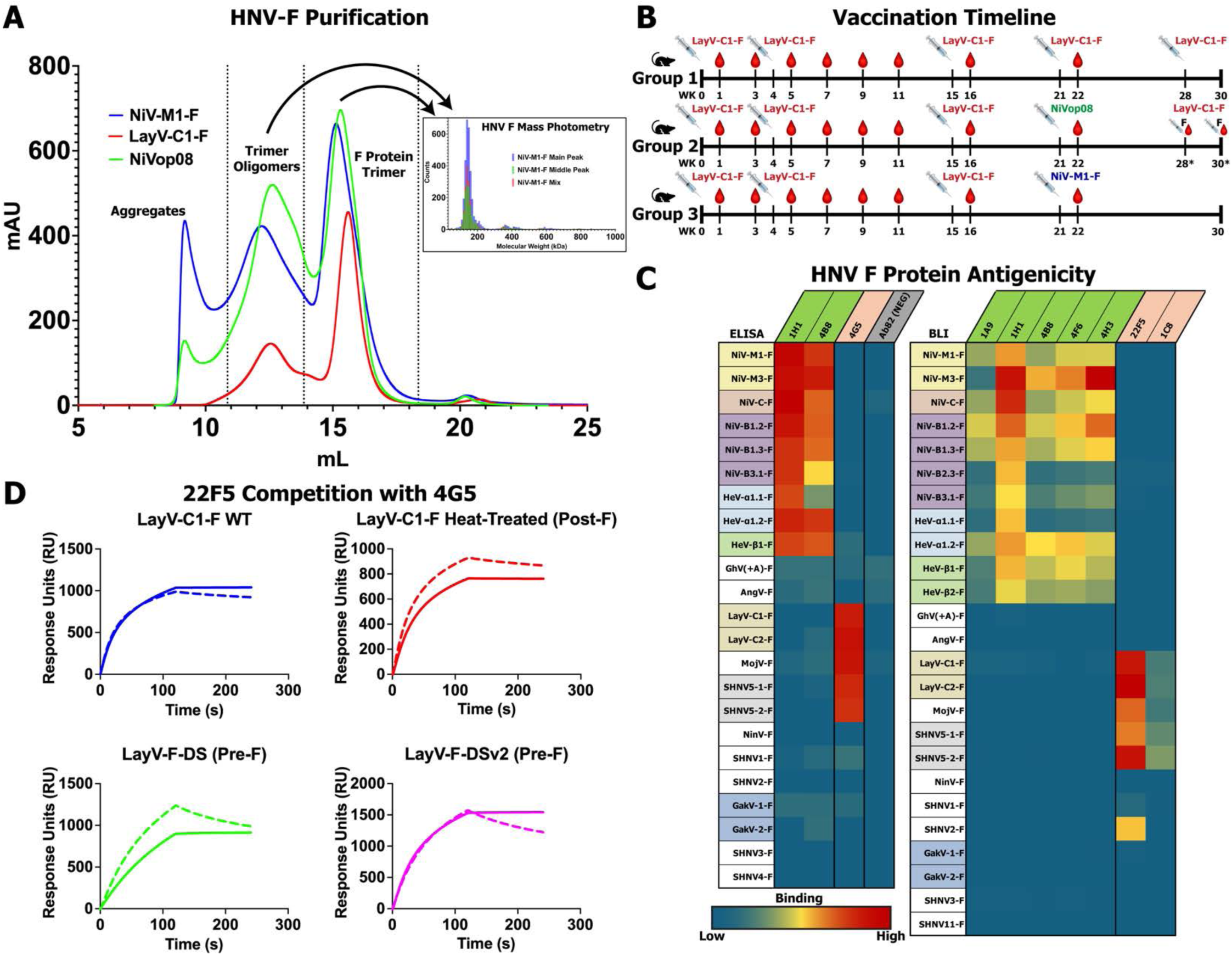
Purification and Antigenic Characterization of Henipavirus Fusion Proteins. **(A)** Representative SEC profiles of HNV-F Proteins with peaks annotated. The oligomer-of-trimer peak, sometimes called the “middle peak,” and the main trimer peaks are collected separately. NiVop08 is a pre-fusion stabilized construct ^31^ not part of the panel but purified for a vaccination study and for comparison to wild-type proteins. An inset to the right shows mass photometry analysis of NiV-M1-F SEC components. Each individual measurement is assigned a calculated molecular weight by the instrument software, and individual results are binned in 10 kDa bin sizes. The graph reports the count of events for each molecular weight bin with the main, middle, and mixed peak data overlayed. **(B)** Graphical timeline of a vaccination study in mice using HNV-F antigens. Each group was vaccinated with LayV-C1-F and boosted with LayV-C1-F or NiV antigens as marked. Blood draws and antigen injections occurred at the indicated weeks after the start of the study. The weeks with an asterisk, with the syringe and blood drop marked with F are weeks where a fusion was performed on an individual mouse. **(C)** Heatmap of antibody binding to HNV-F proteins, on a spectrum of blue for low binding to red for high binding. The left-side graph is based on ELISA data with immobilized F proteins, antibody analytes, and anti-Human Fab-conjugated HRP as a reporter. The values for each combination are based on the log of the area under the curve for the plotted absorbance over increasing concentrations. The right-side graph is based on Bio-layer interferometry (BLI) with immobilized F proteins and direct measurement of analyte to the BLI sensors, either mouse Fabs or mouse IgG (22F5 and 1C8). The values for each combination are based on the maximum wavelength shift measured during the association phase of the BLI experiment. HNV-F antigens colored as in Figure 1, antibody analytes colored as follows: green, known anti-NiV-F binding; salmon, known anti-LayV-F binding; gray, anti-influenza HA negative control antibody (ELISA only). **(D)** Surface plasmonic resonance (SPR) analysis of 22F5 and 4G5 antibody competition for binding of a series of LayV-F constructs in a variety of conformational states. Solid lines represent 22F5 alone and dashed lines with 4G5.

### Antigenicity of HNV-F Ectodomains

We next characterized the antigenicity of the F ectodomain panel. Since many new HNV species have been recently identified and given the limited availability of antibodies that react with LayV and other newly categorized members of our panel, we sought to identify antibodies of novel specificities by vaccinating mice with HNV-F antigens (**Figure 2B)**. One group was given four injections of the LayV-C1-F ectodomain, and for two others, the fourth injection of LayV-C1-F was replaced with NiV-F protein constructs, either the wild-type NiV-M1-F, or NiVop08, a pre-fusion stabilized NiV-F construct ^31^. By boosting with NiV-F constructs, we sought to elicit broadly reactive antibodies. Using hybridoma fusion technology ^32^, we identified two murine IgG1 antibodies, named 1C8 and 22F5, both from group 2, with reactivity to LayV-1-F. An additional vaccination series, using the same antigens with a similar timeline, led to the isolation of an additional four murine antibodies, all from group 2, named 9A9, 8C7, 20G7, and 8D4 **(Supplemental Figure 6)**.

The binding of antibodies elicited in our immunization experiments, as well as previously identified anti-NiV or anti-LayV antibodies, to our panel of HNV-F ectodomains was assessed by enzyme-linked immunosorbent assay (ELISA), surface plasmon resonance (SPR), or Bio-layer interferometry (BLI) ^20,33^ **(Figure 2C, Supplemental Figures 7-9)**. 1H1 and 4B8, previously identified anti-NiV-F antibodies, displayed broad reactivity to all NiV and HeV strains, but extremely limited reactivity to all other species. A panel of previously characterized mouse anti-NiV-F Fabs broadly reacted to nearly all NiV and HeV antigens, as expected. 4G5, previously identified as an anti-LayV-F and MojV-F antibody that binds both the pre- and post-fusion conformations at an epitope involving the HRA and HRB regions, strongly bound not only to LayV-F and MojV-F, but also to SHNV5-1-F and SHNV5-2-F, thus indicating that antibodies targeting the 4G5 epitope have greater breadth than was previously known (**Figure 2C, Supplemental Figure 7-8**).

Of the antibodies isolated from the immunization studies, 8C7, 20G7, and 9A9 bound only to NiV-F ectodomains. 1C8, 22F5, and 8D4 demonstrated strong reactivity not only to LayV-F proteins, but also to MojV-F and both SHNV5 strains. Moreover, 22F5, which generated stronger binding signals than 1C8, bound to SHNV2-F as well **(Figure 2C)**. 8D4 demonstrated the broadest binding profile, with binding detected to F proteins from each of GakV-1, GakV-2, NinV, SHNV1, SHNV2, and SHNV3 **(Supplemental Figure 9C).** Considering the phylogenetic relationships of these strains **(Figure 1)**, it appears that 8D4’s reactivity covers nearly the entirety of the known Parahenipaviruses. However, 8D4 demonstrated sharply reduced binding to pre-fusion stabilized F constructs, indicating that it specifically bound to the post-fusion F conformation **(Supplemental Figure 9C).** Since there are very few well-characterized antibodies that have been described for Parahenipaviruses, we selected 22F5 for detailed characterization, given its combination of a broad cross-reactivity profile and ability to bind pre-fusion F proteins.

To assess the potential to target the HNV-F protein glycans, we tested the binding of 2G12, a glycan reactive antibody previously noted for its broad specificity to diverse viral fusion proteins and ability to neutralize influenza and HIV-1^34–36^. We did not detect any binding between 2G12 and the HNV-F panel **(Supplemental Figure 9C)**. We also tested the reactivity of recombinant Griffithsin, a lectin discovered in *Griffithsia* red algae that has been investigated for antiviral properties ^37,38^ and has been shown to be protective against NiV challenge in hamsters^39^. Griffithsin bound to most of the NiV/HeV strains in the panel, while also binding to two pre-fusion stabilized *Parahenipavirus* constructs, of SHN5-1 and GakV-1, specifically **(Supplemental Figure 9C)**, underscoring the potential for a glycan-directed approach for broad spectrum targeting of HNVs.

Using (SPR), we tested the binding of 22F5 to LayV-F ectodomains in various conformational states, using two separate disulfide stabilized pre-fusion LayV-F constructs described previously, that we refer to here as DS ^30^ and DSv2 ^20^. Additionally, we tested binding with and without competition with 4G5 **(Figure 2D)**. 22F5 bound to LayV-F in both pre- and post-fusion conformations, indicating conformational bispecificity, and its binding was mostly unaffected by the presence of 4G5, indicating a distinct epitope. Given the strong and diverse binding profile of 22F5, we selected it for further characterization, including sequencing and structure determination **(Supplemental Figure 10-12)**.

Taken together, our findings align with the current understanding that Henipaviruses and Parahenipaviruses have distinct antigenic profiles. Parahenipavirus F proteins are not bound by known anti-NiV/HeV antibodies. However, the breadth of strains bound by our LayV-F-reactive antibodies indicates that cross-neutralization of a sizeable portion of known Parahenipaviruses with a small number of antibodies may be an achievable goal.

### Structural Characterization of 22F5 Antibody

Using single particle cryo-EM, we determined a structure of the 22F5 Fab bound to the post-fusion form of the LayV-C1-F protein, at a global resolution of 4.5 Å **(Figure 3A, Supplemental Figure 11, Table S2)**, with local resolution of ∼3Å at the interface of 22F5 with the F protein. 22F5 bound to the DII region ^13^ of LayV-F, with a few DI contacts as well. While there have been several previously observed antibodies targeting DII epitopes in NiV-F and HeV-F proteins ^33^, the only other available structure for an anti-LayV-F antibody, 4G5, details an epitope that includes contributions from HRA and HRB ^20^. Additionally, we determined the structure of the 22F5 Fab bound to the LayV-F-DS ectodomain at ∼4Å local resolution at the interface (**Figure 3B Supplemental Figure 12, Table S2**), confirming that the 22F5 epitope is accessible in both the pre- and post-fusion conformation of LayV-F. For both LayV-C1-F and LayV-F-DS, 22F5 was seen bound at all three DII sites.

**Figure 3:**
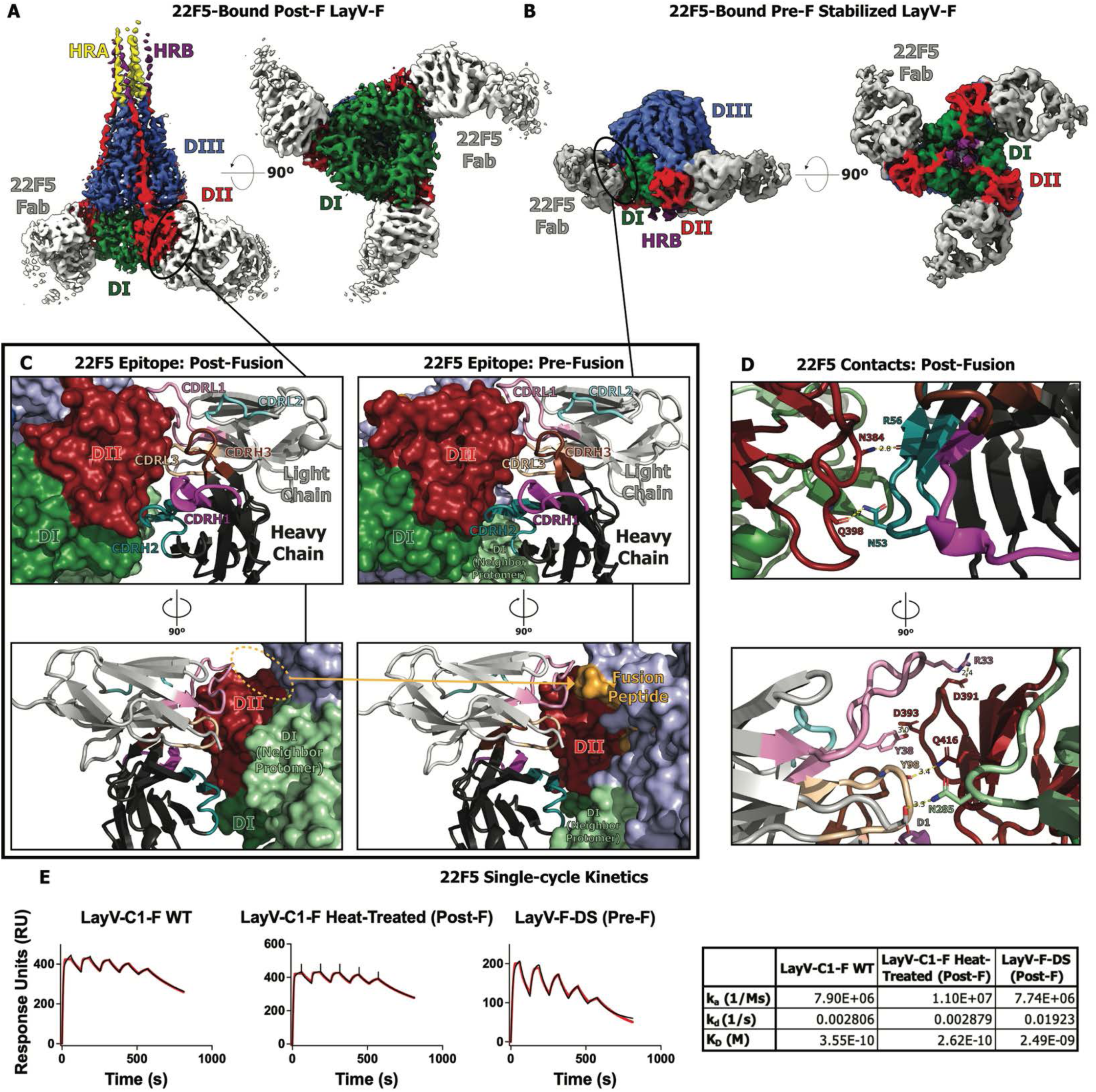
Structure and Binding Kinetics of 22F5 Fab with LayV-F Ectodomain. **A-B.** Cryo-EM maps of 22F5 Fab bound to post-fusion LayV-1-F **(A)** or pre-fusion LayV-F-DS (**B)** with domains colored and labelled. **C.** Views of the models (Left: Post-Fusion, Right: Pre-Fusion) built from the Cryo-EM maps in cartoon representation and colored by domain or CDR. **D.** Polar contacts between 22F5 Fab and the post-fusion LayV-1-F. Interacting residues labeled and shown in stick representation, with the color scheme continued from C**. E.** Measurement of affinity and kinetics of binding of 22F5 Fab to LayV-F ectodomains. The black lines in the graphs are the blank subtracted sensorgrams and the red lines are the fit of the data to a 1:1 Langmuir binding model. The table to the right shows the affinity and kinetic parameters obtained from the measurements.

The 22F5 Fab interacts with both the pre- and post-fusion LayV-F ectodomains utilizing both heavy and light chains. The L1, L3, and H2 complementarity-determining regions (CDR) are particularly involved **(Figure 3C, Supplemental Figure 13)**. In the pre-fusion conformation of LayV-F, a stretch of residues within DIII known as the fusion peptide is inserted into an interprotomer pocket formed by a neighboring protomer’s DII ^19^. During pre-to-post-fusion conformational conversion, the fusion peptide leaves this pocket. The pre-fusion 22F5 epitope includes surface-accessible portions of the fusion peptide, but in the post-fusion conformation, these residues no longer contribute to the 22F5 epitope **(Figure 3C, bottom)**. 22F5 binding appears to rely heavily on polar interactions **(Figure 3D)**. Specifically, heavy chain residues N54 and R57 contact LayV-F DII residues N384 and the main chain of Q398, respectively. In the light chain, R27, Y32, and Y92 interact with DII residues D391, D393, and Q416, respectively. Additionally, the 22F5 light chain N terminal residue D1 interacts with N285, a residue that belongs to domain DI of the neighboring protomer. When bound to the pre-fusion conformation, 22F5 contacts the neighboring protomer DI via its CDRH2 and framework region residues. Due to the shifting of protomers relative to each other during pre- to post-fusion conformational conversion, 22F5 shifts its interaction with the neighboring protomer DI region, losing its CDRH2 and heavy chain framework region contacts and gaining the light chain D1 contact with the post-fusion F protein. **(Figure 3C and 3D)**. Using SPR, we determined the affinity of 22F5 for both ectodomain constructs, as well as a heat-treated sample of the wild-type LayV-C1-F that ensured uniform post-fusion conformation, with binding observed at low nM affinities in all cases **(Figure 3E)**. Interestingly, despite the loss of contacts with the fusion peptide residues when bound to the post-fusion conformation of LayV-F, 22F5 bound tighter to the post-fusion conformation than the pre-fusion conformation. Given that the fusion peptide is a flexible region, it is possible that contact with it is entropically unfavorable, thereby explaining the improved affinity with the post-fusion LayV-F where this contact was absent.

In summary, 22F5 binding and structural data describe an epitope that is accessible in both pre-and post-fusion conformations and reveal a region on the *Parahenipavirus* F protein that can be targeted for greater breadth than previously recognized. 22F5 is a valuable addition to the antibody toolbox for the *Parahenipavirus* genus for which relatively few antibodies are known.

### Differential Scanning Fluorimetry of HNV F ectodomains

We assessed the stability of the purified F ectodomains using Differential Scanning Fluorimetry (DSF), a label-free method that measures changes in intrinsic fluorescence as the protein unfolds. The DSF profiles of the two wild-type proteins, NiV-M1-F and LayV-C1-F, and one prefusion-stabilized NiV-F construct, NiVop08 ^31^, were consistent with our previous reports^19^. With each peak in a DSF profile indicating a conformational transition, for the unstabilized F ectodomains we observed one transition between 45-65°C and one or more transitions at higher temperatures, up to over 90°C (**Figure 4A**). LayV-C1-F showed two transitions with a negative peak at 50°C and a positive peak at 86°C. NiV-M1-F showed three transitions with a negative peak 56°C, and positive peaks at 76°C, and 92°C. NiVop08 showed two transitions, a negative peak at 66°C and a positive peak at 76°C. We next measured the DSF profiles of the same set of proteins after pre-incubation at 60°C for 90 minutes, based on previous observations that heat treatment could force pre- to post-fusion conformational conversion in *Paramyxoviridae* F proteins ^40^. Overlaying all DSF profiles revealed that the lower-temperature negative peak present in all three proteins prior to heat treatment was eliminated in only the two wild-type proteins after heat incubation (**Figure 4A**). NSEM analysis confirmed that the wild type proteins have been converted to the postfusion conformation, as revealed by the characteristic elongated six-helix bundle present in the 2D class averages (**Figure 4B**). The NiVop08 prefusion stabilized construct retained the more compact shape characteristic of the prefusion state (**Figure 4B**). By exposing NiV-M1-F and LayV-C1-F to a series of times and temperatures, we found that 50°C, 15 minutes was the minimum amount of heat treatment sufficient to eliminate the pre-to-post-fusion conversion DSF peak, indicating that the F protein population was predominantly in the post-fusion state (**Supplemental Figure 14)**. Together, these results indicate that the negative DSF peak present around 45-60°C is indicative of a pre- to post-fusion conformational change taking place, and this DSF signature can be utilized to follow population level conformational transition from the pre- to post-fusion forms.

**Figure 4:**
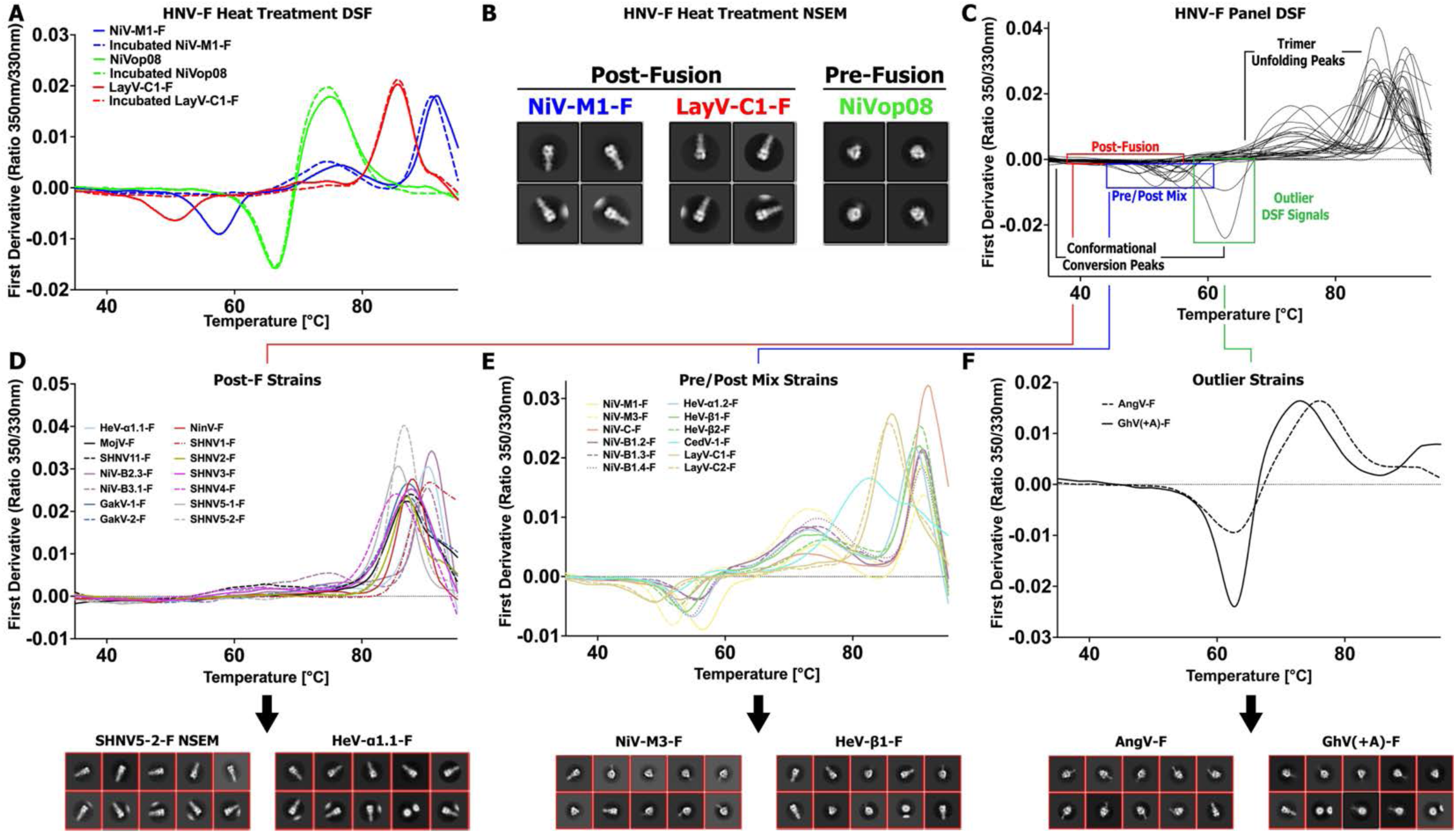
Thermostability of Henipavirus F Proteins. **(A)** DSF profiles of a set of HNV proteins showing the first derivative of 350/330nm emission ratio as a function of temperature. Dotted lines of the same color represent the DSF profile of the same proteins after being incubated at 60°C for 90 minutes. **(B)** Two-dimensional NSEM class averages of a portion of the heat-treated samples from A. **(C)** DSF profiles of diverse HNV-F species. Sections of the profile are noted, indicating the conformational steps occurring at those temperatures, with highlighting of peaks of interest. **(D)** A subset of DSF profiles from C, containing only those species that do not demonstrate any noticeable conformational conversion peak. Below are representative two-dimensional NSEM class averages for select members of this subset. **(E)** A subset of DSF profiles from C, containing only those species that demonstrate some presence of the standard conformational conversion signal. Below are representative two-dimensional NSEM class averages for select members of this subset. **(F)** A subset of DSF profiles from C, containing only those species that demonstrate a profile notably different from the typical HNV-F species. Below are representative two-dimensional NSEM class averages for the two members of this subset.

Next, measuring DSF profiles for all purified F proteins for which there was sufficient material purified, typically at least 5µg, we observed a wide range of conversion peak intensities and temperatures (**Figure 4C-F**). While all proteins generated a large positive peak at high temperature indicative of a denaturation event, not all proteins produced a detectable conversion peak. 14 proteins had essentially no detectable signal prior to the high temperature peak (**Figure 4D**). For these F ectodomain samples, two-dimensional class averages from NSEM analysis revealed the predominance of postfusion conformation for these proteins with no detectable pre-fusion F species observed in the datasets, supporting the DSF results. Though not exclusively, most of the proteins with no conversion peak are Parahenipaviruses, and inversely, all of the Parahenipaviruses have no conversion peak, except for Langya virus strains. Many of the Parahenipavirus F proteins could not be assessed due to insufficient purification yield, and these results may suggest it could be due to inherent instability. Among the F proteins that generated a conversion peak, there were shifts in temperature and intensity (**Figure 4E**). These changes did not appear related to species differences, with NiV-M1-F and NiV-M3-F, separated only by one amino acid mutation, being right- and left-shifted compared to the average, respectively. Both Langya virus strains were the most left-shifted of the set, indicative of lower pre-prefusion F stability. NSEM analysis showed a mix of pre- and postfusion conformations in this set, therefore the presence of this negative DSF peak indicates some amount of prefusion conformation present, rather than the presence of a uniform set of prefusion proteins. Between the conversion peak and the final high temperature peak is typically a broad, positive peak, present with differing intensities and shapes. Further investigation would be required to determine if this peak is either part of a natural, yet undefined, conformational event or part of the denaturation process.

Two proteins of the panel, AngV-F and the sequence-corrected version of GhV-F, GhV(+A)-F (**Supplemental Note 1**), display DSF profiles that are outliers to the rest of the panel (**Figure 4F**). Interestingly, both proteins displayed a profile that seemed closer to NiVop08, with their high temperature positive peak significantly left-shifted. Two-dimensional NSEM classes for these two proteins showed the presence of largely, but not exclusively, pre-fusion conformations. Despite their apparent similarities to a pre-fusion stabilized F construct, these are wild-type proteins that are expected to undergo normal conformational conversion, barring unexpected and dramatic mechanistic differences. The ability of the AngV-F and GhV(+A)-F proteins to undergo conformational conversion from the pre- to post-fusion states is supported by presence of post-fusion F forms in the NSEM 2D classes (**Supplemental Figure 5**). We incubated these outlier F proteins under both the 60°C, 90 minute and 50°C, 15 minute conditions. After incubation at 50°C for 15 minutes, a condition under which NiV-F and LayV-F proteins show effectively complete transition to the post-fusion conformation, we found that AngV-F and GhV(+A)-F proteins retain much of their native DSF profile, suggesting greater pre-fusion stability. Incubation at 60°C for 90 minutes, however, was sufficient to induce a dramatic change in their DSF profiles, suggesting more complete conformational conversion.

Since DSF is reliant on primarily Tryptophan residues to generate signal, it is possible that differences in DSF profile could arise from differing number and position of Tryptophans; however, the F ectodomain has only one Tryptophan that is almost universally conserved, with only CedV-1-F mutated at this spot (**Supplemental Figure 15**). Furthermore, AngV-F contains one additional Tryptophan that is unique to AngV-F. All F ectodomain constructs additionally contain one Tryptophan in the foldon tag used to assist with trimerization and two in the twin-strep tag used for purification. In general, sequence variability is likely not a significant factor in the differences observed in the DSF profiles of HNV-F proteins. Our results support the utilization of DSF as a rapid analytical technique requiring a small amount of protein for assessing population-level pre-to-post fusion transition of HNV-F proteins. In this study, the DSF analysis was central to our identification of two species that exhibit F pre-fusion conformations that appear more stabilized than in other HNV species.

### Cryo-EM Structures of AngV-F Protein

Based upon both amino acid sequence and pre-fusion stability as revealed by DSF analysis, AngV-F has divergent properties from the currently characterized F proteins. Therefore, we determined the structure of the AngV-F protein by cryo-EM single particle analysis (**Figure 5, Supplemental Figure 16-18, Supplemental Table 2**). The cryo-EM dataset revealed three particle populations (**Figure 5A**). The most populated class was of the isolated F ectodomain trimer, followed by a dimer-of-trimers population, and finally a population where the F proteins formed a hexameric array. The AngV-F trimer was refined to a global resolution of 4.0 Å and used for detailed examination of the F ectodomain structure and for comparing to the previously known structures of other F proteins. In cryo-EM structures of the HNV-F proteins the stalk region is typically poorly defined ^15,19,33,41,42^. Our AngV-F cryo-EM structure revealed well-defined density for the stalk, allowing atomic-level modeling.

**Figure 5.**
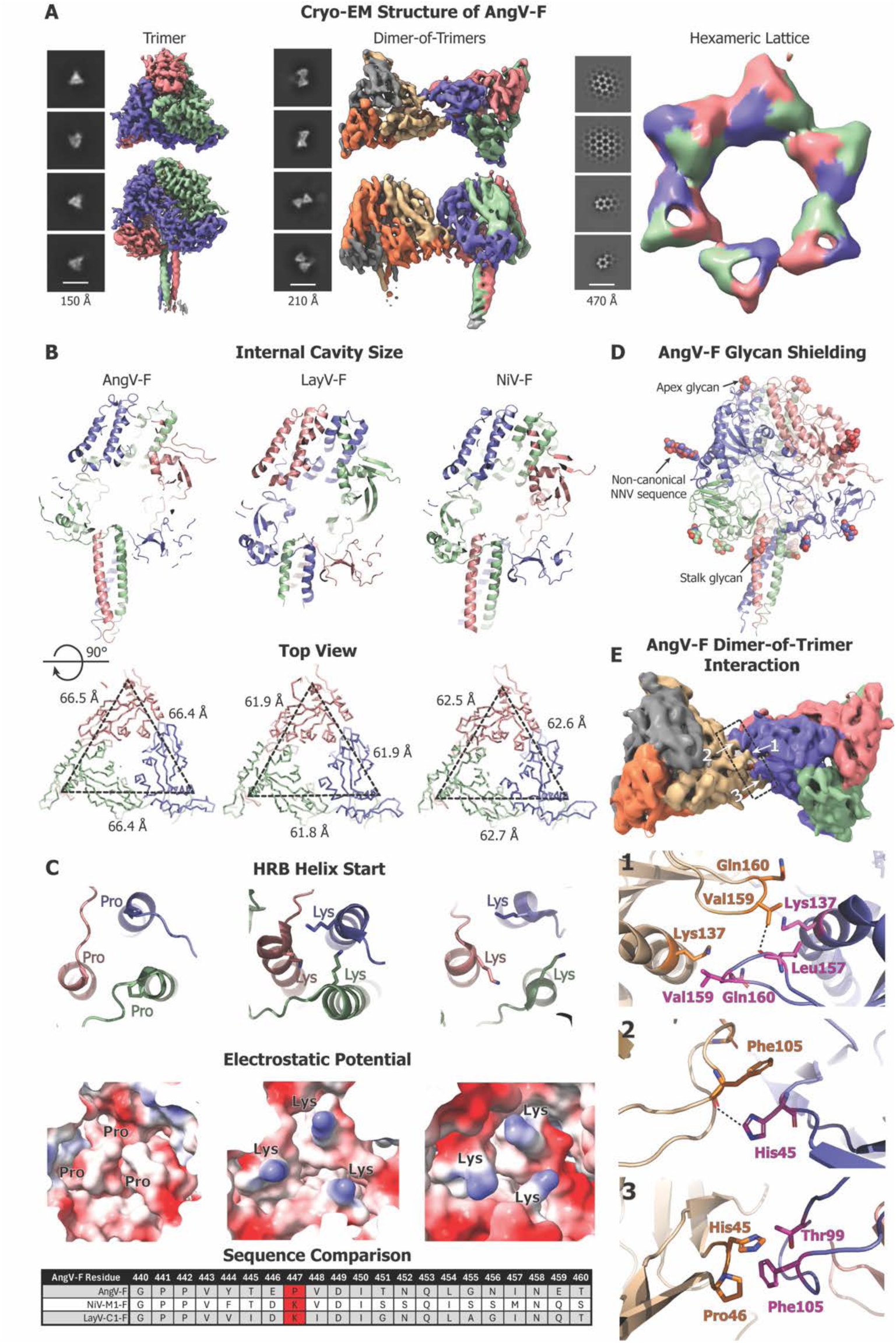
Cryo-EM Structure of the AngV-F Ectodomain. **(A)** Three different particle populations in the AngV-F and their representative 2D classes. Cryo-EM reconstruction of top and side view shown for AngV F trimer (left), dimer-of-trimers (middle) and hexameric lattice (right). **(B)** Comparison of the AngV-F, LayV-F and NiV-F proteins. Top. A slice through the F-protein visualizing the central cavity and the placement of the stalk region within the F-protein. Bottom. A top-down view showing measurements. **(C)** Zoom-in view of the stalk region featuring amino acid present at the start of helix bundle (top) and the electrostatics of the region (bottom), with sequence alignment at the local region between each of the three species below. The HRB start residue is marked red in the sequence alignment. **(D)** AngV-F cryo-EM structure colored by protomer with the glycans shown as spheres and colored by element. Glycans at the different regions (Apex, non-canonical NNV, stalk) are marked with arrows. **(E)** The dimer-of-trimer interaction interface is shown in surface representation (top), along with their molecular-level interactions and zoomed-in views showing details of the interface contacts (bottom).

The AngV-F ectodomain exhibited a larger central cavity compared to NiV-F and LayV-F proteins. This expanded cavity is the result of both the stalk region not inserting as far into the cavity as in other HNV-F proteins and the domains surrounding the cavity being spaced farther apart (**Figure 5B**). In previously determined HNV-F structures, the helical stalk region begins with a Lysine at its N-terminal end, thus positioning a trio of Lysines, one from each of the protomers, at the bottom of the internal cavity. In AngV-F, we instead observed a Proline at the beginning of the stalk (HRB) helix (**Figure 5C**). The surface electrostatics of this region are, therefore, strikingly different at this position for AngV-F compared to NiV-F and LayV-F structures. The proximity of the Lysines in LayV-F and NiV-F and their electrostatic repulsion could be a source of metastability in the pre-fusion F protein, as charge repulsion may be a mechanism to force apart the HRB helix during conformational conversion. The absence of this Lysine trio and replacement with a Proline cap at the N terminus of the HRB helix may lend greater stability to the pre-fusion form of the AngV-F protein. Notably, all species in our panel have either a Lysine or Arginine at this site, with only AngV-F substituting with a Proline (**Supplemental Data 1**).

Several glycans could be resolved (N61, N93, N353, N434 and N458), including ones analogous to those previously defined in HNV-F structures ^41,42^. The glycans shield known sites of vulnerability in *Paramyxoviridae* F proteins, including at the apex (N61), near the fusion peptide (N93 in Domain III), and on the stalk domain (N458). Interestingly, the glycan proximal to the fusion peptide utilizes a non-canonical NNV sequence (**Figure 5D**). The MojV-F and SHNV2-F proteins also have NNV sequences at this site that may be similarly glycosylated.

Glycans in the apex region near N61 and the HRB/stalk region near N458 are observed in many HNV-F protein structures (**Supplemental Figure 19**). However, there is no AngV-F analog to the N414 glycan of HeV-F and NiV-F. Instead, AngV-F contains two novel glycans in DI (N434) and DII (N353). The non-canonical NNV glycan at N93 is also found at the same site in HeV-F and NiV-F using canonical sequons, but interestingly, not in LayV-F. The differential glycosylation patterns observed across diverse HNV-F species may play a role in determining the potential for cross-reactivity, but critically, the presence of the NNV glycan in AngV-F indicates that prediction of glycosylation through sequence alone may be difficult.

Interactions between the AngV-F trimers were mediated through intermolecular contacts along the DI and DIII domains. These include hydrophobic interactions in DIII between L157 and V159 on neighboring ectodomains, both positioned on a loop that is part of an important antigenic site ^13,33^ (**Figure 5E**). Furthermore, there are charge interactions involving H45 from DI and T99 and F105 of DIII (**Figure 5E)**. Similar oligomeric states between pre-fusion F protein molecules (trimer, dimer-of-trimers, trimer-of-trimers) were also found in the cryo-EM dataset of a Hendra virus F protein (HeV-α1.2-F) (**Supplemental Figure 18**). AngV-F dimers resemble “mirror-like” protein dimers, with each exterior interaction surface of a protomer available for dimerization. These dimers, in turn, interact with protomers from three other trimers, forming hexameric structures. Each trimer is consistently aligned within the plane of the hexameric assembly, leading to a horizontally level lattice, distinct from an NiV-F crystal structure in which the lattice was not within one plane ^41^.

Taken together, our cryo-EM structures revealed that, while maintaining the overall conserved HNV-F architecture as observed in previously characterized HNV-F proteins, key mutations imparted notably distinct pre-fusion characteristics on the AngV-F protein. They also revealed multimerized pre-fusion F proteins, adding to the growing body of literature that suggests a role for the multimeric assemblies in the context of the virion and suggests a distinct mode of F protein multimerization leading to the formation of hexameric lattices for the AngV-F protein.

### Structure-guided Pre-fusion Stablization of diverse HNV-F Proteins

We hypothesized that the proline substitution observed at the start of the HRB helix, unique to AngV-F, was the cause of the increased pre-fusion stability. We expressed a series of HNV-F constructs from a variety of HNV species incorporating this proline substitution. Available HNV-F structures indicate that the site analogous to K453 in NiV-M1-F is consistently positioned at the N-terminal start of HRB, and even among other species for which structures are not available, the region surrounding this site is relatively well-conserved **(Figure 6A, Supplemental Data 1)**.

**Figure 6:**
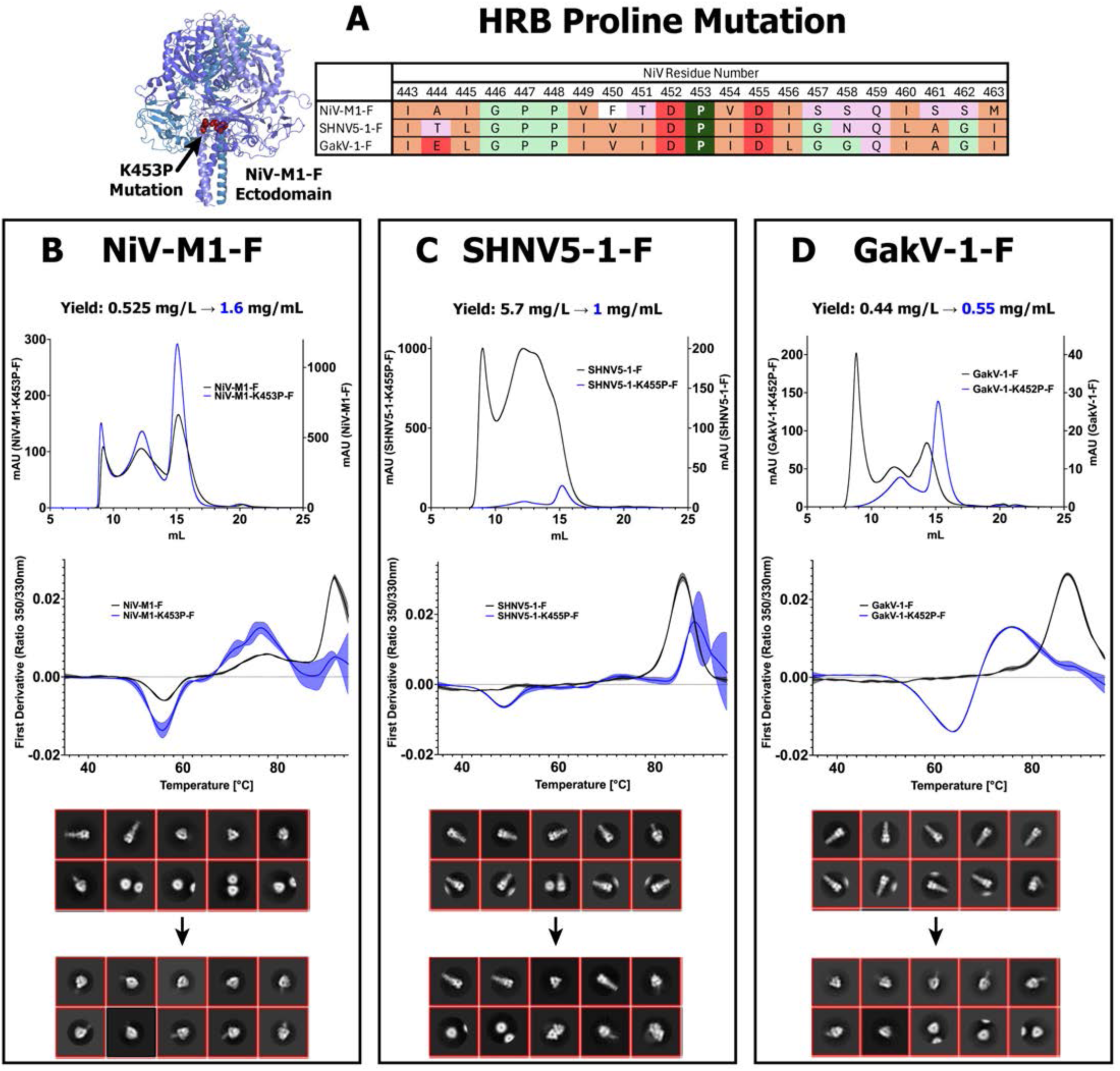
Translatable Pre-fusion Stabilization of Henipavirus F Proteins. **(A)** Location of HRB proline mutation in select HNV-F proteins. Left: Site of NiV K453P mutation shown as red spheres in the HNV-F ectodomain structure, with protomers colored in shades of blue. Right: A sequence alignment of the residues surrounding the HRB proline mutation, colored by residue type: hydrophobic, orange, polar, pink, negative charge, red, aromatic, white, conformationally significant, green, HRB proline mutation, dark green. **(B-D)** Characterization of the HRB proline mutation in three HNV-F species, **(B)** NiV-M1-F, **(C)** SHNV5-1-F, **(D)** GakV-1-F. Top: SEC overlay of the WT (right axis) and mutant (left axis) with mAU values scaled to a 1L transfection volume. Middle: DSF analysis overlay of the WT and mutant. The middle line represents the average with standard deviation represented with thin lines above and below the average line and shaded in-between. Bottom: Comparison of representative NSEM 2D classes from WT (top) and mutant (Bottom).

All of the proline mutants were able to be purified in sufficient amounts to permit downstream quantitation, with yields per transfection volume mostly exceeding those of the wild-type construct **(Figure 6B-D, Supplemental Figure 20A)**. In the case of the SHNV5-1-K455P mutant, the yield sharply decreased, but yet all mutants, including SHNV5-1-K445P-F, DSF analysis revealed significant pre-fusion stabilization **(Figure 6B-D, Supplemental Figure 20B)**. The trademark fusion conversion peak was present in all constructs, often with greater intensity than the wild-type, or demonstrating a right-shift of conversion temperature, possibly indicating higher stability of the pre-fusion conformation. In the case of the NiV, HeV, and LayV mutants, the 70-80°C peak was also present with higher intensity than the wild-type. Whereas GakV-1-F had displayed exclusively post-fusion conformation through both DSF and NSEM analysis **(Figure 6D),** the GakV-1-K452P-F DSF profile was dramatically altered, even appearing similar to the profile of the pre-fusion NiVop08 mutant or outliers like AngV-F and GhV(+A)-F. NSEM analysis confirmed the findings from DSF, with pre-fusion 2D classes observable with all mutants. The altered 2D classes of GakV-1-K452P-F, as with it’s DSF profile, was particularly striking. Whereas the wild-type classes contained solely post-fusion conformations, the mutant classes were almost universally pre-fusion. Despite the strong pre-fusion stabilization across all species, it was still common to observe post-fusion classes, reflecting that as with AngV-F, this mutation stabilizes pre-fusion conformation without preventing conversion **(Figure 6B-D, Supplemental Figure 20C)**.

We assessed the binding of the NiV and HeV-based mutants, NiV-M1-K458P-F and HeV-α1.2-K453P-F, to the pre-fusion specific 4B8 antibody. Strong binding was observed between 4B8 IgG and all F constructs tested: both wild-type and mutant NiV-M1-F and HeV-α1.2-F, and NiVop08 **(Supplemental Figure 20D)**. However, after incubation at 50°C for 15 minutes, while binding to NiVop08 was maintained, binding to wild-type NiV/HeV was nearly eliminated. The HeV proline mutatnt’s binding to 4B8 was similarly eliminated, while the NiV mutant’s binding was only moderately reduced. Taken together, our results indicate that pre-fusion stabilization of NiV/HeV through the substitution for the HRB start proline lends greater pre-fusion state stability but still allows for conformational conversion and antigenicity consistent with wild-type constructs.

We collected larger NSEM datasets and reconstructed 3D classes to quantify pre- and post-fusion ratios for these mutants and their wild-type counterparts. In all cases, the proportion of pre-fusion particles was increased in the mutant construct ccompared to wild-type **(Supplemental Figure 21A-F).** Interestingly, even for GakV-1-F and SHNV5-1-F, species where no pre-fusion conversion peak was present in DSF, a small amount of pre-fusion classes were detected in the larger NSEM datasets **(Supplemental Figure 21A-B, F)**. Most striking was the conversion of GakV-1-F from ∼10% pre-fusion classes, the least of any species, to over 75% pre-fusion with the proline mutant, the most of any species **(Supplemental Figure 21A-B)**. Although the use of NSEM results in low-resolution 3D volumes, it is still apparent that the typical pre- and post-fusion class shapes are observed in all species, indicating that the proline mutation has not significantly altered the pre-fusion conformation.

In summary, the introduction of this proline presents an effective and translateable means of pre-fusion stabilization for HNV-F proteins, and one that is expected to have minimal impact on any of the antigenic sites on the surface of the protein.

### Purification and Characterization of Diverse HNV-G Head Domains

To analyze our panel of HNV-G proteins, in this study we focused on the isolated head domain, which is responsible for receptor binding and is an important antigenic site ^13^. To allow for the previously described head domain sequences to be secreted in a mammalian expression system, we added an artificial secretion signal ^43^, and appended a C-terminal 8x His tag. The G head domain constructs were transiently transfected and affinity purified with cobalt resin affinity chromatography, followed by SEC. The typical SEC profile for the head domain proteins reflects the simple nature of their single-domain fold. All proteins eluted with a consistent single main peak (**Supplemental Figure 22**), typically with a small, higher molecular shoulder or peak likely consisting of aggregates. Yields for the head domains ranged from 2.5 to 35.3 mg/L (**Supplemental Figure 1**), with the majority yielding above 10 mg/L.

It has previously been noted that Langya and Mojiang virus attachment proteins do not bind to Ephrin B2 or B3 ligands (EFNB2/B3), the receptors used by Henipaviruses ^20,44^. Through BLI and SPR experiments, we assayed the binding of all G head domains in our panel to recombinant Fc-tagged Ephrin-B2 and -B3. As expected, all Nipah and Hendra virus head domains bound to both B2 and B3, with stronger binding observed for B2 rather than B3 **(Figure 7A, Supplemental Figure 23A-B)**. CedV-1-G and GhV-G also bound to B2, with very faint binding to B3. Consistent with the previous observations, Langya and Mojiang virus G head domains, as well as the rest of the panel, including all Parahenipaviruses, did not show any appreciable binding to either B2 or B3.

**Figure 7:**
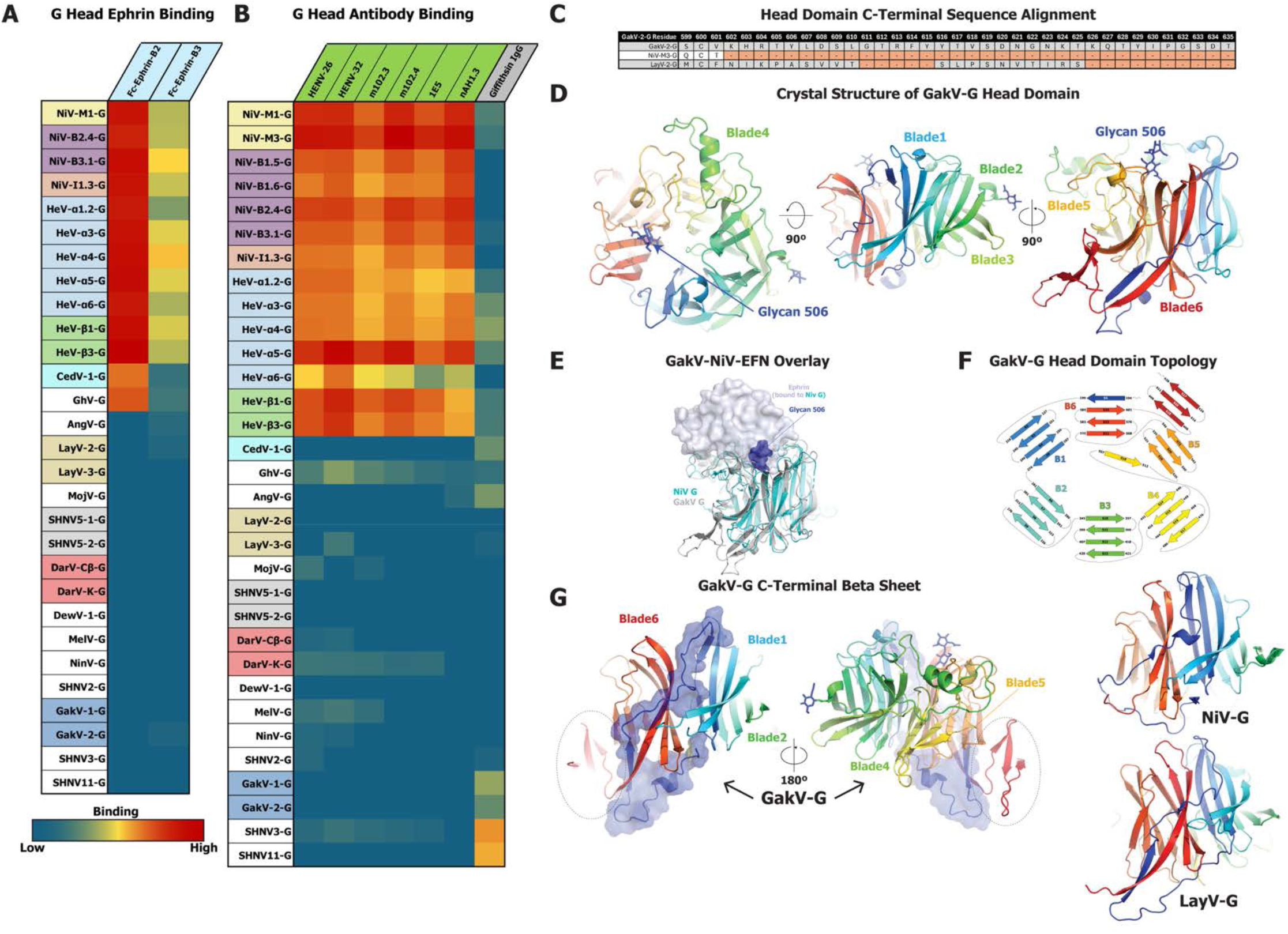
Structure and Characterization of Henipavirus G Proteins. **(A)** Heatmap of recombinant Fc-Ephrin-B2 and B3 binding to G head domains, on a spectrum of blue for low binding to red for high binding. The graph is based on BLI data with immobilized Ephrin and G head domain analytes. The value for each combination is based on the maximum response value from the association step. G head domain antigens in the table colored by group as in Figure 1. **(B)** Heatmap of antibody binding to G head domains, with color scheme identical to (A). The graph is based on BLI data as described in (A) using immobilized anti-G antibodies. **(C)** Sequence alignment of the C terminal end of select HNV-G proteins. **(D)** Three views of the GakV-G head domain colored in rainbow from N to C terminal. **(E)** Superposition of GakV-G head domain on Ephrin-bound NiV-G head domain (PDB: 2VSM ^51^). GakV-G is colored gray, NiV-G cyan and Ephrin light blue. Ephrin is shown as a transparent surface. The GakV-G glycan 506 is shown as blue spheres. **(F)** Topology diagram of GakV-G protein with blades of the beta propellor colored in rainbow from blade 1 through 6. **(G)** Left: Two views of the GakV-G head domain colored in rainbow with the N terminal Asp to His218 shown as a transparent surface. The additional minidomain defined in the GakV-2-G head structure is highlighted within a dashed oval. Right: G head domains of LayV and NiV.

We assessed the antigenicity of our G head domain panel with a series of previously characterized anti-NiV/HeV-G antibodies: HENV-26 and -32 ^16^, m102.3 and .4 ^25^, nAH1.3 ^45^ , and 1E5 ^26^. We also included the same Griffithsin construct used with the F panel.The known anti-NiV/HeV antibodies demonstrated broad, strong binding to all tested NiV and HeV strains **(Figure 7B, Supplemental Figure 23C-D)**. There were lower levels of binding observed for some of these antibodies outside of NiV/HeV strains, particularly with Ghana virus. Griffithsin showed moderate binding to several strains, albeit inconsistent and not seemingly linked to sequence similarity. It bound most tightly to the SHNV3-G and SHNV11-G head domains. Although it did not show consistent binding to NiV/HeV proteins, as with HNV-F, nevertheless, the ability to bind to several *Parahenipavirus* G proteins is notable, given the lack of known binders for these novel strains. Slight variations in binding were observed through ELISA, mostly in the form of low to moderate levels of 1E5 binding to GhV-G, DarV-K-G, and MelV-G **(Supplemental Figure 23D)**.

We used DSF to assess the thermostability of the G head domains. Unlike F proteins, the head domains are not known to undergo a dramatic conformational change when heated, and most proteins in the panel generated a profile with a single peak between 60-70°C, with some small shifts in inflection temperature that were largely consistent within species groupings. **(Supplemental Figure 24A-B)**. Several species generated profiles distinct from the most common result, such as broad, negative, or double peaks **(Supplemental Figure 24C-D)**. Unlike the relatively high conservation of Tryptophan positions in F proteins, G proteins display a much higher level of sequence diversity (**Supplemental Figure 15**). Indeed, there are substantial differences in Tryptophan positions in species that generated outlier profiles, CedV, GhV, AngV, and SHNV2 especially. The variability of Tryptophans in the G protein across species groupings precludes genus-wide comparisons of DSF profiles, yet these results provide interesting comparative insights within species groupings that may have implications for differences in receptor binding and virus fitness between species. As observed in the case of the SARS-CoV-2 spike protein, where thermostability of the receptor binding domain (RBD) is a property that is optimized during virus evolution ^46–48^, stability of the G head domain may impact its ability to bind receptor and antibodies, thus impacting its biological function.

### Structure of Gamak Virus Attachment Protein Head Domain

HNV-G proteins are tetramers (dimer of dimers), with each monomer composed of a helical stalk domain, a neck domain and a 6-bladed, β-propeller head domain where the receptor-binding site is located ^13^. When comparing the sequence diversity of the HNV-G proteins, we noted that the C-terminal end of G head domains are a site of substantial variability **(Figure 7C, Supplemental Data 2)**. GakV-G has among the longest C-terminal end, over 30 residues longer than NiV-G. A crystal structure of the GakV-2-G protein head domain at 1.4 Å resolution (**Figure 7D, Supplemental Table 3)** showed the expected 6-bladed β-propeller architecture that is well-conserved among HNV-G proteins. A well-resolved glycan was observed in the region that is analogous to the Ephrin (EFN) receptor binding surface in the NiV-G head domain (**Figure 7E).** The presence of this glycan further illustrates the changed receptor specificity of the Parahenipaviruses, as it would likely cause a steric hindrance to Ephrin binding if present in NiV or HeV. The elongated C-terminal end was resolved in our structure, revealing a novel domain spanning residues 606 to 628 that formed a three-stranded β-sheet stacked against blade 5 of the β-propeller (**Figure 7D, F, Supplemental Figure 25**). This additional compact and ordered domain is distinct from the previously known structures of *Henipavirus* G protein head domains. Furthermore, the N-terminus of the head domain appeared well ordered and packed against the 6-blade β-propeller core (**Figure 7G**). Residues 194-199 threaded between blades 5 and 6, contributing a β-strand to blade 6 as it continued to lead into blade 1. This stretch of residues also contacted blades 2, 4, and 5, thus forming a connection between different secondary structures within the β-propeller. Taken together, the structure of the GakV G head domain illustrates the structural diversity that can be accommodated within the conserved β-propeller fold to possibly impact receptor tropism and antigenicity.

## Discussion

Over the past few years, there has been a rapid expansion of identified species and strains in the *Henipavirus* genus, ultimately leading to the separation of these species into the *Henipavirus* and *Parahenipavirus* genera ^10^. Given the history of zoonotic transmission with these viruses, characterization of these novel species is an important step for pandemic readiness. Here we characterize a diversified panel of HNV F and G proteins to construct a panel that reflects the global diversity of HNVs. Until now, many of the strains in our expanded panel of Henipaviruses were known by their sequences alone, their surface glycoproteins totally uncharacterized and their relationships to other Henipaviruses unknown. Focusing particularly on Parahenipavirsues, closely related to Henipaviruses such as NiV and HeV, we elucidate their genetic diversity, antigenic relationships, structural novelties and mechanistic features.

Purifying and characterizing the F protein ectodomains of these strains has revealed that despite low sequence identity, HNV-F proteins share many architectural and biophysical similarities, a trend we previously noted by comparing Langya virus to Nipah virus ^19^. One of the most notable similarities was the presence of some level of oligomerization between F trimers. This was apparent from the SEC profiles, but the presence of trimer multimers did not seem to correlate with other parameters, such as purification yield, metastability, or phylogeny. It appears instead that the formation of these complexes is transient and dependent on the concentration and environment of the F protein, as suggested by the absence or very small proportions of oligomers detected by mass photometry or NSEM where sample concentration is much lower than during size-exclusion chromatography and cryo-EM. The ubiquity of this type of interaction indicates that oligomerization of F proteins is likely a conserved trait within HNVs.

F protein oligomerization was most striking in the case of Angavokely virus, where the formation of a hexameric lattice of F protein trimers was observed in the cryo-EM datasets. The existence of a hexamer-of-trimers arrangement has been noted previously with NiV ^41^, though the biological relevance was limited by the use of crosslinking to stabilize the arrangement. Our structure demonstrates the ability of AngV-F to multimerize in this way without any such assistance and, critically, without the involvement of the transmembrane domain, viral envelope, cytoplasmic domain, or viral matrix protein. The hexameric lattice is formed purely through ectodomain interactions, validating the characterizations of inter-trimer interactions previously reported for NiV ^30^, yet distinct in the implicated residues, with the AngV-F interaction involving DI and DIII, while the NiV-F interaction is dominated by DII residues. Previous studies have defined the lattice of *Paramyxoviridae* and *Pneumoviridae* F proteins, with various arrangements being determined. In one case, hexameric lattices have been observed, as with a whole-virion tomography structure of parainfluenza virus 3 ^40^. In others, such as with measles virus and respiratory syncytial virus, alternate arrangements of fusion and attachment proteins have been observed ^29,49^. Our results with AngV-F suggest that HNV-F proteins may adopt a hexameric arrangement, though structural studies of pseudoviruses and whole virions would help elucidate the physiological relevance of this finding. Furthermore, it cannot be ruled out that the formation of a hexameric F lattice may comprise one stage of HNV-F functionality, as the arrangement could shift during some type of maturation event or during G-mediated triggering.

Our biophysical characterizations of HNV-F proteins give us a rapid and inexpensive tool with which to assess metastability without relying on time-consuming structure determination. Using this assay we were able to identify distinct signatures for pre-to-post fusion conversion and F protein denaturation, revealing how differences in sequence or structure could impact metastability. Though there are limitations in biological relevance working with soluble F_0_ ectodomains, this assay has proven reliable in quickly detecting altered pre-fusion stability in HNV-F proteins. It was through this DSF assay that we identified the strongly pre-fusion stabilized character of AngV-F, leading to our discovery of the HRB helix proline substitution as a translatable means for pre-fusion stabilization. Previous attempts to translate successful pre-fusion stabilization strategies for NiV-F to Parahenipaviruses have required structure-based evaluation of analogous sites to the disulfide, cavity-filling, or proline substitution strategies used previously ^20,30^. However, the region surrounding the start of the HRB helix as well as specifically the positively charged residue substituted for a proline are so well-conserved as to have allowed successful and efficacious transfer of the stabilizing mutant to each species attempted without further decision on positioning of the substitution. Additionally, while strategies like the triple mutation found in NiVop08 appear to fully lock the F protein in a pre-fusion conformation, the HRB start proline more moderately stabilizes without completely preventing conformational conversion. Though the incorporation of such a mutation would impart significant limitations on the biological relevance of any studies that evaluate conformational change in a mutated F construct, its utility is highlighted in situations where wild-type constructs do not produce any pre-fusion protein whatsoever, as we have seen with SHNV5-1-F and GakV-1-F.

DSF analysis of G head domains also reveals a diversity of thermostability patterns. Although G head domains are comprised of a single domain 6-stranded β-propeller fold and are not known to have any major internal conformational rearrangements as part of their receptor-binding mechanism, some studies have indicated that receptor binding by HNV-G proteins could effect subtle changes in protein flexibility that do not manifest as large-scale conformational events ^50^. The differences in thermostability could potentially correlate with mechanistic differences between strains; however, more structural data detailing the structural changes that take place after receptor binding in G proteins will be needed to confirm this. From our crystal structure of the Gamak virus head domain, we can observe other likely sources of mechanistic differences between species. Though receptors for the Parhanehipaviruses remain unkown, the presence of the glycan in the traditional Ephrin-binding suggests an altered mode of receptor engagement. While the glycan may not be solely responsible for the lack of Ephrin binding in GakV-G, it would likely hinder binding of a similar protein receptor in the EFNB2/3 site, suggesting that GakV either uses a smaller receptor or perhaps a different receptor binding site entirely. This would not be without precedent in Paramyxoviruses. For example, though EFNB2/3 engages the same face of the head domain in NiV and HeV that sialic acid does in other Paramyxoviruses ^51,52^, the measles virus receptor, CD150, binds at the side surface of the head domain, making major contacts along blades 5 and 6 ^53^.

The C-terminal diversity in the head domains, and especially the additional exterior blade revealed in the GakV-2-G structure, may be the source of mechanistic changes as well. Together with the altered arrangement of the N- and C-terminal ends of the head domain we observed, these features could suggest a different pattern for quaternary interactions in GakV-G ectodomain tetramers, different types of conformational changes within GakV-G after receptor binding, or may even serve as a possible binding site for the as of yet undiscovered receptor.

While several anti-NiV and anti-HeV neutralizing antibodies have been well-characterized, the antigenicity of novel HNV strains had not yet been explored. By assessing the binding of a panel of both previously identified anti-HNV antibodies and newly discovered antibodies elicited through vaccination, we can now better describe the antigenic landscape of HNV proteins. As expected, we saw little ability of anti-NiV/HeV antibodies to bind to non-NiV/HeV proteins. Most interestingly, we found clear antigenic overlap between the F proteins of LayV, MojV, SHNV5, and to a lesser extent, GakV. Both a previously reported anti-MojV-F antibody, 4G5, and our vaccine-elicited antibodies, 22F5 especially, bound to these strains. When considering our phylogenetic analysis, both by full genome and amino acid sequence, it can be seen this cross-reactivity spans distantly related strains. SHNV5 appears to be more distantly related to LayV than HeV is to NiV. GakV is even more distant phylogenetically, appearing to be roughly as distant from LayV as CedV is from NiV.

While the G and F proteins are not particularly heavily glycosylated relative to other viral glycoproteins, they ultimately do exhibit a varied glycosylation landscape, with glycans appearing in important antigenic sites in some species. While the lectin griffithsin only showed moderate levels of binding to most antigens tested here, it was able to bind to a very diverse set of HNV-G head domains. Most interestingly, its strongest binding was to the SHNV3 and SHNV11-G head domains, proteins for which there are currently no known antibodies or ligands that can bind them. As investigation of griffithsin for antiviral potency across several families continues, it may prove a useful tool for analysis of novel HNV proteins.

With a group of viruses exhibiting such great sequence diversity, one major concern in the event of a possible widespread outbreak of disease in humans is that emergent infective strains may escape immunity conferred from prior infection or immunization with other HNV strains. This phenomenon can be observed with yearly outbreaks of new influenza strains or with the progression of SARS-CoV-2 evolution, where new strains emerged and quickly dominated the pool of circulating virus as they evaded prior immunity ^54,55^. A key focus of efforts to address this possibility has been the design of immunogens that can elicit antibodies with broad coverage of the currently exisitng and future diversity of strains. Being able to draw antigenic boundaries within the *Henipavirus* and *Parahenipavirus* genera is a foundation for similar efforts, and here we see that while the prospect of cross-reactivity over the entire genus remains elusive, there is already demonstrated cross-reactivity among a large portion of the Parahenipaviruses. While it will be important to assess this observed cross-reactivity in neutralization assays and animal challenge models, there is currently limited knowledge of both the host receptors for Parahenipaviruses and of the proper cell lines or animal models to study infection. While some progress has been made in identifying permissive cell lines for Parahenipaviruses^44,56^, the rate of fusion is typically much lower than seen with more well-studied Nipah/Hendra models, indicating that such viral entry assays could be in need of further optimization. Ultimately, in the absence of robust experimental systems to study the virological properties and the breadth of protection from potential HNV infections, the discoveries reported here build a critical foundation for future exploration of *Parahenipavirus* antigenicity.

Ultimately, this broad characterization of HNV proteins provides a set of essential biophysical and biochemical data for many species and strains that, to this point, had only been known as a sequence in a database. As these novel species are studied in more depth, our analysis can be used to provide context for the comparison of other key aspects of *Henipavirus* biochemistry, structure, and immunology.

## Acknowledgements

Cryo-EM data were collected at the Duke Krios at the Duke University Shared Materials Instrumentation Facility (SMIF), a member of the North Carolina Research Triangle Nanotechnology Network (RTNN), which is supported by the National Science Foundation (award number ECCS-2025064) as part of the National Nanotechnology Coordinated Infrastructure (NNCI). We thank Dr. Nilakshee Bhattacharya for assistance with microscope alignments. This study utilized the computational resources offered by Duke Research Computing (http://rc.duke.edu; NIH 1S10OD018164-01) at Duke University. This research used resources of the National Synchrotron Light Source II, a U.S. Department of Energy (DOE) Office of Science User Facility operated for the DOE Office of Science by Brookhaven National Laboratory under Contract No. DE-SC0012704. Use of the NYX beamline 19-ID at the National Synchrotron Light Source II was supported by the New York Structural Biology Center.

NYX detector instrumentation was supported by grant S10OD030394 through the Office of the Director, National Institutes of Health. This work was supported by funding from the Translating Duke Health Initiative, Duke School of Medicine, and NIH R01 AI 165147 (P.A.). We thank Fred Alt, Ming Tian and Sai Luo for providing the V_H_1-2R^JH2^/Vκ1-33R^CSΔ/hTdT^ (SE13) mice, Hannah Violette and Nicholas Levering for technical assistance, and Nilakshee Bhattacharya for assistance with cryo-EM data collection.

## Author contributions

Conceptualization – A.J.M. and P.A.; Methodology –A.J.M., A.B., B.F.H. and P.A.; Investigation – A.J.M., M.L., J.J.L., A.B., M.D., M.B., R.P., X.L., M.B., A.N., X.H., A.M., U.K., K.S., V.I., S.S., C.S.P., R.D.A., P.D., K.J., Y.L., R.J.E., B.F.H., P.A. Data Curation – A.J.M., M.L., J.J.L, A.B. and P.A. Validation – A.M., M.L., J.J.L, A.B. and P.A..; Visualization – A.J.M., M.L., J.J.L, A.B. and P.A.; Writing - Original Draft – A.J.M., A.B. and P.A.; Writing - Review & Editing – all authors; Supervision – A.J.M., K.W., R.J.E, K.O.S., B.F.H. and P.A.; Project Administration –A.J.M. and P.A.; Funding acquisition – B.F.H. and P.A.

## Declaration of interest

A.J.M., M.L., J.J.L, A.B., B.F.H. and P.A. are named in provisional patents submitted on the methods for stabilizing the pre-fusion HNV F and G proteins based on the findings reported in this paper. Other authors declare no competing interests.

## Data and Material Availability

Cryo-EM reconstructions and atomic models generated during this study are available at wwPDB and EMBD (www.rcsb.org; http://emsearch.rutgers.edu) under the following accession codes: AngV-F trimer - EMD ID: EMD-48423 and PDB ID: 9MNH; AngV-F dimer-of-trimer - EMD ID: EMD-48535 and PDB ID: 9MQN. The crystal structure of the GakV G protein head domain was deposited with the PDB ID: 9EHU.

All materials will be made available upon request.

## Ethics Statement

All mice were cared for in a facility accredited by the Association for Assessment and Accreditation of Laboratory Animal Care International (AAALAC). All study protocols and all veterinarian procedures were approved by the Duke University Institutional Animal Care and Use Committee (IACUC).

Environment: All rooms are on a 12/12 light cycle unless otherwise requested. Heat and humidity are maintained within the parameters outlined in The Guide for the Care and Use of Laboratory Animals and animals are fed a standard rodent diet. Breeding or specialty diets will be provided upon request. Rodents and Guinea pigs are housed in individually ventilated micro-isolator caging on corn cob bedding. DLAR will provide alternative bedding based on scientific need. Mice are changed every two weeks, rats are changed twice a week, and Guinea pigs are changed three times a week or more frequently as needed. Group housing of up to 5 mice per cage is strongly encouraged. Environmental enrichment for singly housed animals will be provided unless an exception is granted by the IACUC.

## Methods

### Phylogenetic Analysis

Candidate sequences were identified by using the NCBI Protein BLAST tool ^66^. Nipah-Malaysia sequence NC_002728 was used as an input sequence. Sequences were aligned using the MAFFT algorithm ^67^. We removed any sequences for which there were gaps in the amino acid sequence, ensuring each entry contained a full ectodomain protein sequence. The final tree was built as a UPGMA based on the distance matrix using the Tree Viewer software ^68^.

### Protein Production

For HNV proteins, human codon-optimized DNA sequences were synthesized by GeneImmune Biotechnology and cloned into a pαH plasmid vector. Fusion protein ectodomain sequences (**Supplemental Data 1**) were followed at the C-terminal end by a foldon trimerization domain, HRV3C protease site, 8x his tag, and a twin-strep tag (IBA Lifesciences) Attachment protein head domains (**Supplemental Data 2**) had an artificial secretion signal, secrecon ^43^, added to the N-terminal end and an 8x his tag added to the C-terminal end. HNV protein plasmids were transiently transfected into HEK 293 cell lines, either Expi293 or 293F (Thermo Fischer Scientific) (**Supplemental Table 1**). The cells were allowed to incubate at 37°C, 8% CO_2_ and the supernatant was harvested 5 days post-transfection.

Fusion proteins were purified by first sequestering free biotin by applying BioLock blocking solution (IBA) and then using the Strep-Tactin affinity chromatography resin (IBA). Eluted protein was then further purified using a Superose 6 Increase 10/300 GL size-exclusion column (Cytiva) with PBS buffer at pH 8. The elution from SEC was spin concentrated with 100kDa filters and flash frozen with liquid nitrogen.

Attachment protein head domains were purified first by sequestering EDTA by adding 2mg of Cobalt Chloride per 1L of supernatant. TALON Cobalt Resin (Takara) was used for purification, with 20mM HEPES and 150mM NaCl at pH 8 as a buffer, adding 50mM Imidazole for elution. Proteins were further purified with a Superdex 75 Increase 10/300 GL SEC column (Cytiva), again using the HEPES-NaCl buffer. The elution from SEC was spin concentrated with 10kDa filters and flash frozen with liquid nitrogen.

Antibody human codon-optimized DNA sequences were synthesized by GeneImmune Biotechnology and cloned into pVRC8400 plasmid vectors. Heavy chains were designed to begin with the beginning of the antibody sequence and continue through the human IgG1 Fc region. Light chains included both the VL and CL domains. Depending on the sequence source, some antibody sequences either utilized mouse or human VL and VH domains. Mouse anti-NiV-F fabs were designed to end after the mouse CH1 domain and had an 8x his tag added to the C-terminal end, as described previously ^33^.

Plasmids encoding the heavy and light chains of the antibodies IgG 1E5, nAH1.3, 4G5, and DH851.3, or an Fc-tagged Griffithsin construct, were co-transfected into Expi293 cells at a 1:1 ratio, using the ExpiFectamine 293 Transfection Kit (Thermo Fisher Scientific, Cat. No. A14525), according to the manufacturer’s instructions. Following a six-day incubation period post-transfection, the cells were harvested, and the culture supernatant was clarified by filtration through a 0.2 µm PES membrane filter to remove cellular debris. IgG purification was subsequently carried out using Protein A affinity chromatography. The filtered supernatant was passed through the Protein A affinity column (Pierce^TM^ Recombinant Protein A Agarose Cat No. 20334), and the bound IgG was eluted using IgG elution buffer by adjusting pH via Tris Base.

Eluted fractions were pooled and concentrated with Centricon-70 30 kDA filter (Millipore Sigma) followed by size-exclusion chromatography using a HiLoad 16/600 Superdex 200pg column (Cytiva), pre-equilibrated with 1× phosphate-buffered saline (PBS, pH 7.4) containing 0.02% sodium azide to prevent microbial contamination. The purified IgG was collected, concentrated, and analyzed for purity and integrity by SDS-PAGE.

Anti-NiV-F Fabs were purified using the same Cobalt resin-based procedure as described above for head domains.

To generate Fab fragments for 22F5 mouse IgG and 1C8 mouse IgG, 2 mg of each IgGs were passed through Zeba Spin Desalting Columns (ThermoFisher), then washed with digestion buffer containing 35 mg cysteine. IgGs were incubated with 0.250 ml of the 50% slurry of

Immobilized Papain resin (ThermoFisher) followed by incubation with digestion buffer containing 35 mg cysteine at 37°C overnight in microcentrifuge tubes. The digested Fabs were separated from the Immobilized Papain by centrifuging at 5000 x g for 1 minute, followed by washing with digestion buffer. After digestion, the mixture was subjected to protein A affinity chromatography (ThermoFisher), then monitored by SDS-PAGE to confirm Fab generation. To further purify the Fab fragments, Size-Exclusion Chromatography (SEC) was used for purification of the Fab fragments using Superose 6 Increase 10/300 column GL (Cytiva). The sample was loaded onto the column at a volume of 0.5 mL and the elution was carried out with PBS at a flow rate of 0.5 mL/min. Fractions were collected, yielding a final concentration of 3.2 mg/mL and 4.2 mg/ml for 22F5 Fab and 1C8 Fab, respectively. The purified Fab was stored at −80°C.

The ELISA control antibody Ab82 was purified as described previously ^69^.

### DSF

Samples were analyzed using a Prometheus Panta (Nanotemper Technologies). All samples were diluted to 0.125mg/mL using PBS, pH 8. Roughly 10µL was added to glass capillaries in triplicate for measurements. The instrument was programmed to measure 350nm and 330nm fluorescence from 35.0-95.0°C using a 6.0°C/min temperature gradient. All curves reported represent averages from the triplicate readings.

### BLI

Bio-layer interferometry (BLI) was employed to assess the binding between HNV proteins and various antibodies. For fusion proteins (Figure 2), the analysis was performed using an Octet RH16 (Octet RED384) system (Sartorius) at 25°C with kinetics buffer (Sartorius) as the running buffer. HNV-F proteins with the twin-strep tag at 6 µg/mL were captured onto Streptavidin (SA) Biosensors (Sartorius) for 300s, followed by a 60s wash in kinetics buffer. The biosensors were then exposed to 20 µg/mL mAbs or 50 µg/mL Fabs for 300s to allow for association with the fusion protein. Dissociation in kinetics buffer was monitored for an additional 300s. Sensor data was analyzed using Octet Analysis Studio (Sartorius), with reference biosensor and zero-analyte control sensorgrams subtracted from the raw data to obtain the curves corresponding to specific binding events. For additional fusion protein experiments (Supplemental Figure 9), the analysis was performed with the same equipment, tips, and buffer. HNV-F proteins were loaded at 6µg/mL for 300s and equilibrated in buffer for 60s before a 120s association step and 200s dissociation step. All analytes were at 50µg/mL. Regeneration with 10mM Glycine, pH 1.7 was employed, using tips for 6 cycles and regenerating with 10s pulses between regeneration solution and buffer. SARS-S2-HexaPro ^70^ was loaded onto a tip as a reference sensor. Kinetics buffer was used as a reference sample. Double reference subtraction was performed, except for in the case of Griffithsin IgG, which bound to the SARS-S2-HexaPro, so only reference sample subtraction was used. For each cycle, NiVop08 was loaded and bound to 4B8 Fab, and the signal relative to the first use of the tip was used to adjust the response values of all association curves during that cycle. For attachment head domains with human Fc-containing Abs, the analysis was performed using an Octet RH16 (Octet RED384) system (Sartorius) at 30°C with HBS-EP+ Buffer (Cytiva) as the running buffer. Antibodies at 2 µg/mL were loaded onto anti-human antibody capture (AHC) biosensors (Sartorius) for 300s, followed by a 60s buffer baseline. Association with head domains at 25 µg/mL lasted 120s, followed by dissociation in buffer for 200s. Between cycles, biosensors were regenerated by alternating 3x in 10mM Glycine pH 1.0 buffer and running buffer for 5s each. Sensors were used for no more than 10 cycles of regeneration. Double reference subtraction was performed using both a control-loaded sensor and a zero-analyte reference sample.

### SPR

For 22F5-LayV-F interactions, binding and kinetic assays were performed on a T-200 Biacore system (Cytiva) at 25°C. All the binding and kinetic experiments were carried out in the HBS-EP+ (10 mM HEPES, pH 7.4, 150 mM NaCl, 3 mM EDTA and 0.05% surfactant P-20) running buffer. For the binding assay, an anti-Fc mouse SPR chip was used to capture 200nM of 22F5 IgG at 10µl/min flow rate and 50nM of LayV-F ectodomains as analyte were run at 30µl/min flow rate and association and dissociation were monitored at 120 seconds. In the competitive assay, 200nM of 4G5 IgG was captured on an anti-Fc human SPR chip at 10µl/min and analyte as 22F5 Fab with LayV-F ectodomain complex in 5:1 ratio was flowed at 30µl/min flow rate for 60 seconds. SPR SA chip was used to capture the 50nM of the LayV-F ectodomains at 5µl/min flow rate by biotin tag for kinetics analysis. 22F5 Fab 50nM with two-fold dilution (50nM, 25nM, 12.5nM, 6.25nM and 3.12nM) as analyte was flowed with 50µl/min flow rate and association and dissociation times were analyzed at 60 and 240 seconds, respectively. For the reference curves, we kept the same parameter in reference flow cells without protein capture. The sensorgrams were blank corrected and analyzed in the Biacore T-200 evaluation software.

For Ephrin-HNV-G interactions, Binding experiments were performed on a Biacore T-200 (Cytiva) with HBS buffer supplemented with 3 mM EDTA and 0.05% surfactant P-20 (HBS-EP+, Cytiva, MA). All binding assays were performed at 25°C. Ephrin binding to head domains was assessed using a Series S CM5 chip (Cytiva, MA), which was labeled with anti-human IgG (Fc) antibody using a Human Antibody Capture Kit (Cytiva, MA). Ephrin was coated onto the chip at 12.5 nM (120 s at 5 μL/min). Head domains were injected at 100 nM over the ephrin using a single injection (60 s on time, 180 s off time at 50 µL/min). The surface was regenerated with 3 pulses of a 3 M MgCl_2_ solution for 10 s at 100 μL/min.

### Negative-Stain Electron Microscopy

Frozen samples from -80 °C were thawed at room temperature for 5 minutes. Samples were then diluted to 40 µg/mL with 0.02 g/dL Ruthenium Red in HBS (20 mM HEPES, 150 mM NaCl pH 7.4) buffer containing 8 mM glutaraldehyde. After 5-minute incubation, glutaraldehyde was quenched by adding sufficient 1M Tris stock, pH 7.4, to give 80 mM final Tris concentration and incubated for 5 minutes. Quenched sample was applied to a glow-discharged carbon-coated EM grid for 10-12 seconds, blotted, consecutively rinsed with 2 drops of 1/20X HBS, and stained with 2 g/dL uranyl formate for 1 minute, blotted and air-dried. Grids were examined on a Philips EM420 electron microscope operating at 120 kV and a nominal magnification of 49,000x, and images were collected on a 76 Mpix CCD camera at 2.4 Å/pixel. Images were analyzed by 2D class averages using standard protocols with Relion 3.0 ^71^.

### Mass Photometry

A TwoMP (Refeyn) mass photometer was used for analysis. The AcquireMP software was used for calibration and data acquisition. Calibration was performed with a three-point curve consisting of β-amylase, Apoferritin, and Thyroglobulin. Samples were diluted to 40 nM in PBS. To each well, 10µL of PBS was added to focus the system before adding 10µL of sample and mixing for a final concentration of 20 nM. Recorded data for 60s, with the machine counting individual adsorption events. The DiscoverMP processing software (Refeyn) was used to assign molecular weights to individual events.

### Mouse Immunizations

Immunization in V_H_1-2R^JH2^/Vκ1-33R^CSΔ/hTdT^ (SE13) mice ^72^ were intramuscularly immunized with LayV-F_WT (25 mcg/animal) and NiVop8 (25 mcg/animal) adjuvanted with GLA-SE (IDRI EM-082, 5 mcg/animal) at weeks 0, 4, 15 and 21. Serum titers were monitored by ELISA as described below. Mice with high-binding antibody titers were selected for the subsequent spleen cell fusion experiments.

### Hybridoma cell line generation and monoclonal antibody production

Mice were boosted with the indicated priming antigen 3 days prior to fusion. Spleen cells were harvested and fused with NS0 murine myeloma cells using PEG1500 to generate hybridomas ^32^. After 2 weeks, supernatant of hybridoma clones were collected and screened by binding ELISA as described below. Hybridomas that secreted LayV-F reactive antibodies were cloned by limiting dilution until the phenotypes of all limiting dilution wells were identical. IgG mAbs were purified by protein G purification.

### Indirect binding ELISA method

ELISA assays were utilized for three purposes, measuring mouse serum antibody titers, screening for antigen specific hybridoma B cell clones, and characterizing the binding of monoclonal human IgG and mouse IgG. 384 well ELISA plates (Costar #3700) were coated with 2 µg/mL streptavidin (Thermo Fisher Scientific Inc. Cat. No. S-888) or protein antigen in 0.1M sodium bicarbonate overnight at 4°C. Plates were washed with PBS/0.1% Tween-20 and blocked for one hour with assay diluent (PBS containing 4%(w/v) whey protein/15% Normal Goat Serum/0.5% Tween-20/ 0.05% Sodium Azide). Streptavidin coated plates were washed and followed by 10 μL Twin-Strep-tag® (IBA Lifesciences GmbH) protein at 2 µg/mL in assay diluent for one hour. All plates were then washed and samples added in 10μl volumes with dilution scheme and secondary specifics depending on sample type as follows. Mouse serum was titrated from 1 to 30 in three-fold steps and incubated for 1 hour. Plates were washed and followed with 10µL goat anti-mouse IgG-HRP secondary antibody diluted 1:10,000 (Southern Biotech #1030-05). Hybridoma supernatants were added undiluted as a single well for 1 hour, washed and followed with 10 µL goat anti-mouse IgG-HRP. Monoclonal mouse IgG and Human IgG were diluted to 100 µg/mL, titrated in three-fold steps and incubated for 1 hour. Plates were washed and followed with 10 µL goat anti-Mouse IgG Fab-HRP diluted 1:10,000 (Southern Biotech 1015-05) for mouse IgGs or 10 µL of goat anti-human IgG Fab-HRP diluted 1:15,000 (Jackson Immunoresearch, 109-035-097 ) for human IgG monoclonal mAbs. After 1 hour, plates were washed again and detected with 20 µL SureBlue Reserve (Seracare 5120-0081) for 15 minutes. Reactions were stopped with the addition of 20 µL HCl stop solution. Plates were read at 450 nm using a Spectramax 384 plus.

### Antibody gene cloning and sequencing

Cell suspensions of the 22F5 hybridoma cell line were pelleted by centrifugation at 300 g for 5 minutes. The cell pellets were washed twice with Phosphate-buffered saline (PBS, 1X) to remove residual culture medium. The 22F5 antibody genes were amplified from the hybridoma cells by RT-PCR. Briefly, RNA was isolated from the cell pellets by adding them to reverse transcription solution. Immunoglobulin genes were reverse-transcribed using Superscript III (Thermo Fisher Scientific) with random hexamer oligonucleotides (Gene Link, Hawthorne, NY) as primers. The cDNA served as a template for nested PCR amplification of the antibody heavy and light chain genes. The PCR used mouse antibody gene-specific primers ^73^ and was carried out with AmpliTaq Gold 360 (Thermo Fisher Scientific). The PCR products were purified using the Biomek FX Laboratory Automation Workstation (Beckman Coulter, Brea, CA). Purified PCR products were then sequenced using Sanger sequencing to obtain the antibody gene sequences. The V(D)J rearrangement, somatic hypermutation frequency, and CDR3 (complementarity-determining region 3) length of the VH (heavy chain variable region) and VK/L (light chain variable region) genes were analyzed using a custom in-house bioinformatics pipeline for immunogenetic analysis. Annotation was performed using Partis ^74^ with a custom knock-in mouse reference that combined the knocked-in human V_H_1-2*02, J_H_2*01, and V_K_1-33*01 with mouse endogenous V, D and J gene segments.

### Cryo-EM grid Preparation and Imaging

To prepare grids, Quantifoil R1.2/1.3 (Cu, 300-mesh; Electron Microscopy Sciences, PA) grids were glow discharged for 15 seconds at 15 mA using a PELCO easiGlow Cleaning System (Ted Pella Inc). The final concentration of purified AngV-F, LayV-1-F, and LayV-F-DS ectodomains was maintained as 1.5 mg/ml in PBS, with a 5x molar ratio of 22F5 for LayV-1-F and 4.87x ratio for LayV-F-DS. 0.005 % (w/v) of n-dodecyl β-D-maltoside (DDM) was added to the AngV-F samples and 0.005% (w/v) DDM and 2% glycerol to the LayV-1-F and LayV-F-DS samples before applying to the grid to prevent the protein from adhering to the air-water interface. 3.0 μL of the protein sample was applied to the grid and incubated for 30 seconds at >95% humidity. Excess protein was blotted away for 2.5 seconds before being plunge frozen into liquid ethane using Leica EM GP2 plunge freezer (Leica Microsystems). Cryo-grids were imaged using either an FEI Titan Krios (Thermo Scientific) or Tundra cryo TEM (Thermo Scientific).

The Titan Krios TEM was used for the determination of the AngV-F trimer, dimer-of-trimer, and 22F5 Fab-LayV-1-F complex structures. The Krios was equipped with a K3 camera (Gatan) at 81k magnification, operated at 300keV. For AngV-F, approximately 24,000 micrographs were collected at nominal defocused range between -2.8 to -1.7 μm with a dose range of 63-65 e/A^2^. For LayV-1-F, 9,715 micrographs were collected at a nominal defocused range between -2.8 to - 1.2 µm with a dose range of 58.6-60.6 e/A^2^.

The Tundra TEM was used for determination of the AngV-F hexameric lattice and 22F5 Fab-LayV-F-DS complex structures. For the AngV-F hexameric lattice, the Tundra was equipped with a CETA-F camera (Thermo Scientific) at 180k magnification, operated at 100keV. Approximately 4,000 micrographs were collected at nominal defocused range between -3.0 to - 1.5 μm with a dose range of 23-25 e/A^2^. For the 22F5 Fab-LayV-F-DS complex, the Tundra was equipped with a Falcon 4i camera (Thermo Scientific) at 180k magnification, operated at 100keV. 9,514 micrographs were collected at a nominal defocused range between -2.5 to -0.7 µm with a dose of 30.72 e/A^2^.

#### Cryo-EM Data Processing

Cryo-EM image quality was monitored on-the-fly during data collection using automated processing routines. Data processing was carried out using cryoSPARC ^75,76^. Micrographs were curated through contrast transfer function (CTF) where greater than >30 Å were discarded. Automated blob picker was used to assign the particle position. Different box sizes are utilized to accommodate the varying dimensions of specific molecular assemblies. For instance, monomer particles are extracted using a box size of 320 pixels, while dimer particles require a larger box size of 512 pixels. For even larger molecular assemblies, such as hexamer lattices, a box size of 1024 pixels was employed. Following particle extraction, multiple rounds of 2D classification was performed to remove junk. *ab-initio* 3D reconstruction was used to create 3D reconstructions and poor-quality particles were discarded after thorough heterogeneous refinement. Final resulting volumes were subjected to non-uniform refinement to build a high-resolution 3D reconstruction. Phenix ^77^, Coot ^78^, Pymol, nextPYP ^79^ and ChimeraX ^80^ were used for model building and refinement. Models 8TVE ^20^ and 8U1R ^30^ were used as startup models for post-fusion and pre-fusion LayV-F model building, respectively.

### X**-** ray crystallography

Crystals were grown via the vapor diffusion method at room temperature in a siting drop well format. GakV-2-G head domain protein crystals were formed by mixing 0.20 μL of protein at 15 mg/mL with 0.20 μL of well solution containing 0.2 M magnesium sulfate heptahydrate, 20% w/v PEG3350 at pH 7.1. Crystals appeared within one week.

Cryoprotection of the crystals was done by mixing glycerol to a final concentration of 50% with the well solution. Crystals were moved to the drop containing the well solution and glycerol and soaked for one minute then flash-cooled in liquid nitrogen and data were collected at the NSLS-II, Beamline 19-ID at 100 K and a wavelength of 1 Å. All data were indexed and integrated using iMosflm ^81^ and scaled using AIMLESS. Phaser ^82^ in the PHENIX suite was used to perform molecular replacement using a model generated from the AlphaFold3 prediction server ^83^ using the GakV-2-G head domain sequence. Phenix refine ^84^ was used for data refinement, and manual refinement was done in Coot ^78^.

## Notes

### Competing Interest Statement

A.M., M.L., J.J.L, A.B., B.F.H. and P.A. are named in provisional patents submitted on the methods for stabilizing the pre-fusion HNV F and G proteins based on the findings reported in this paper. Other authors declare no competing interests.

### Summary of Updates

Several figures have been reorganized, expanded, or split. A new main figure and several supplemental figures have been added, along with updated text to match.

