## Supplemental Figures 1-25, Supplemental Note 1, Supplemental Tables 2-3, Supplemental Data 1-2 for "Mechanistic and Antigenic Boundaries of *Henipavirus* and *Parahenipavirus* Glycoproteins"

[$]Lead contact

#These authors contributed equally


**This PDF File Includes:**

Supplemental Note 1

Supplemental Figures 1-25

Supplemental Tables 2-3

Supplemental Data 1-2

Note: Supplemental Table 1 provided as a separate excel file.

#### Supporting Material Contents

**Content** **Page(s)**


| Content | Page(s) |
| --- | --- |
| Supplemental Note 1: Classification of HNV Species | 3-4 |
| Supplemental Figure 1: Yields and Protein Similarity for HNV-F and G proteins | 5 |
| Supplemental Figure 2: Amino Acid-based Phylogeny of HNV Species | 6 |
| Supplemental Figure 3: Size Exclusion Chromatography (SEC) Profiles of HNV-F Ectodomains | 7-10 |
| Supplemental Figure 4: SDS-PAGE Profiles of HNV-F Ectodomains | 11 |
| Supplemental Figure 5: NSEM Analysis of HNV-F Ectodomains | 12-15 |
| Supplemental Figure 6: Mouse Immunizations with HNV-F Proteins | 16 |
| Supplemental Figure 7: Measurement of Antibody Binding to HNV-F Proteins using Biolayer Interferometry (BLI) | 17 |
| Supplemental Figure 8: Measurement of Antibody Binding to HNV-F Proteins using ELISA | 18 |
| Supplemental Figure 9: Measurement of Antibody Binding to HNV-F Proteins | 19-20 |
| Supplemental Figure 10: SPR Analysis of 22F5 Binding to LayV-F Ectodomain | 21 |
| Supplemental Figure 11: Cryo-EM Data Processing Workflow for LayV-C1-F Ectodomain and 22F5 Fab Complex | 22 |
| Supplemental Figure 12: Cryo-EM Data Processing Workflow for LayV-F-DS Ectodomain and 22F5 Fab Complex | 23 |
| Supplemental Figure 13: Model to map fit of 22F5-LayV-F structures | 24 |
| Supplemental Figure 14: Heat Incubation of HNV-F Proteins | 25 |
| Supplemental Figure 15: DSF-relevant Residue Variability | 26 |
| Supplemental Figure 16: Workflow of Particle Extraction and 2D Classification | 27 |
| Supplemental Figure 17: Cryo-EM Data Processing and Refinement Outline of Monomer Pre-fusion AngV-F Protein | 28 |
| Supplemental Figure 18: Oligomeric States of AngV-F and HeV-α1.2-F | 29 |
| Supplemental Figure 19: AngV-F Glycan Distribution and Differences Compared to other HNV-F Proteins | 30 |
| Supplemental Figure 20: Characterization of HRB Start Proline HNV-F Mutants | 31 |
| Supplemental Figure 21: NSEM 3D Reconstruction of HRB Start Proline Mutants | 32 |
| Supplemental Figure 22: Size Exclusion Chromatography (SEC) Profiles of HNV-G Head Domains | 33 |
| Supplemental Figure 23: HNV-G Binding to Ephrin and Antibodies | 34-39 |
| Supplemental Figure 24: Thermostability of HNV G Head Domains | 40 |
| Supplemental Figure 25: Head Domain Comparison | 41 |
| Supplemental Table 2. Cryo-Em Data Collection and Refinement Statistics | 42 |
| Supplemental Table 3. Data Collection and Refinement Statistics (Molecular Replacement) | 43 |
| Supplemental Data 1: Sequence Alignment of HNV-F Proteins | 44-46 |
| Supplemental Data 2: Sequence Alignment of HNV-G Proteins | 46-49 |

**Supplemental Note 1: Classification of HNV Species**

For clarity in referring to specific sequences with greater granularity than the species level, we based our abbreviations and numbering on conventions of Nipah virus naming in the literature. Previously, strains have been named based on the location where the sequence was reported, specifically, Bangladesh (NiV-B) and Malaysia (NiV-M). Recent discovery and classification of new Nipah virus strains support the existence of potentially two further geographical groupings, India (NiV-I) and Cambodia (NiV-C) [57,58], and these distinctions are apparent in our tree as well. With the Bangladesh strains, the phylogeny revealed the grouping of strains into two clades, prompting us to denote the two groupings as NiV-B1 and NiV-B2, respectively. From here, unique entries were designated an additional number to further classify the variant (i.e. NiV-B2.1, NiV-B2.2, NiV-B3.1, etc.). An identical scheme was applied for the India strains, which previously have been referred to as members of the Bangladesh clade [57]. Malaysia strains were classified by number as well. Entry MK801755 was identified during surveillance in Cambodia in 2003, where it was isolated from *Pteropus lylei* fruit bats [59], and as the only unique NiV-C member of the panel, requires no numbering.

Previous studies have established two clades for Hendra virus [60], in which the more recently discovered clade is referred to as HeV-g2. However, due to the use of the letter G as an abbreviation for the attachment protein, we sought to adopt a different naming scheme to avoid confusion. Unlike Nipah, where genomic differences align with country borders, all Hendra virus discoveries have been in Australia. Therefore, the two distinct clades have been named Alpha (α) and Beta (β), with each member of the clade given a number. The other Henipaviruses that have emerged from a fruit bat reservoir have all previously been named in the literature. These include Cedar Virus (CedV, [61]), Ghana Virus (GhV, [62]), and Angavokely Virus (AngV, [63]). Our panel includes two unique Cedar virus attachment proteins; therefore, these strains were numbered 1 and 2. There are only singular strains of Ghana and Angavokely viruses identified, however some evidence indicated that the originally deposited sequence for Ghana virus F protein may have included an inaccurate N-terminal end of the protein and a corrected sequence was suggested [64]. While our phylogenetic tree only includes the one strain reported, our panel of purified proteins included both F protein sequences.

Among the species found in shrews, now officially classified as members of the *Parahenipavirus* genus, several already have established names in the literature, including Mojiang Virus (MojV), Langya Virus (LayV), Gamak Virus (GakV), Denwin Virus (DewV), Melian Virus (MelV), and Ninorex Virus (NinV) [5-7,65]. For those with more than one strain reported, the same numbering approach used with the Nipah-Bangladesh clade was used. In the case of Langya and Daeryong viruses, distinct clades are found with geographic separation, as with Nipah virus. Similarly, these clades were assigned as Korea and China for clarity.

Many of the recently discovered *Parahenipavirus* sequences have been given repetitive names based on their shared collection site and host species. However, in some situations, this could imply overly close or distant phylogenetic relationships between strains. For example, strains listed as Jingmen Crocidura shantungensis virus 1 (e.g. OM030314) [9] are not closely related to those listed as Jingmen Crocidura shantungensis virus 2 (e.g. OM030315). Inversely, strains listed as Wenzhou shrew Henipavirus 1 (OQ715593.1) and Wenzhou Apodemus agrarius Henipavirus 1 (MZ328275.1) were shown to be quite closely related (**Figure 1).** For simplicity and clarity, we grouped many of these sequences using variants of the name "Shrew Henipavirus (SHNV)" with a number, thus establishing SHNV1 through SHNV11. For SHNV5 and SHNV10 specifically, there were several strains that the phylogenetic tree indicated were closely related enough to be considered the same species (**Figure 1**). As a result, we utilized a numbering scheme as described above.

Lee et al,. 2020 [6] previously reported the discovery of Daeryong Virus in Korea, and our phylogenetic analysis indicates that two strains listed as Jingmen Crocidura shantungensis virus 2 (OM030315, PP272750.1), detected in China, could be considered the same species as Daeryong Virus. Following the naming procedure used for Nipah virus, the strains were given the names Daeryong virus-Korea (DarV-K) and Daeryong virus-China (DarV-C).This demonstrates that several of these newly discovered Parahenipaviruses are geographically well-distributed. A full listing of the panel, including original deposition names, accession codes, genus, and sequence ranges used, is available as **Supplemental Table 1.**

**A** HNV-F Protein Yields and Similarities

| Name | Identity to NiV-M1-F | Similarity to NiV-M1-F | Main Peak Yield/Liter (mg/L) | Main Peak Yield (Percentage of Total) |
|---|---|---|---|---|
| NiV-M1-F | N/A | N/A | 0.525 | 50.7% |
| NiV-M2-F | 99.82 | 99.82 | 0 | N/A |
| NiV-M3-F | 99.82 | 99.82 | 1.8 | 35.2% |
| NiV-M4-F | 99.45 | 99.45 | 0 | N/A |
| NiV-B1.2-F | 99.08 | 99.63 | 1 | 26.8% |
| NiV-C-F | 98.9 | 99.63 | 0.5 | 70.3% |
| NiV-B1.3-F | 98.9 | 99.45 | 0.425 | 46.6% |
| NiV-B1.4-F | 98.72 | 99.45 | 0.03 | 50.8% |
| NiV-B2.3-F | 98.53 | 99.27 | 0.14 | 65.0% |
| NiV-B1.1-F | 98.53 | 99.08 | 0 | N/A |
| NiV-B3.1-F | 98.17 | 98.72 | 0.38 | 25.1% |
| HeV-α1.2-F | 88.3 | 94.33 | 0.8 | 35.1% |
| HeV-α1.1-F | 88.12 | 94.15 | 0.305 | 57.1% |
| HeV-β1-F | 87.91 | 93.22 | 0.65 | 55.9% |
| HeV-β2-F | 87.73 | 93.41 | 0.1 | 65.7% |
| GhV(+A)-F | 51.5 | 69.31 | 0.147 | 58.5% |
| GhV-F | N/A | N/A | 0 | N/A |
| CedV-1-F | 42.93 | 61.72 | 0.05 | 63.7% |
| MojV-F | 41.24 | 61.86 | 0.335 | 38.0% |
| SHNV5-1-F | 40.8 | 61.75 | 5.7 | 23.8% |
| SHNV5-2-F | 40.8 | 61.57 | 2.1 | 25.0% |
| LayV-C1-F | 40.77 | 60.33 | 1.17 | 65.6% |
| LayV-C2-F | 40.77 | 60.33 | 0.85 | 53.1% |
| SHNV2-F | 40.61 | 61.55 | 2.1 | 36.9% |
| DarV-K-F | 40.59 | 61.61 | 0 | N/A |
| SHNV4-F | 40.56 | 57.32 | 0.165 | 41.6% |
| DarV-Cβ-F | 40.4 | 61.06 | 0 | N/A |
| GakV-1-F | 39.86 | 58.02 | 0.44 | 58.6% |
| GakV-2-F | 39.86 | 57.85 | 0.445 | 61.0% |
| DewV-1-F | 39.85 | 62.52 | 0 | N/A |
| MelV-F | 39.71 | 61.93 | 0 | N/A |
| SHNV3-F | 39.51 | 57.14 | 0.31 | 41.1% |
| AngV-F | 39.27 | 57.64 | 1.35 | 72.2% |
| SHNV11-F | 38.7 | 55.48 | 0.2 | 40.7% |
| NinV-F | 38.02 | 55.11 | 0.9 | 59.1% |
| SHNV1-F | 37.39 | 56.26 | 2.15 | 31.1% |

**B** HNV-G Protein Yields and Similarities

| Name | Identity to NiV-M1-G | Similarity to NiV-M1-G | Head Domain Yield/Liter (mg/L) |
|---|---|---|---|
| NiV-M1-G | N/A | N/A | 26.52 |
| NiV-M3-G | 99.5 | 99.5 | 18.75 |
| NiV-B1.5-G | 95.85 | 97.01 | 26.7 |
| NiV-B3.1-G | 95.68 | 96.84 | 26.7 |
| NiV-B2.4-G | 95.51 | 97.01 | 31.2 |
| NiV-B1.6-G | 95.51 | 96.68 | 25.2 |
| NiV-I1.3-G | 95.18 | 96.68 | 13.45 |
| HeV-β3-G | 78.97 | 87.42 | 15.4 |
| HeV-β1-G | 78.64 | 87.09 | 12.75 |
| HeV-α6-G | 78.31 | 87.42 | 17 |
| HeV-α5-G | 78.31 | 87.42 | 16.15 |
| HeV-α3-G | 78.31 | 87.42 | 12.75 |
| HeV-α1.2-G | 78.15 | 87.42 | 12.75 |
| HeV-α4-G | 77.98 | 87.25 | 6.95 |
| CedV-1-G | 29.65 | 45.43 | 16.07 |
| GhV-G | 25.08 | 40.46 | 6.75 |
| AngV-G | 17.59 | 30.61 | 8.15 |
| MojV-G | 17.57 | 33.18 | 7.1 |
| SHNV5-1-G | 17.1 | 31.74 | 35.3 |
| SHNV5-2-G | 17.1 | 31.74 | 19.95 |
| DarV-K-G | 16.94 | 35.68 | 5.2 |
| DarV-Cβ-G | 16.94 | 35.68 | 2.45 |
| LayV-C2-G | 16.84 | 32.18 | 17.5 |
| LayV-C3-G | 16.69 | 32.18 | 14 |
| MelV-G | 16.64 | 33.43 | 6.4 |
| DewV-1-G | 16.27 | 34.04 | 7.75 |
| SHNV3-G | 15.83 | 32.69 | 14.6 |
| SHNV11-G | 15.68 | 32.1 | 6.55 |
| NinV-G | 15.41 | 30.67 | 3.085 |
| GakV-1-G | 15.38 | 31.8 | 14.8 |
| GakV-2-G | 15.38 | 31.8 | 19.3 |
| SHNV2-G | 14.54 | 32.08 | 3.25 |

**Supplemental Figure 1: Yields and Protein Similarity for HNV-F and G proteins.** Table of protein yields and sequence similarities for each member of the protein panel. Percent identities and similarities of the amino acid sequences of the full-length proteins compared to NiV-M1 are listed. Color scheme for HNV species continued from Figure 1. **(A)** Table for HNV-F proteins. GhV-F similarity values were excluded due to the likely inaccurate N-terminal sequence in the deposited sequence. Main peak yield is based on the final calculated protein yield after SEC. Main peak percentage of total is based on SEC area under the curve of the main peak as a fraction of all collected SEC fractions. **(B)** Table for HNV-G proteins.

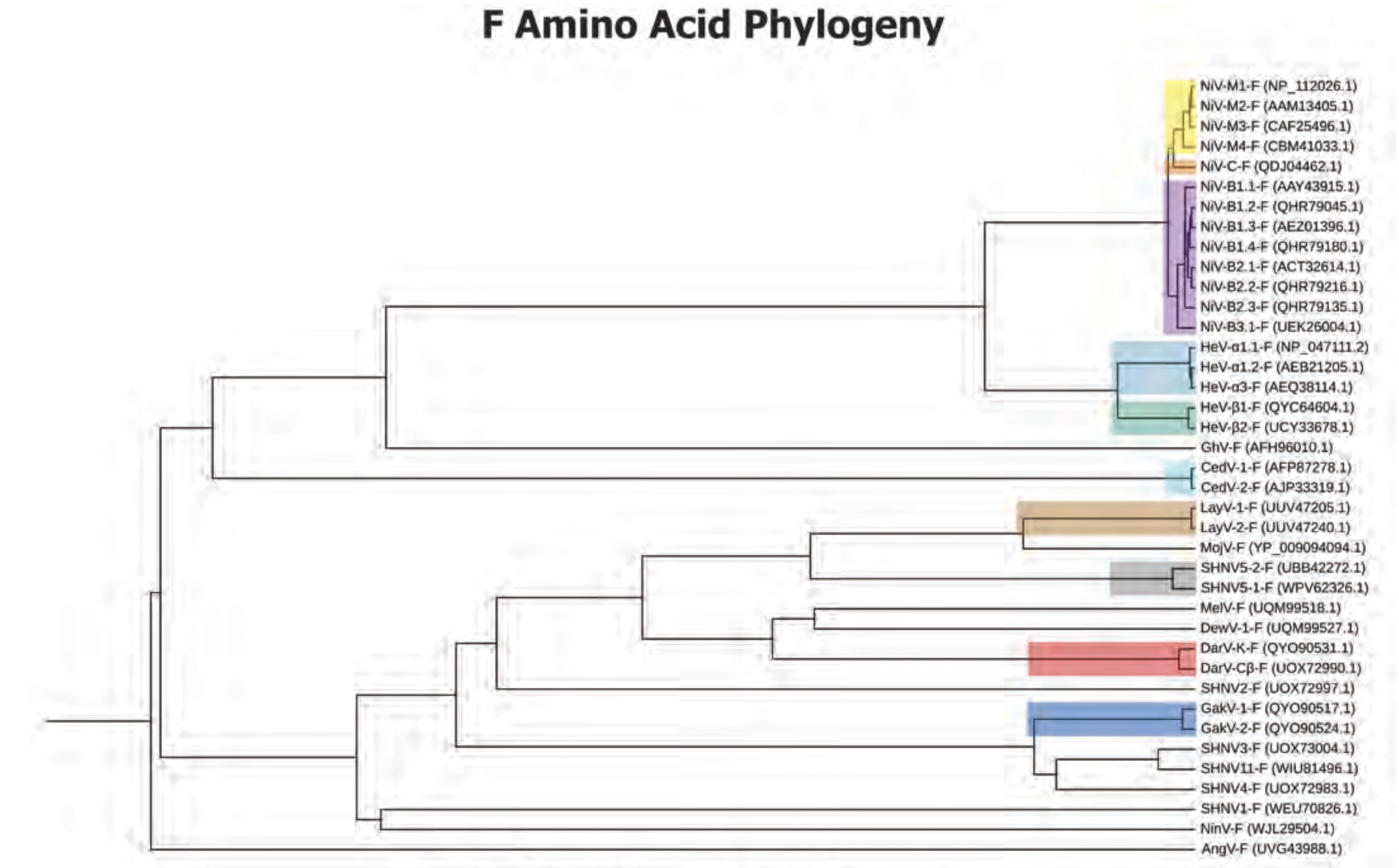


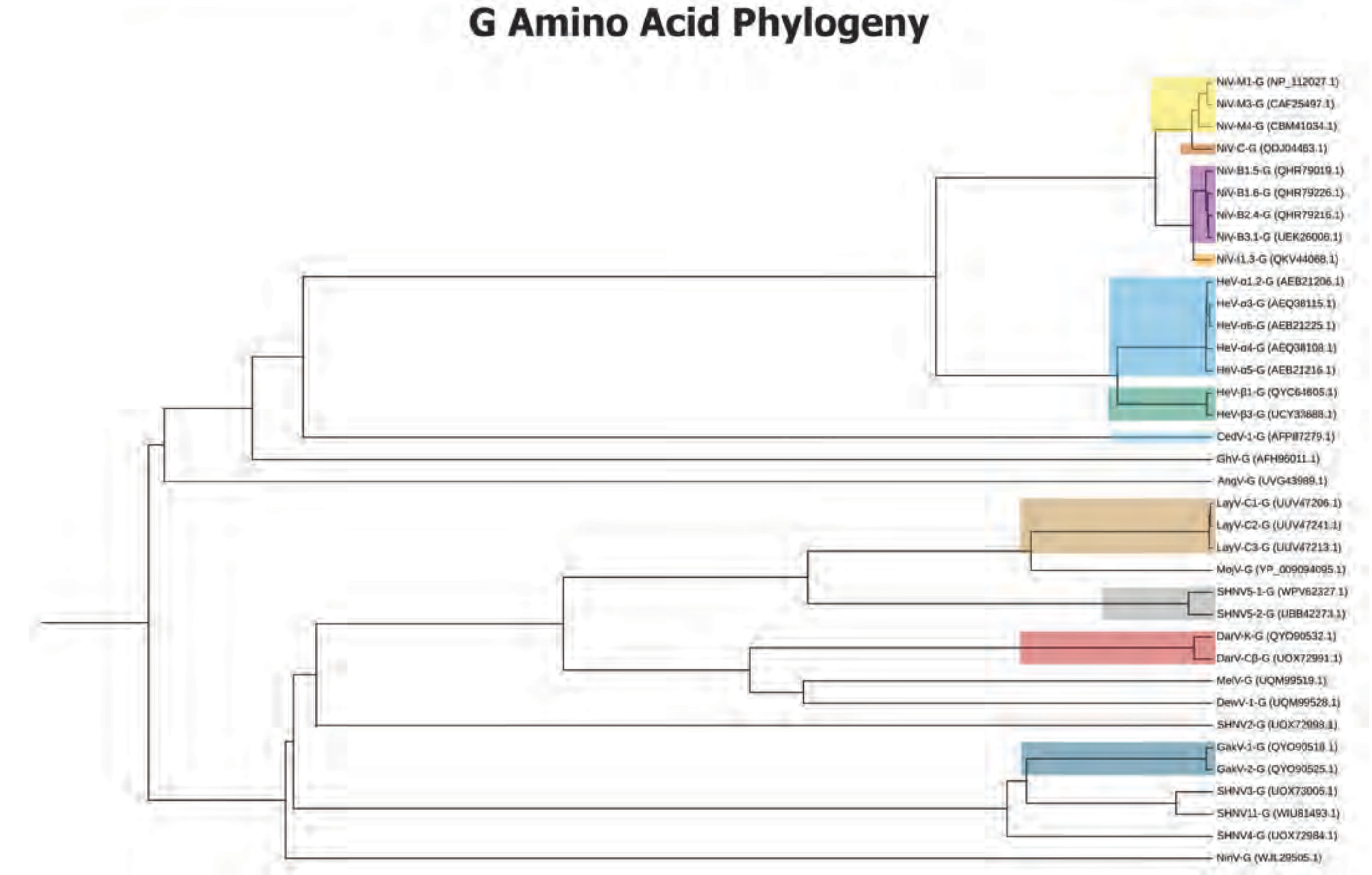


**Supplemental Figure 2: Amino Acid-based Phylogeny of HNV Species.** Phylogenetic trees constructed using either F or G amino acid sequences for purified selections from the full tree in Figure 1. Genbank amino acid accession codes listed next to each tip. Color scheme is continued from Figure 1.

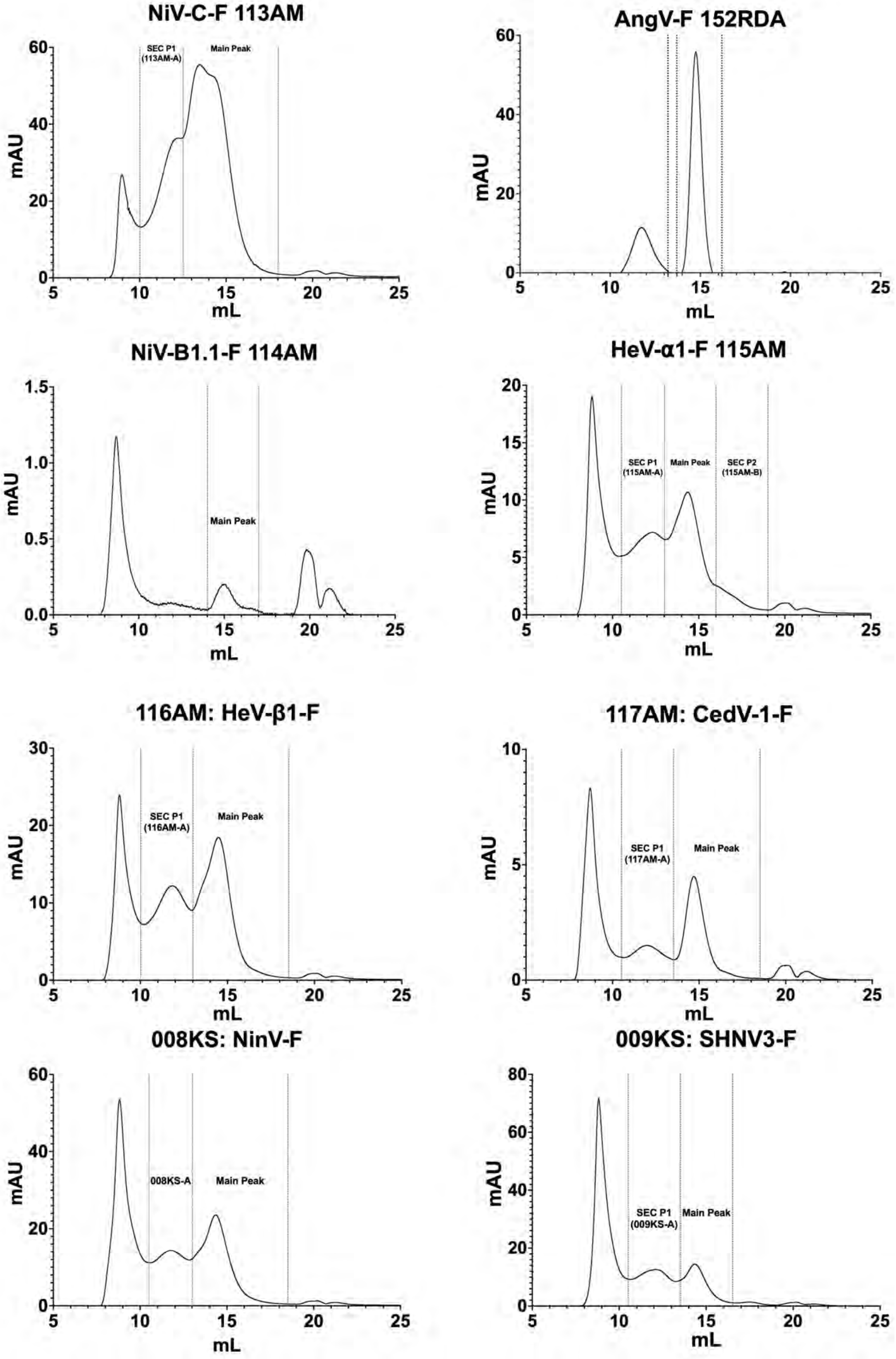


**Supplemental Figure 3: Size Exclusion Chromatography (SEC) Profiles of HNV-F Ectodomains.** Fraction boundaries of collected peaks are labelled. Internal production lot numbers included with strain names.

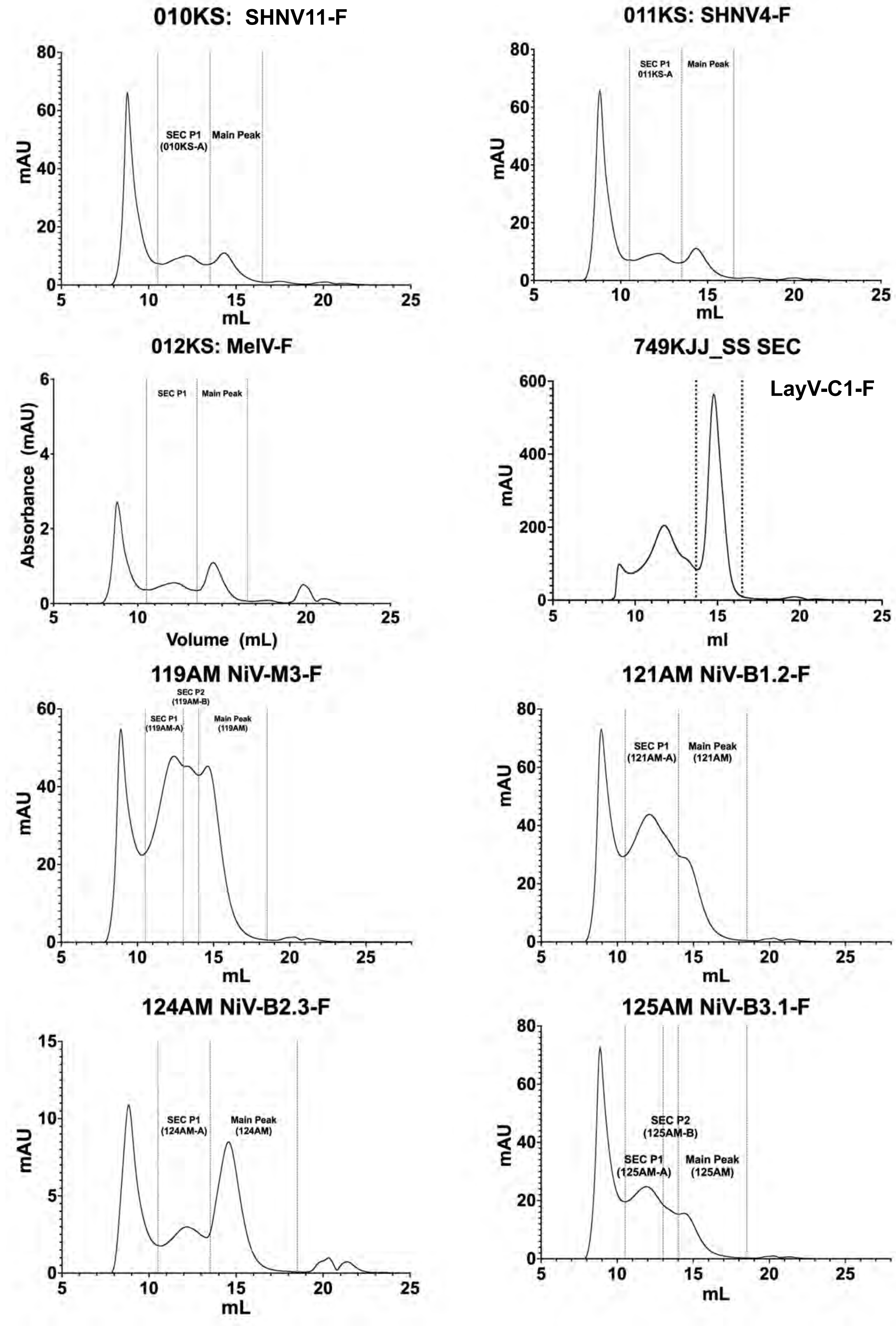


**Supplemental Figure 3 (continued): Size Exclusion Chromatography (SEC) profiles of HNV F ectodomains.**

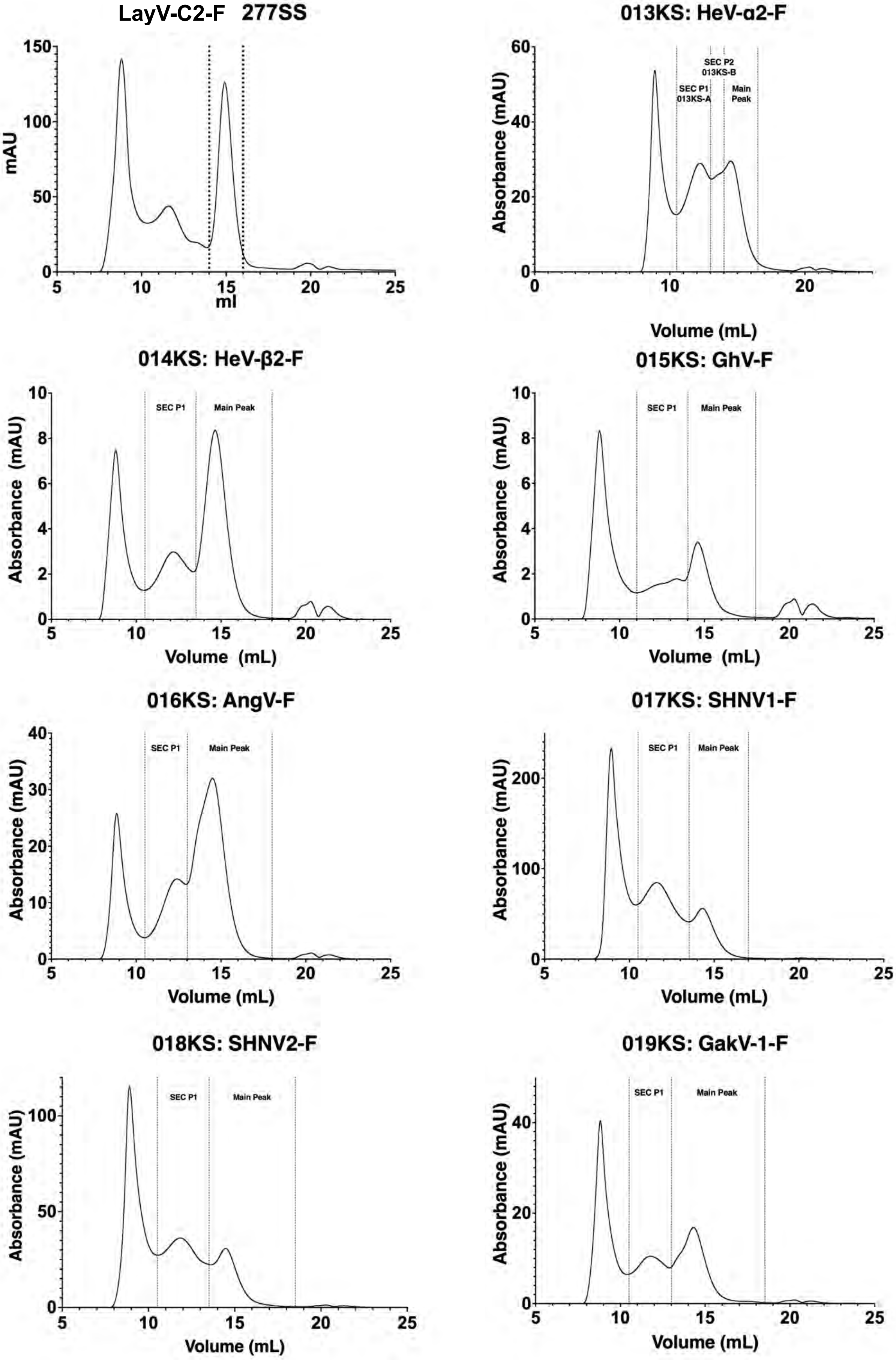


**Supplemental Figure 3 (continued): Size Exclusion Chromatography (SEC) Profiles of HNV-F Ectodomains.** Fraction boundaries of collected peaks are labelled. Internal production lot numbers included with strain names.

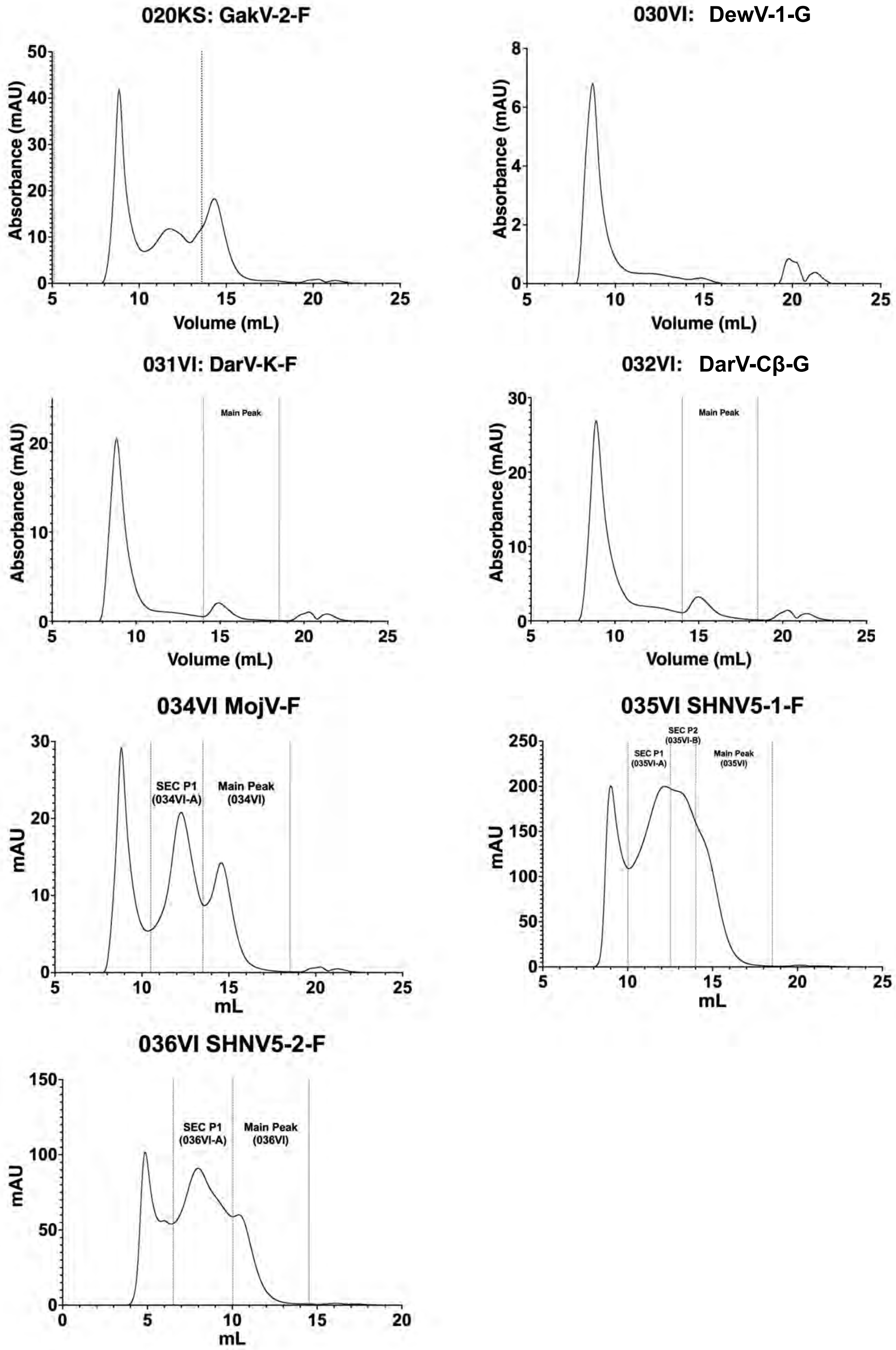


**Supplemental Figure 3 (continued): Size Exclusion Chromatography (SEC) Profiles of HNV-F Ectodomains.** Fraction boundaries of collected peaks are labelled. Internal production lot numbers included with strain names.

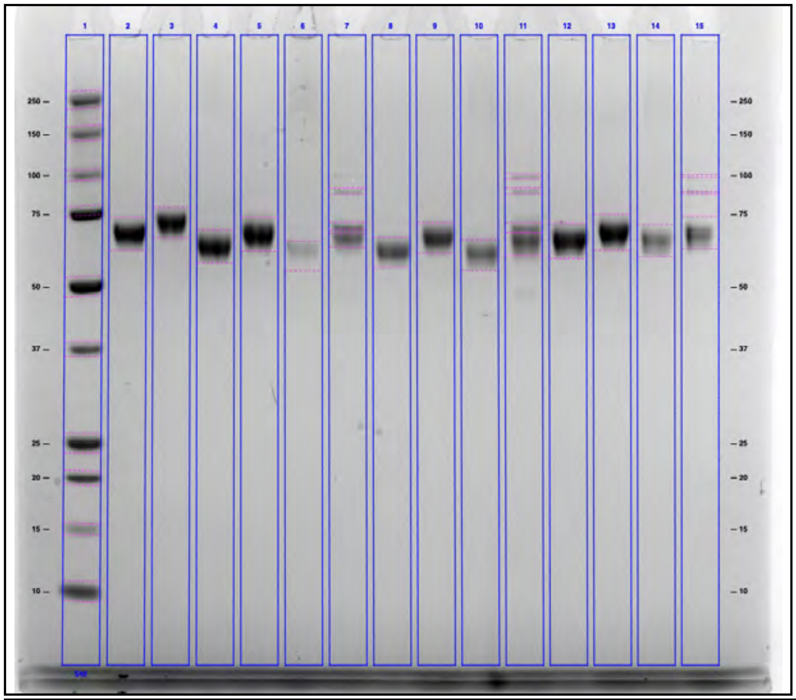


Protocol: 15 µL of each sample was loaded both reduced and non-reduced. The sample was boiled @ 95˚C for 3 min.
DTT: 1M DTT
Protein Marker: Precision Plus Protein Unstained Standards
Sample Buffer: 4X Laemmli Sample buffer
Gel: NuPAGE 4-12% BT 1.0, 15-well combs

| Lane | Sample Name | Amount Loaded (µg) | Volume Loaded (µL) | Reduced (R/NR) | Molecular Weight (kDa) |
|---|---|---|---|---|---|
| 1 | Ladder | N/A | 4 | N/A | N/A |
| 2 | 117AM: CedV-1-F | 1 | 15 | NR | 67.3 |
| 3 | 117AM: CedV-1-F | 1 | 15 | R | 71.5 |
| 4 | 123AM: NiV-B1.4-F | 0.68 | 15 | NR | 62.4 |
| 5 | 123AM: NiV-B1.4-F | 0.68 | 15 | R | 66.8 |
| 6 | 123AM-A: NiV-B1.4-F SEC P1 | 1 | 15 | NR | 61.8 |
| 7 | 123AM-A: NiV-B1.4-F SEC P1 | 1 | 15 | R | 88.9, 69.5, 65.8 |
| 8 | 124AM: NiV-B2.3-F | 1 | 15 | NR | 60.8 |
| 9 | 124AM: NiV-B2.3-F | 1 | 15 | R | 65.5 |
| 10 | 124AM-A: NiV-B2.3-F SEC P1 | 0.81 | 15 | NR | 60.3 |
| 11 | 124AM-A: NiV-B2.3-F SEC P1 | 0.81 | 15 | R | 99.2, 88.9, 68.9, 64.7 |
| 12 | 014KS: HeV-β2-F | 1 | 15 | NR | 64.7 |
| 13 | 014KS: HeV-β2-F | 1 | 15 | R | 67.5 |
| 14 | 014KS-A: HeV-β2-F SEC P1 | 1 | 15 | NR | 64.9 |
| 15 | 014KS-A: HeV-β2-F SEC P1 | 1 | 15 | R | 98.9, 88.3, 67.1 |

Notes: NR: 3.75 µL Dye, Protein, PBS
R: 3.75 µL dye, 4.5 µL DTT , Protein, PBS

**Supplemental Figure 4: SDS-PAGE Profiles of HNV-F Ectodomains.** Representative example of an SDS-PAGE profile shown for HNV F ectodomains. Lane 2 and 3: CedV-1-F (NR= Non-reduced; R= Reduced). Lanes 4 and 5: Main (trimer) peak of NiV-B1.4-F. Lanes 6 and 7: Aggregate of trimers (Middle peak; P1) of NiV-B1.4-F. Lanes 8-11: NiV-B2.3-F, same order of samples as for NiV-B1.4-F. Lanes 12-15: Hev-β2-F, same order of samples as for NiV-B1.4-F. Internal production lot numbers included with strain names.

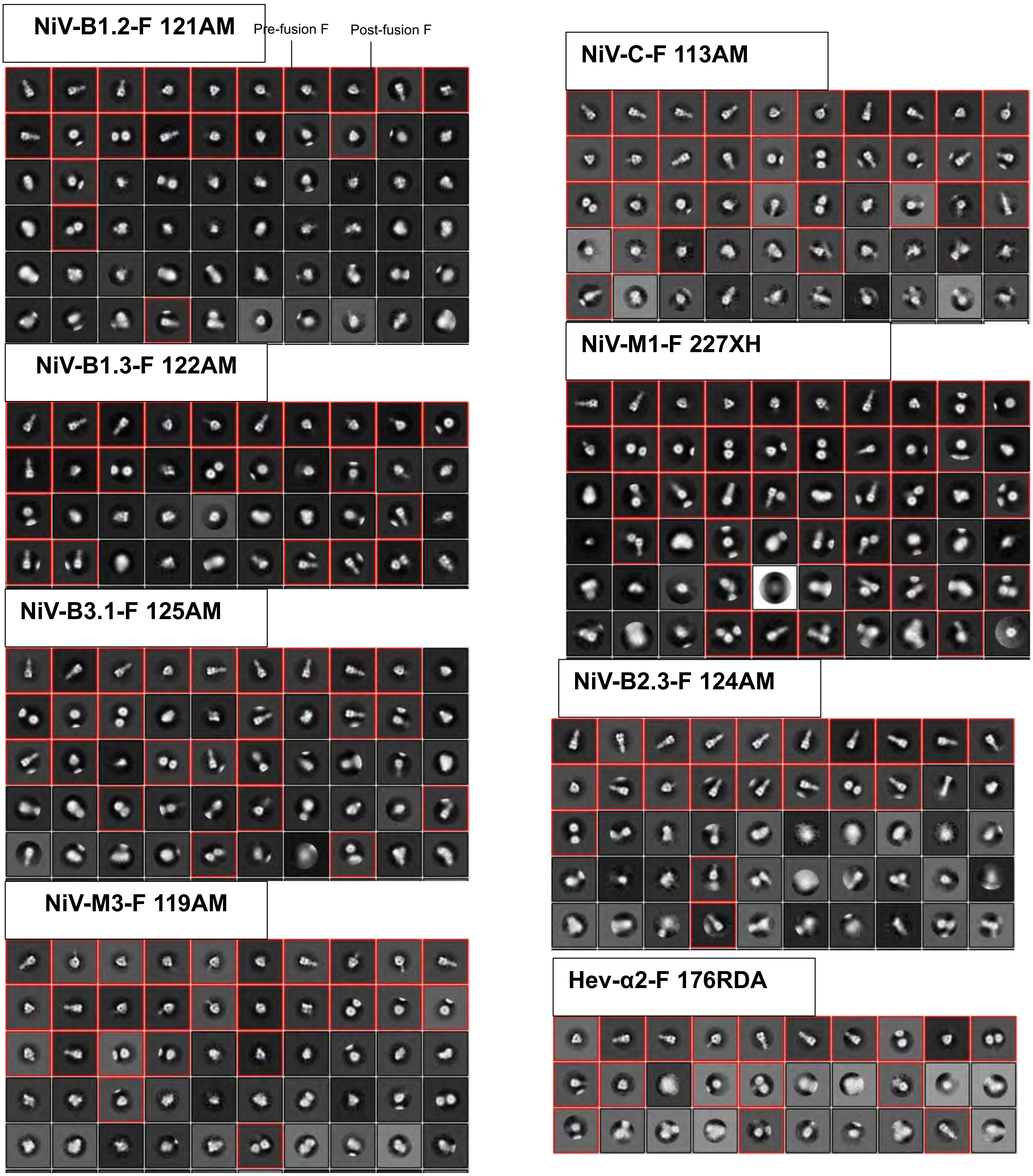


**Supplemental Figure 5: NSEM Analysis of HNV-F Ectodomains.** Representative 2D classes are shown. Classes containing the pre-fusion or post-fusion F protein can be identified by their shapes (as shown for NiV-B1.2-F as representative examples). The classes that correspond to the F proteins are identified by their typical shape and distinguishable from junk classes. Internal production lot numbers included with strain names.

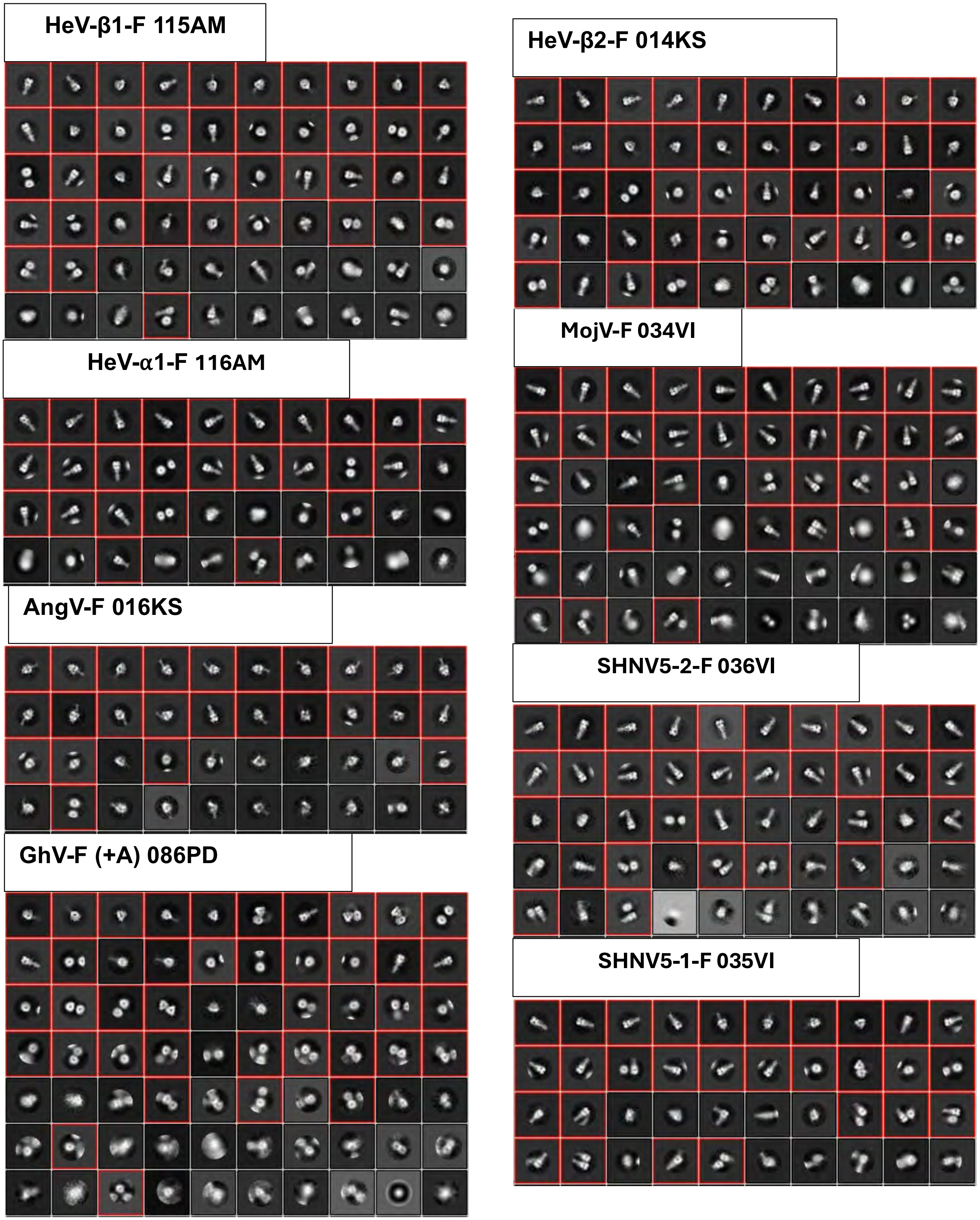


**Supplemental Figure 5 (continued): NSEM Analysis of HNV-F Ectodomains.** Representative 2D classes are shown. Internal production lot numbers included with strain names.

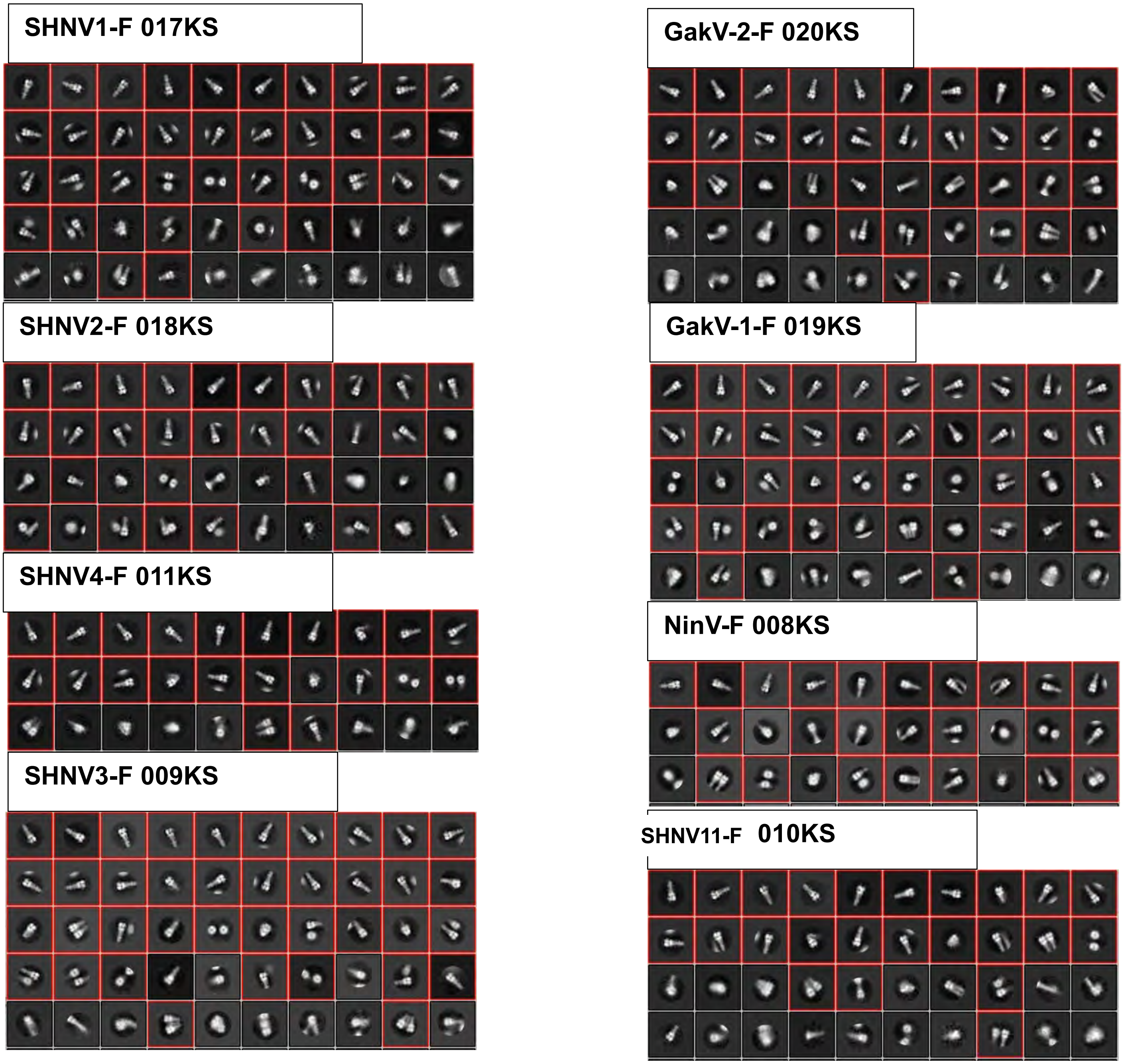


**Supplemental Figure 5 (continued): NSEM Analysis of HNV-F Ectodomains.** Representative 2D classes are shown. Internal production lot numbers included with strain names.

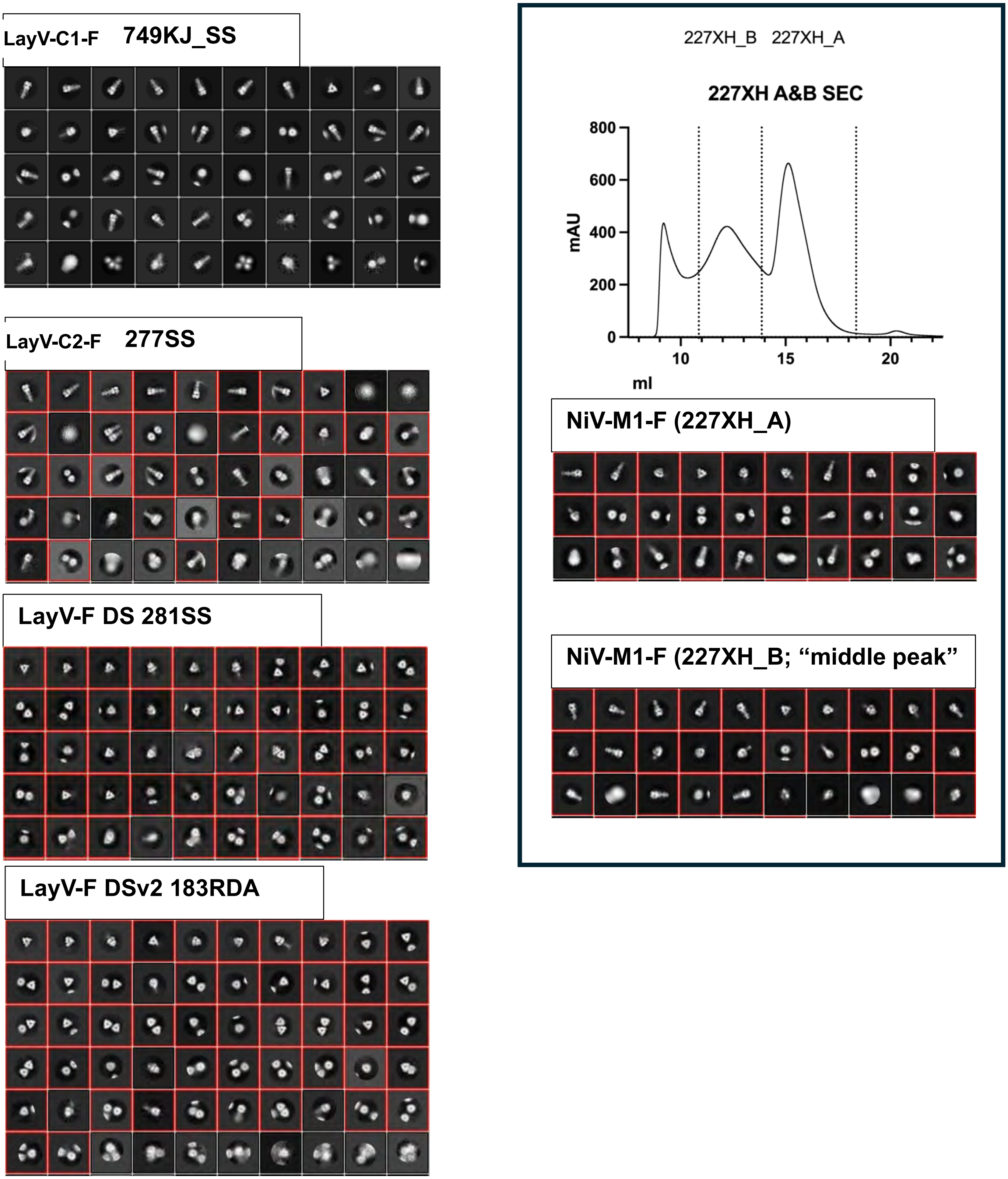


**Supplemental Figure 5 (continued): NSEM Analysis of HNV-F Ectodomains.** Representative 2D classes are shown. Right panel shows SEC profile of a NiV-M1-F ectodomain preparation with panels below showing representative NSEM 2D classes from the two peaks indicated by vertical dotted lines in the SEC profile. Internal production lot numbers included with strain names.

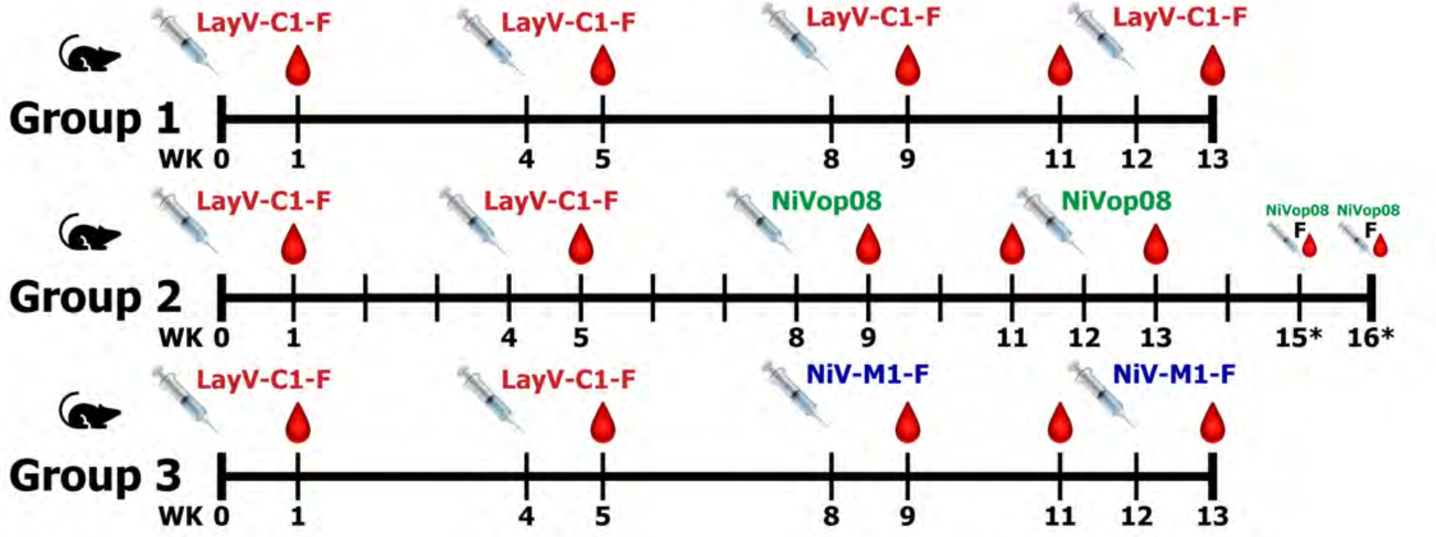


**Supplemental Figure 6: Mouse Immunizations with HNV-F Proteins:** Graphical timeline of a vaccination study in mice using HNV-F antigens. Each group was vaccinated with LayV-C1-F and boosted with LayV-C1-F or NiV antigens as marked. Blood draws and antigen injections occurred at the indicated weeks after the start of the study. The weeks with an asterisk, with the syringe and blood drop marked with F are weeks where a fusion was performed on an individual mouse.

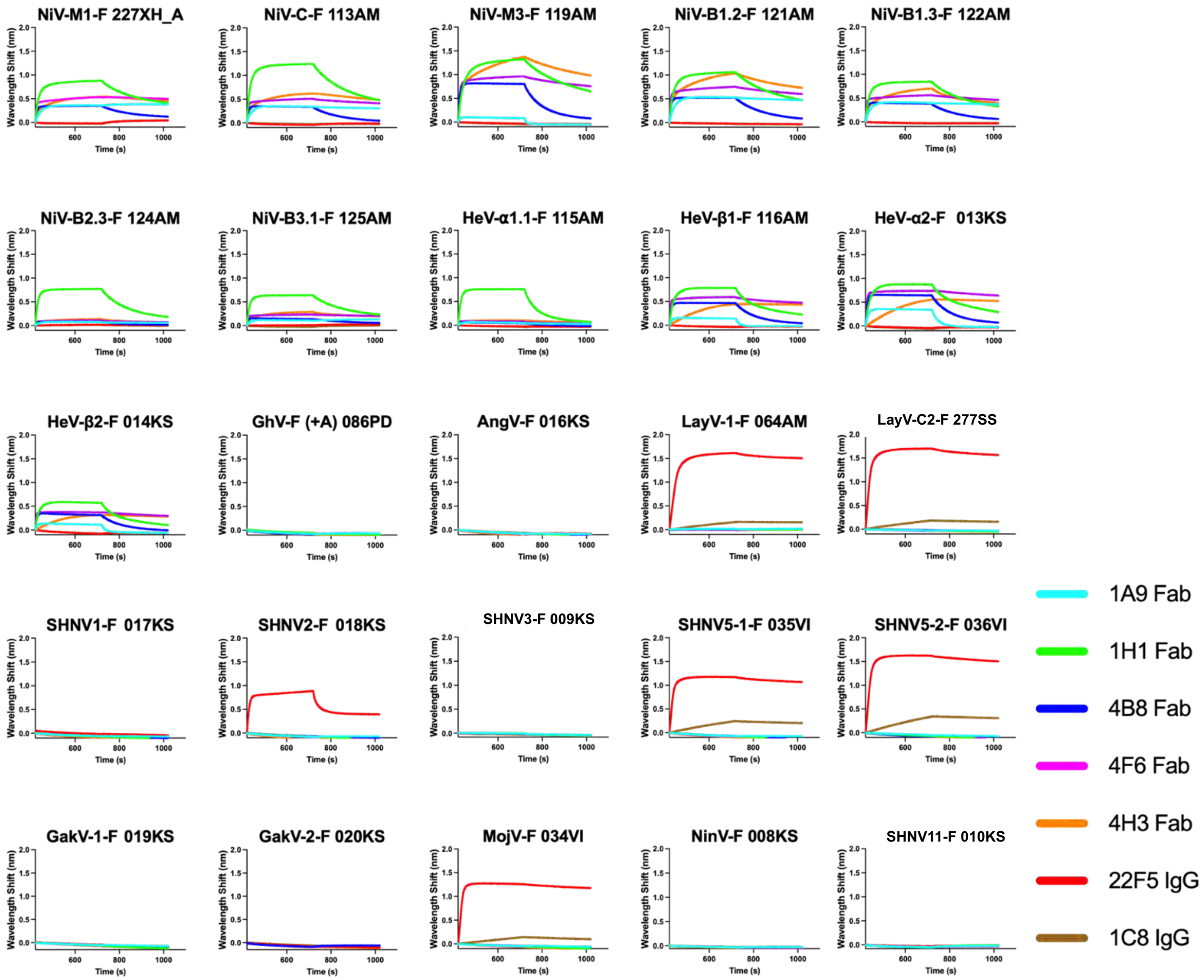


**Supplemental Figure 7: Measurement of Antibody Binding to HNV-F Proteins using Biolayer Interferometry (BLI).** The F proteins were captured on a BLI sensor tips via their C terminal TwinStrep tags. Binding to Fabs (1A9, 1H1, 4B8, 4F6, 4H3) or the vaccine elicited antibodies (22F5, 1C8) were measured. For each graph the virus from which the F ectodomain is derived is indicated at the top along with the lot number of the F ectodomain preparation. Internal production lot numbers included with strain names.

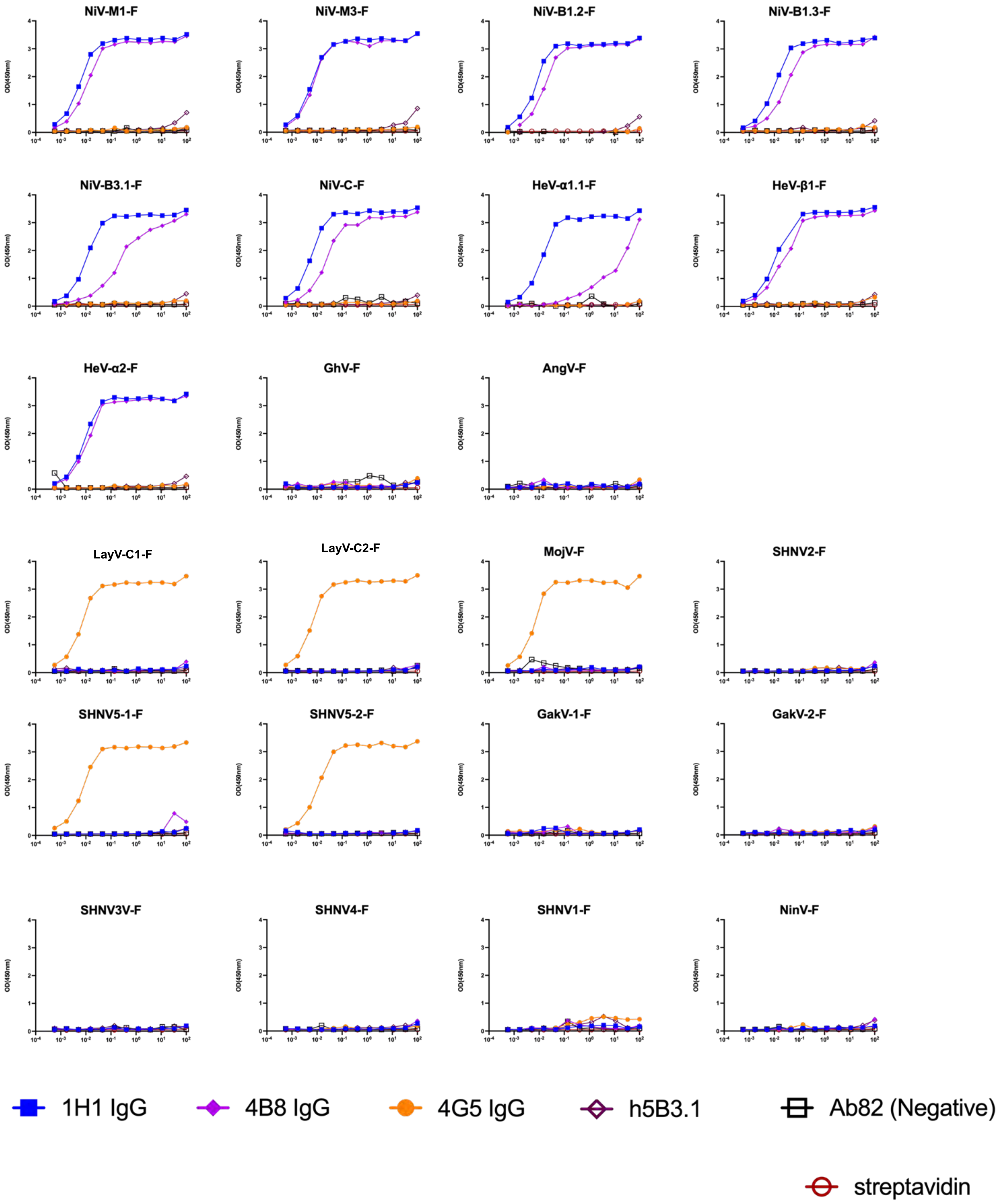


**Supplemental Figure 8: Measurement of Antibody Binding to HNV-F Proteins using ELISA.**

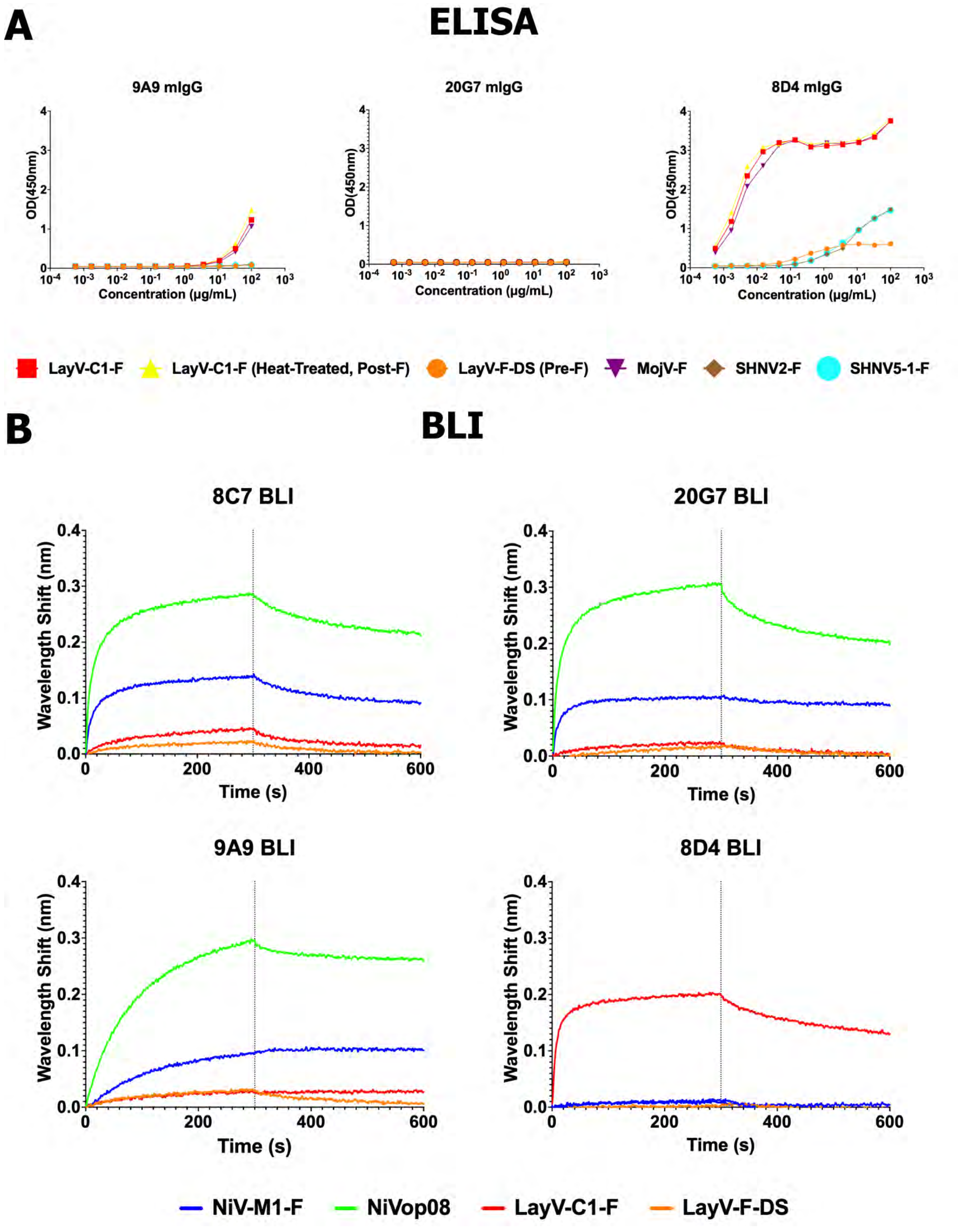


**Supplemental Figure 9: Measurement of Antibody Binding to HNV-F Proteins. (A)** ELISA-based binding of murine antibodies against select members of the HNV-F panel. **(B)** BLI-based binding of murine antibodies against select members of the HNV-F panel.

**C F Panel Binding to Antibodies (BLI)**

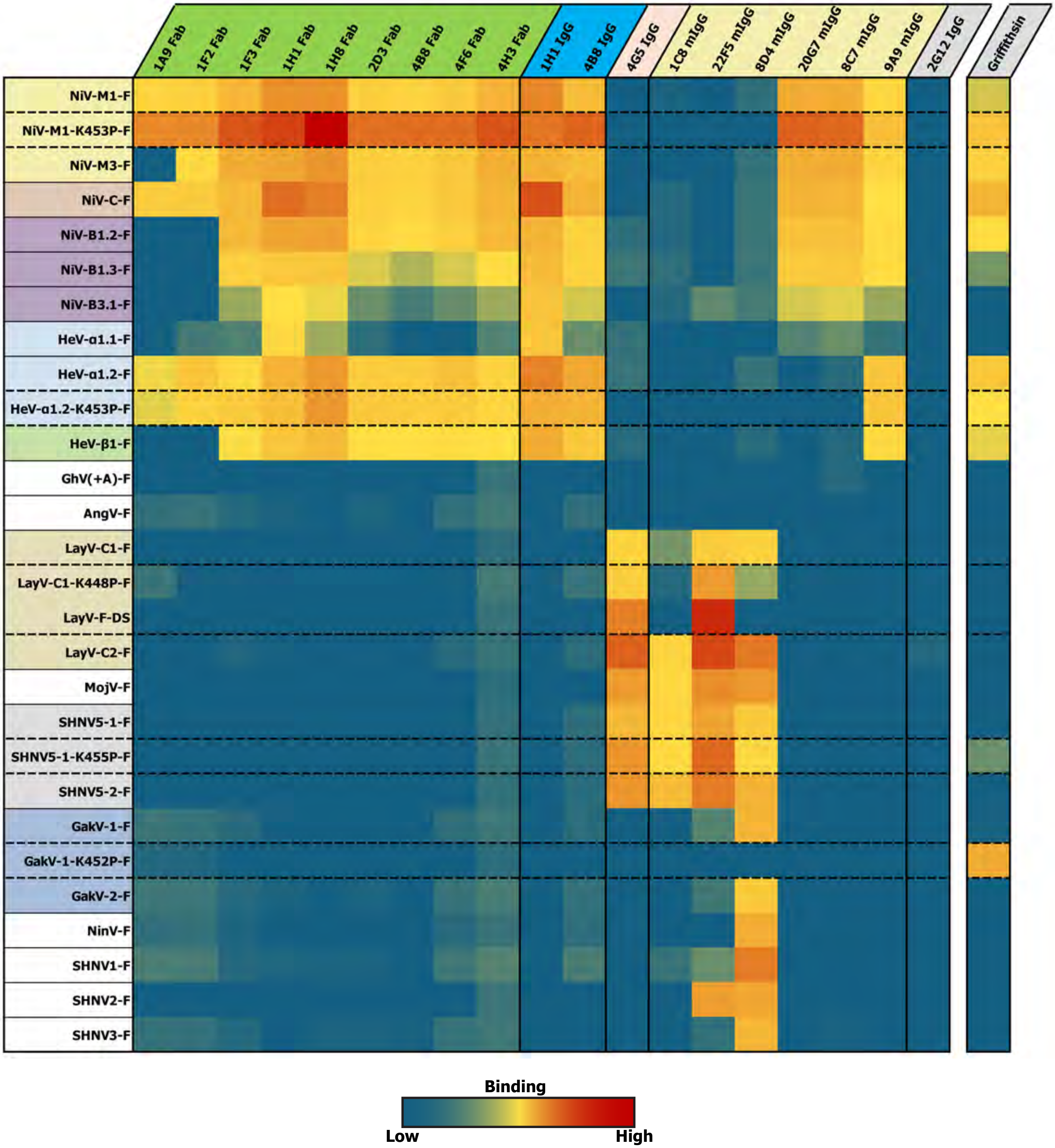


**Supplemental Figure 9 (continued): Measurement of Antibody Binding to HNV-F Proteins. (C)** BLI-based binding of F proteins to antibodies and Fabs. Heatmap represents maximum response value during association phase of BLI experiment (yellow: 0.5nm, red: 3.85nm). Colors for F proteins maintained from figure 1. Pre-fusion stabilized F constructs are signified by horizontal dashed lines. Antibodies are grouped and colored based on known binding specificity and type, green: Anti-NiV Fabs, blue: Anti-NiV IgGs, salmon: Anti-LayV IgG, tan: mIgG isolated during this study, gray: glycan reactive.

### A 22F5 Binding to LayV-F Ectodomains

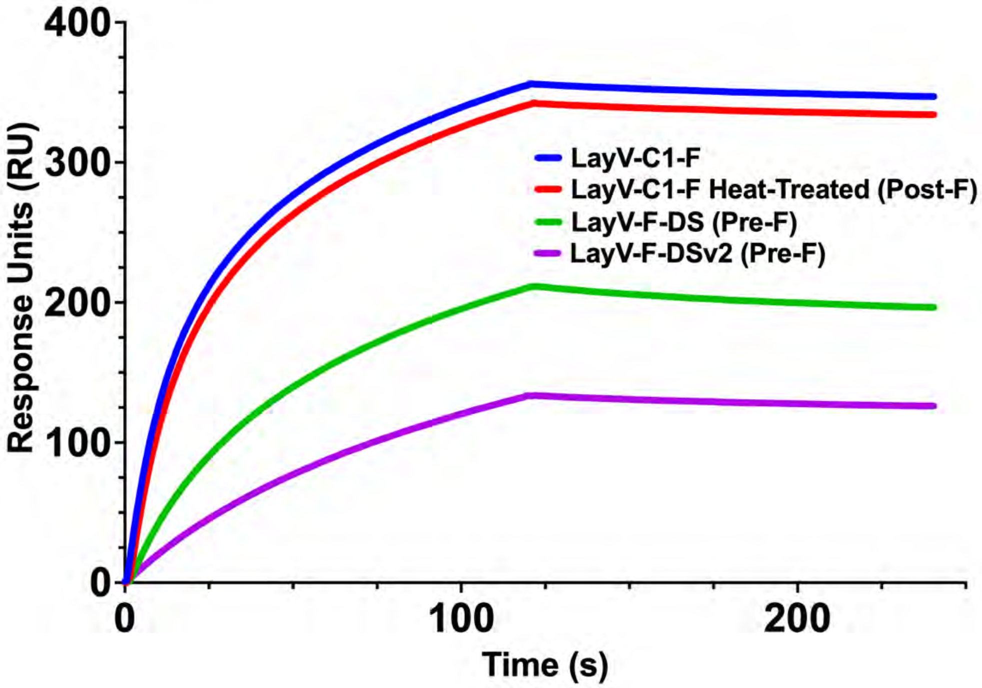


# B

**>22F5_Heavy_Chain:**

QLVQSGAEVKKPGASVKVSCKASGYTFTGYYMHWVRQAPGQGLEWMGWINPNNGGRNYVQ
KFQGRVTMTRDTSISTAYMELSSLSSDDTAVYYCARAGTSDWYFDLWGRGTLVTVSSAS

**>22F5_Light_Chain (>pK8_22F5.v2):**

DIVITQSPSSLAVSVGEKVTMSCKSSQSLLNSRTRRNYLAWYQQKPGQPPKLLIYWASTRESGV
PDRFTGSGSGTDFTLTISSVQAEDLAVYYCKQSYNSWTFGGGTKLEIKRT

**Supplemental Figure 10: SPR Analysis of 22F5 Binding to LayV-F Ectodomain. A)** Binding of LayV-F ectodomains to 22F5 IgG captured on a SPR sensor chip. The F ectodomains tested were wild-type LayV-C1-F, heat treated LayV-C1-F, and two pre-fusion stabilized versions, DS and DSv2. **B)** Sequence of 22F5 heavy and light chain variable regions.

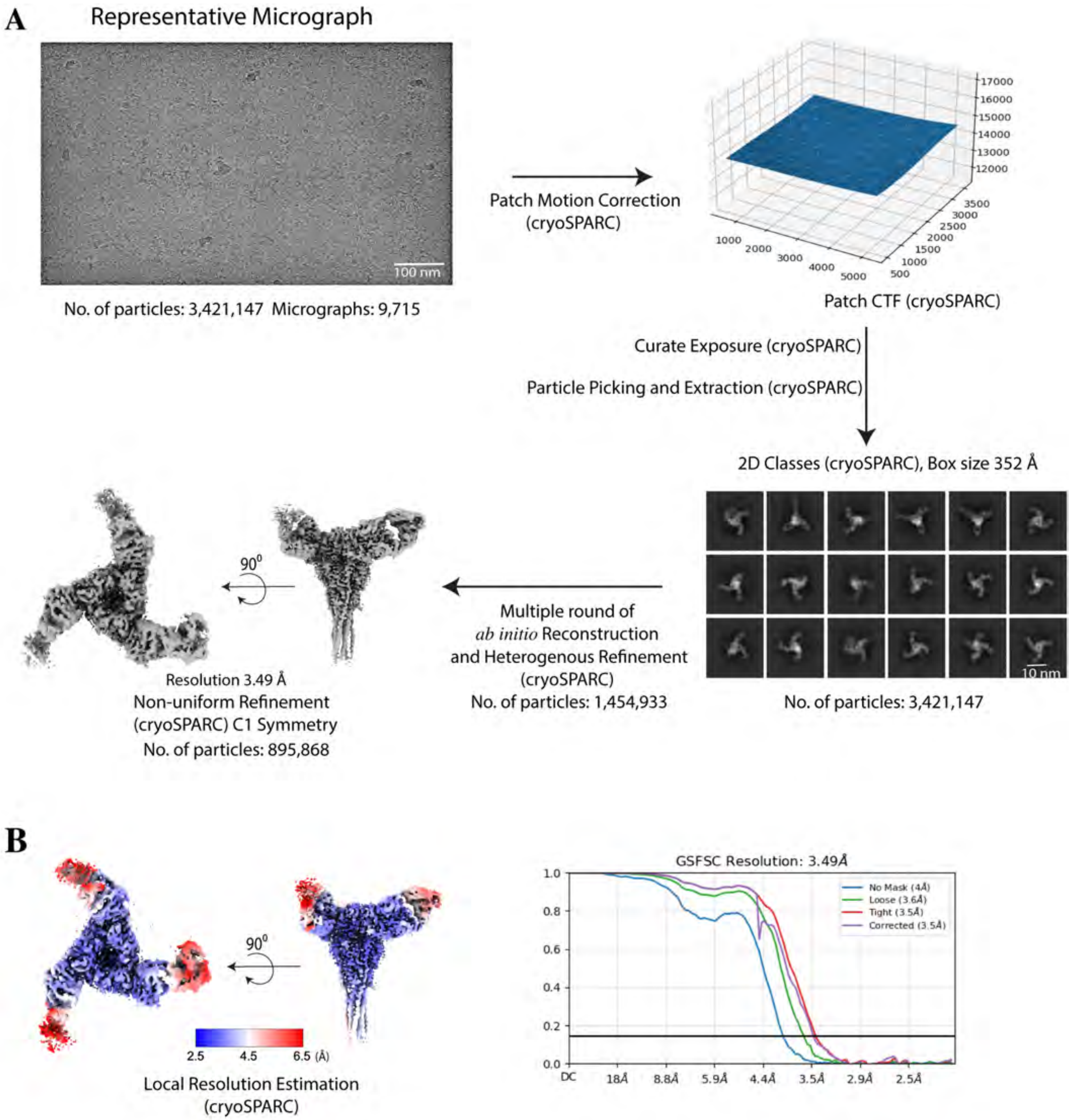


**Supplemental Figure 11: Cryo-EM Data Processing Workflow for LayV-C1-F Ectodomain and 22F5 Fab Complex.** **A)** Representative cryo-EM micrograph of the LayV-C1-F with 22F5 Fab complex after motion correction within cryoSPARC. Patch CTF estimation was performed, followed by exposure curation, particle picking (Blob Picker) and particle extraction. Multiple rounds of 2D classification were performed. A clean stack of particles was used for *ab initio* 3D reconstruction to generate the volume map. To refine further and improve the volume map, selected particles were processed multiple times through heterogeneous refinement and followed by final high resolution 3D reconstruction via non-uniform refinement. Side and top views of final refined map are shown. **B**) Local resolutions were estimated and represented in the color spectrum which indicates high resolution as blue and low resolution as red. The complex was refined using C1 symmetry. Fourier Shell Correlation (FSC) curve from the gold standard refinement of the final map is shown.

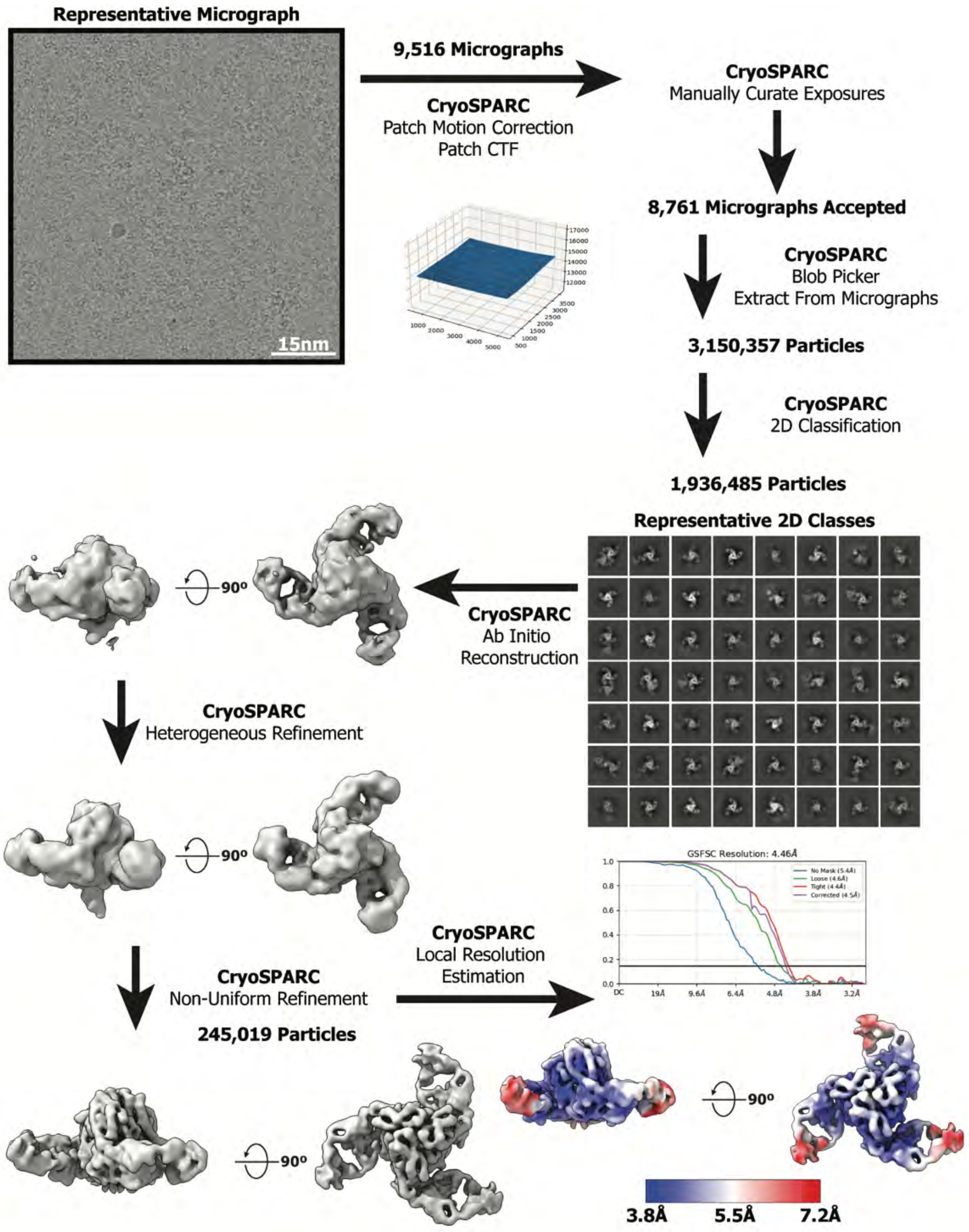


**Supplemental Figure 12: Cryo-EM Data Processing Workflow for LayV-F-DS Ectodomain and 22F5 Fab Complex.** Representative cryo-EM micrograph of the LayV-DS with 22F5 Fab complex after motion correction within cryoSPARC. Patch CTF estimation was performed, followed by exposure curation, particle picking (Blob Picker) and particle extraction. Multiple rounds of 2D classification were performed. A clean stack of particles was used for *ab initio* 3D reconstruction to generate the volume map. To refine further and improve the volume map, selected particles were processed multiple times through heterogeneous refinement and followed by final high resolution 3D reconstruction via non-uniform refinement. Side and top views of final refined map are shown. Local resolutions were estimated and represented in the color spectrum which indicates high resolution as blue and low resolution as red. The complex was refined using C3 symmetry. Fourier Shell Correlation (FSC) curve from the gold standard refinement of the final map is shown.

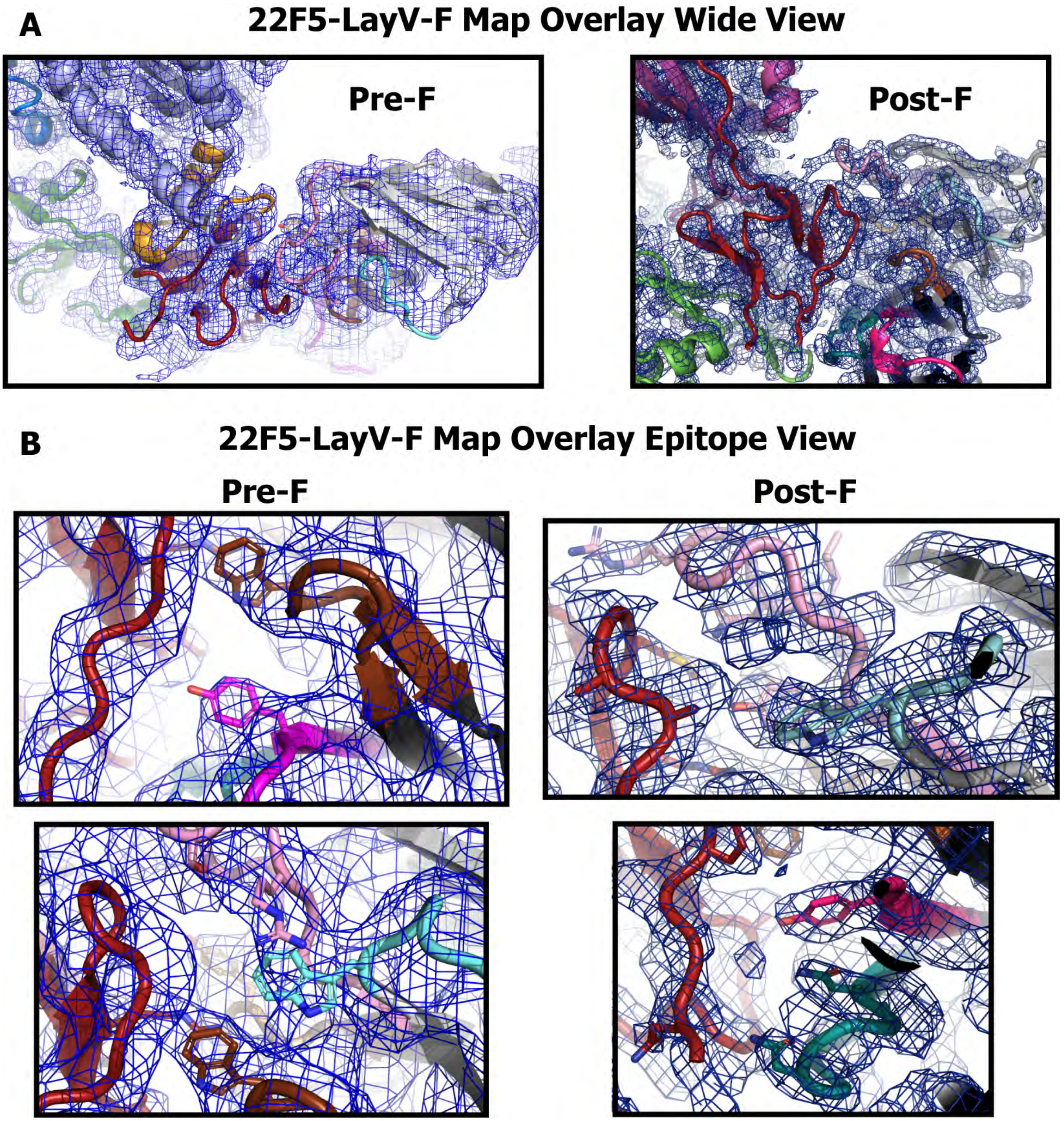


**Supplemental Figure 13: Model to map fit of 22F5-LayV-F structures. (A)** Wide views of the pre-F and post-F 22F5-bound structures, color scheme as in Figure 3 with blue mesh overlay of the electron potential map. **(B)** Epitope-specific views showing key interacting side chains illustrative of model fit.

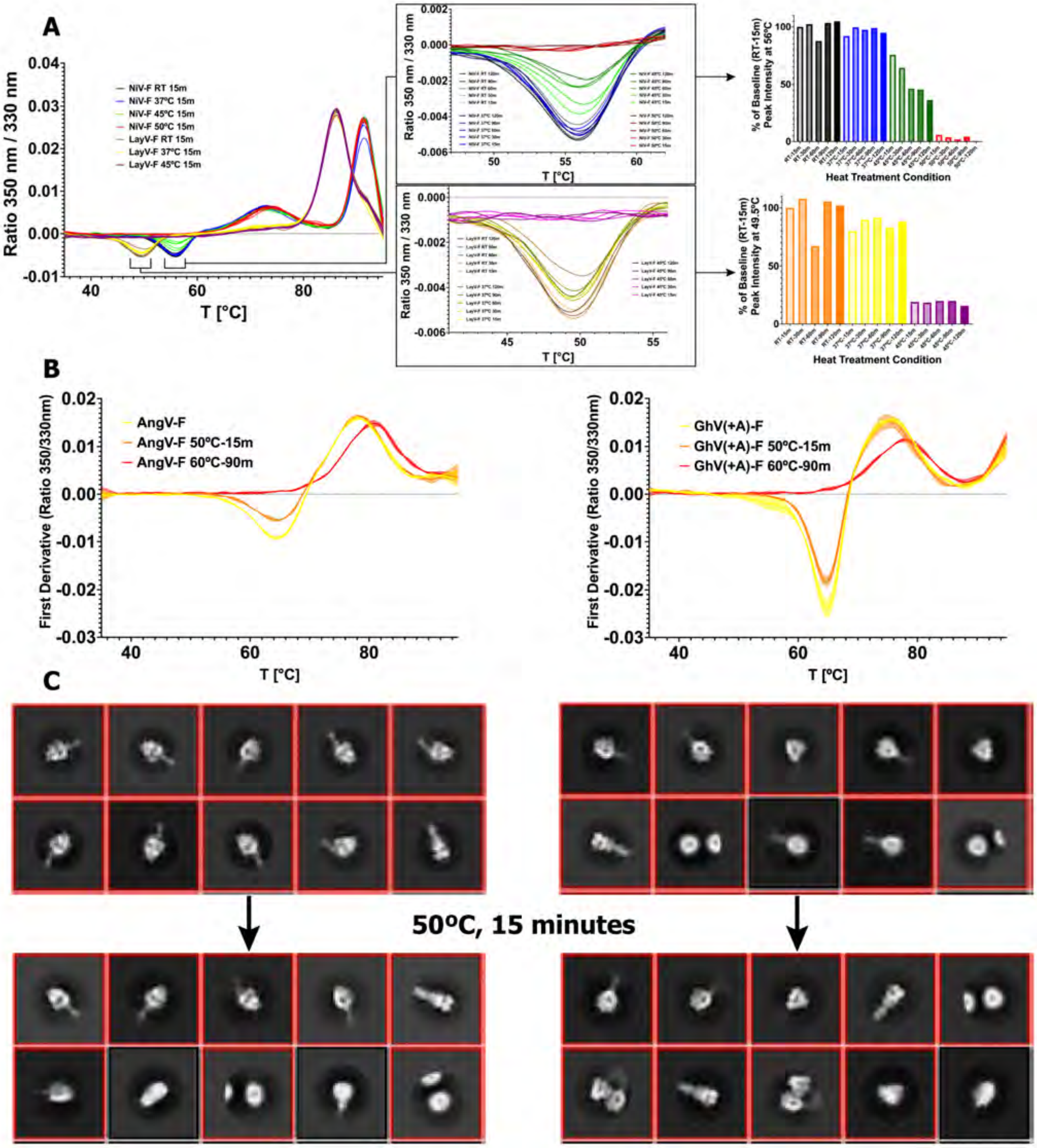


**Supplemental Figure 14: Heat Incubation of HNV-F Proteins. (A)** A series of heat incubation conditions applied to NiV-M1-F and LayV-C1-F. Left: Overlay of full DSF analysis of unfolding following 15-minute treatment with a series of temperatures. Middle: Zoomed-in view of the pre-fusion conversion peak with overlay of a series of time and temperature conditions. Right: A bar graph showing the intensity value of the peak relative to the room temperature, 15-minute condition. **(B)** Overlay of full DSF analysis of unfolding for AngV-F (left) and GhV(+A)-F (right) for several heat treatment conditions. **(C)** Representative 2D classes from NSEM analysis of AngV-F (left) and GhV(+A)-F (right) without treatment (top) and after 50ºC, 15-minute treatment (bottom).

#### F Protein W and Y Variability

#### G Head Domain W and Y Variability

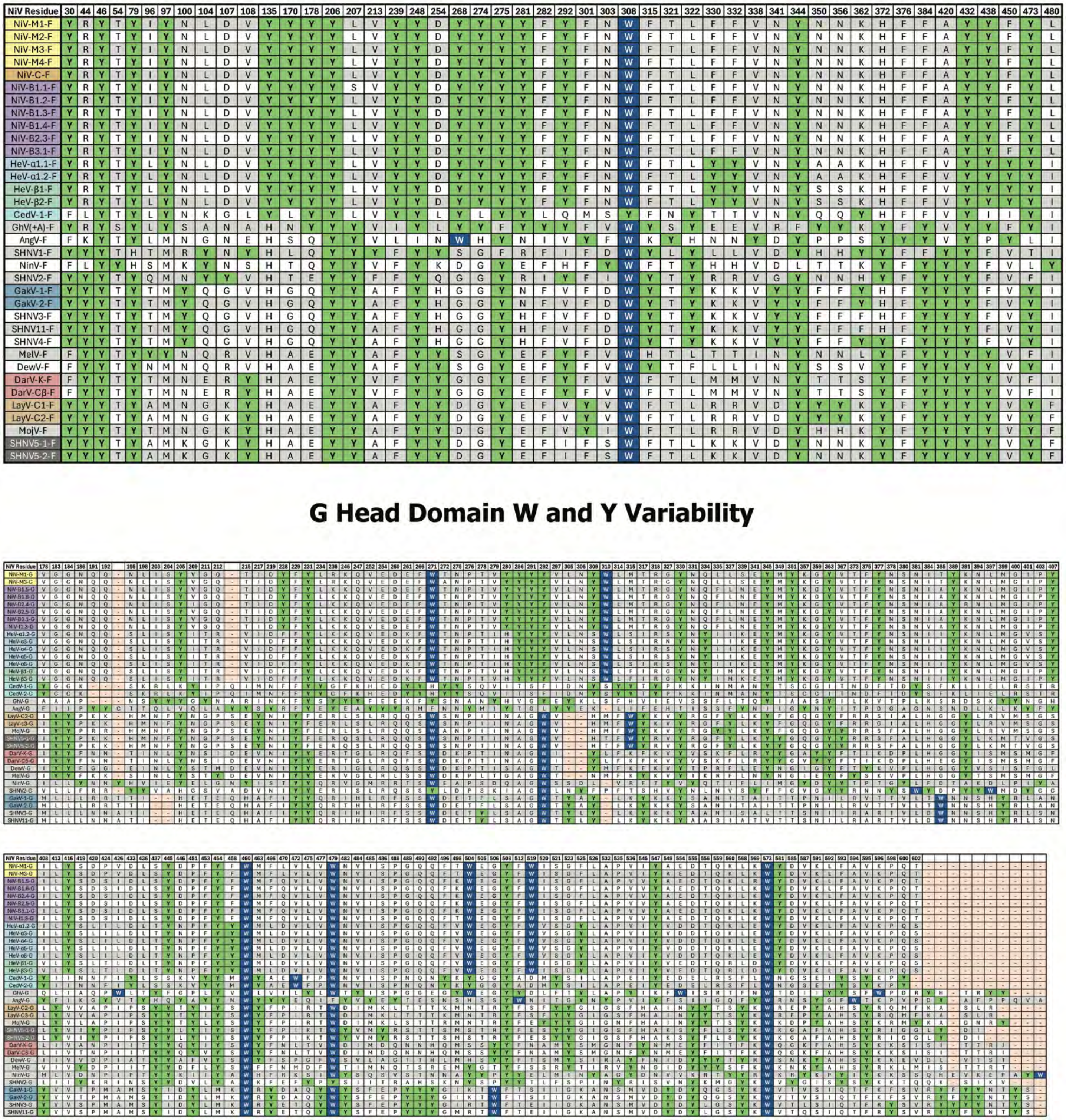

**Supplemental Figure 15: DSF-relevant Residue Variability.** Sequence alignment and highlighting of DSF-relevant tryptophan, blue, and tyrosine, green, residues. Gaps in the alignment are colored salmon. Species color scheme continued from Figure 1.

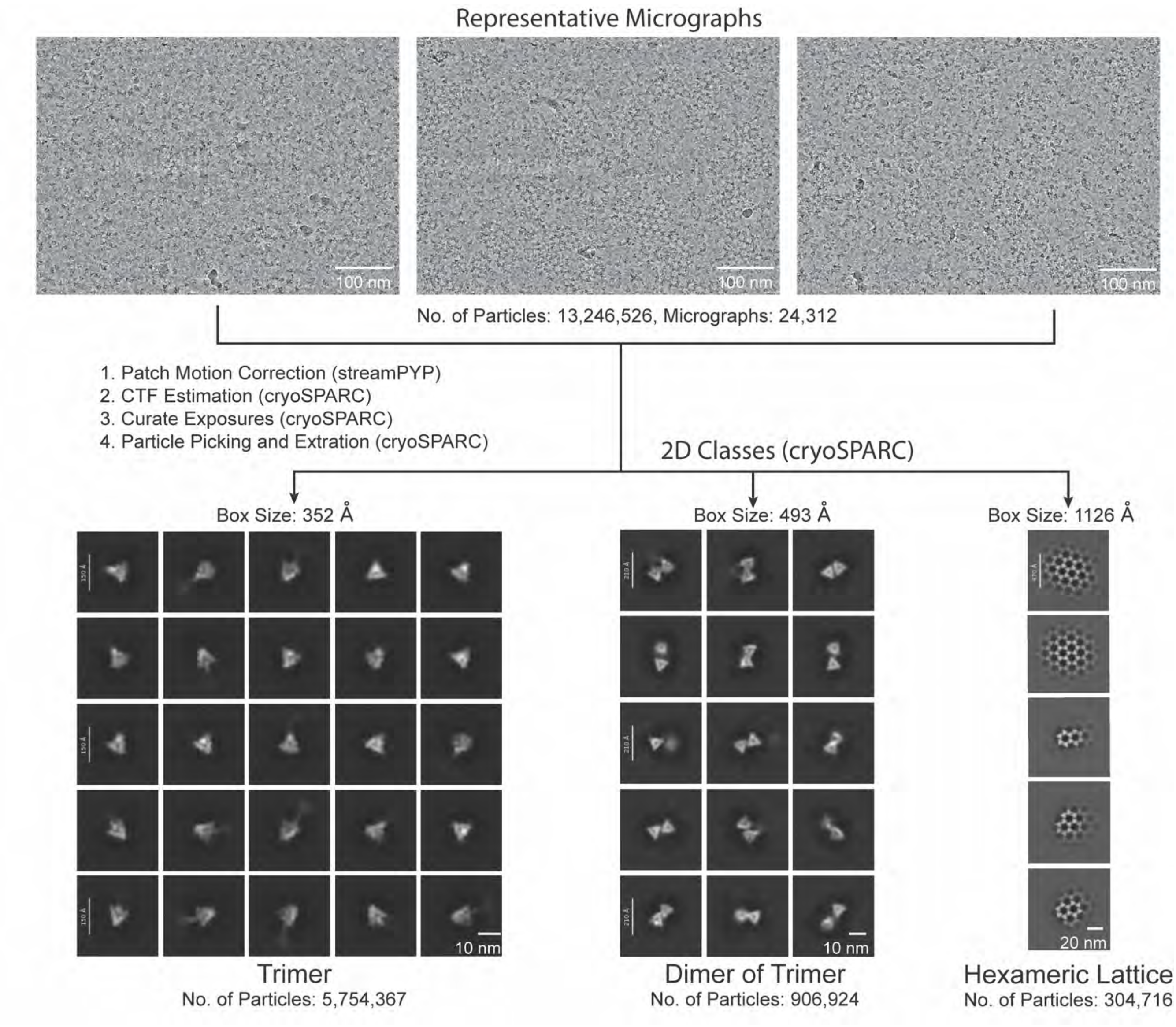


**Supplemental Figure 16: Workflow of Particle Extraction and 2D Classification.** Representative cryo-EM micrographs of the AngV-F protein after motion correction by nextPYP. Particles were picked and extracted after curating micrographs through CTF estimation. Different box sizes are utilized to accommodate the varying dimensions of specific molecular assemblies. Trimer particles are extracted using a box size of 320 pixels, while dimer of trimer particles require a larger box size of 512 pixels. For even larger molecular assemblies, such as hexameric lattices, a box size of 1024 pixels is employed. These extracted particles are then subjected to 2D classification to group and refine them based on their structural features, ensuring accurate downstream analysis (Supplemental Figures 11-12).

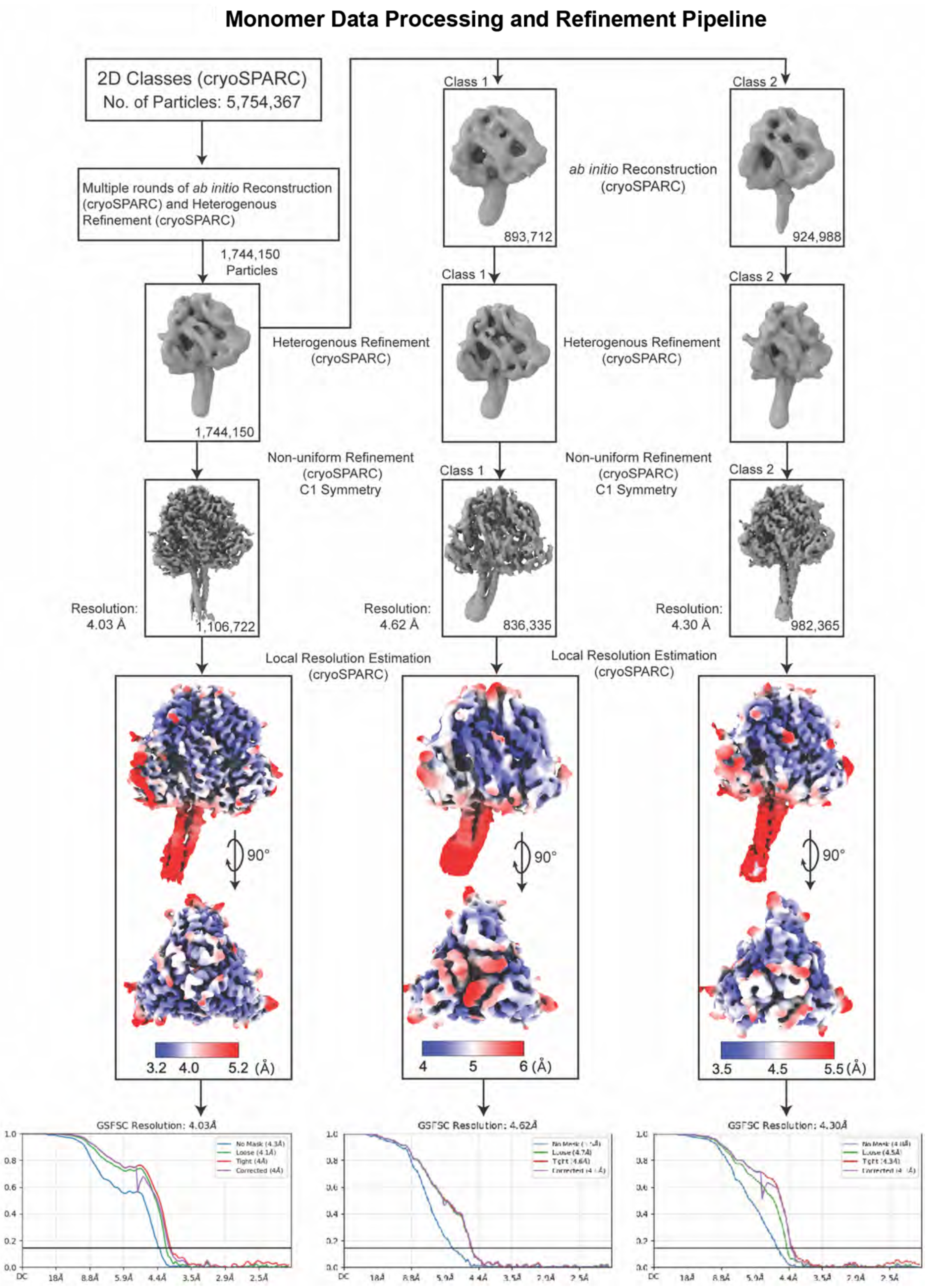


**Supplemental Figure 17: Cryo-EM Data Processing and Refinement Outline of Monomer Pre-fusion AngV-F Protein.** Representative 2D classes (Supplemental Figure 10) were selected for the *ab-intio* 3D reconstruction, with number of particles and software programs used in each step documented. Poor-quality particles were discarded after through heterogenous refinement and resulting volumes were processed using non-uniform refinement to obtain a high-resolution 3D reconstruction. Local resolutions were estimated and plotted on the side and top views of the AngV-F protein. Resolution values are shown as a blue to red spectrum for high-resolution regions (~3-4 Å) blue, and low-resolution regions (~5–6 Å) red. Structures were refined using C1 symmetry, with gold-standard Fourier shell correlation curves displayed at the bottom.

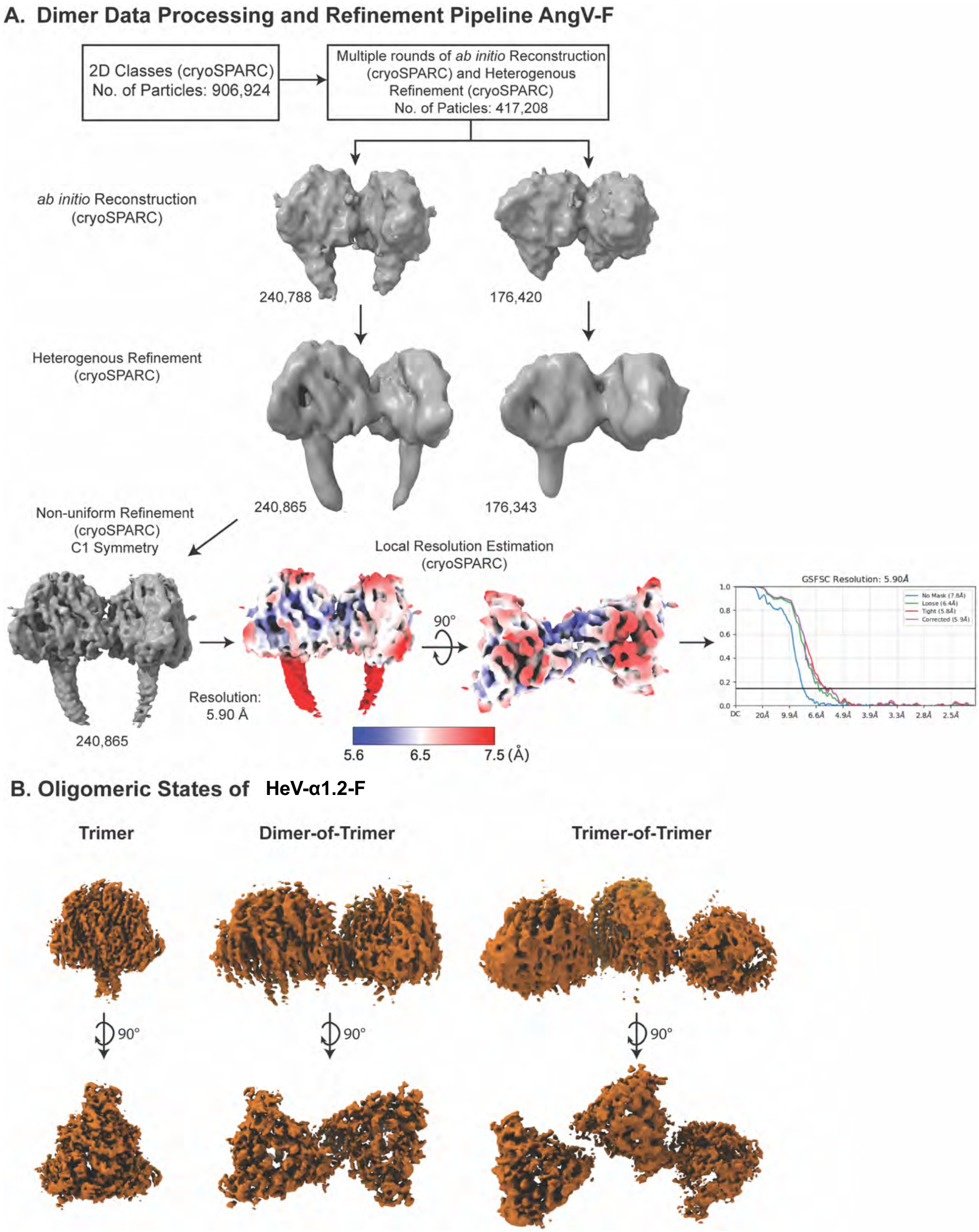


**Supplemental Figure 18: Oligomeric States of AngV-F and HeV-α1.2-F. (A)** Cryo-EM Data Processing and Refinement Outline of Dimer pre-fusion AngV-F protein. Representative 2D classes (Supplemental Figure 10) were selected for the *ab-intio* 3D reconstruction, with number of particles and software programs used in each step documented. Poor-quality particles were discarded after heterogenous refinement and resulting volume was processed with non-uniform refinement to build a high-resolution 3D reconstruction. Local resolutions were estimated and plotted on the side and top views of the AngV-F Dimer. Resolution values are shown as a blue to red spectrum for high-resolution regions (~5-6 Å) blue, and low-resolution regions (~7.5 Å) red. Structures were refined using C1 symmetry, with gold-standard Fourier shell correlation curves displayed at the bottom. **(B)** Cryo-EM HeV-α2-F ectodomain. The non-uniform refinement reconstruction (cryosparc) of trimer (left), dimer-of-trimers (middle) and trimer-of-trimers (right) are shown here.

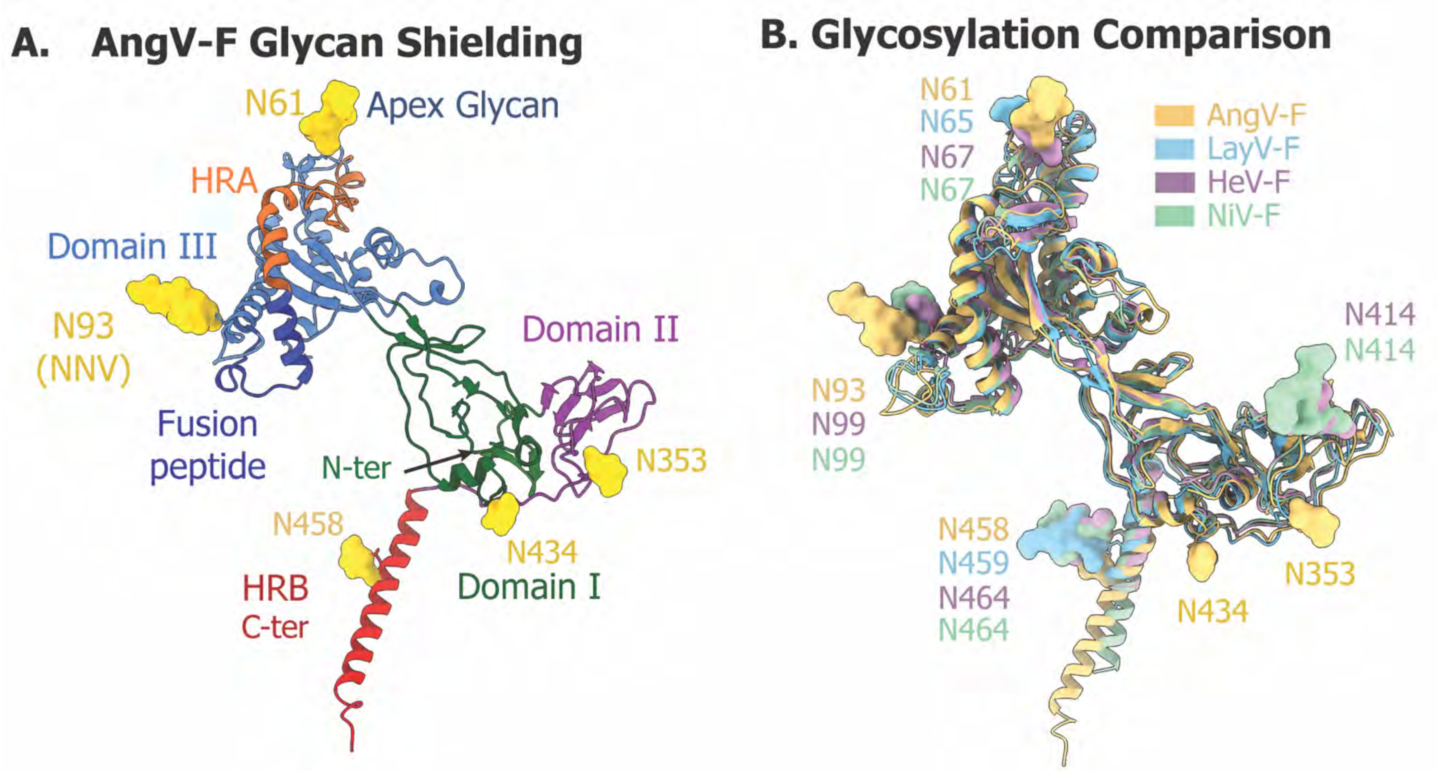


**Supplemental Figure 19: AngV-F Glycan Distribution and Differences Compared to other HNV-F Proteins**. **(A)** Cartoon representation of an AngV-F protomer in the prefusion state, colored and annotated to highlight respective domains by functionality and glycans (yellow). **(B)** Comparison of glycans among the F-proteins: AngV-F (yellow, from this study), LayV-F (blue, PDB: 8FMX), HeV-F (violet, PDB: 7KI6), and NiV-F (green, PDB: 5EVM).

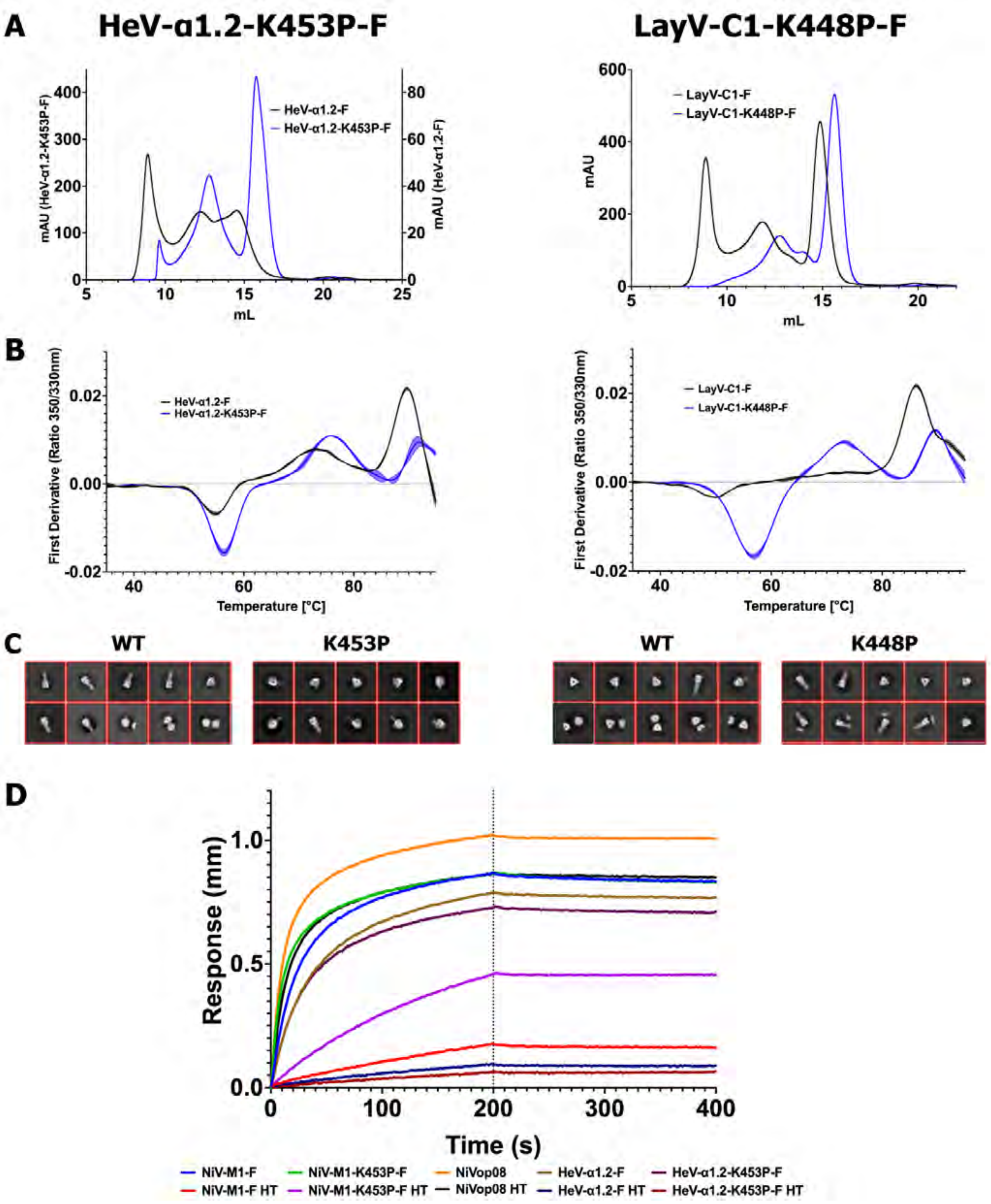


**Supplemental Figure 20: Characterization of HRB Start Proline HNV-F Mutants**. **(A)** SEC overlay of HeV-α1.2-F WT and mutant (left) and LayV-C1-F WT and mutant (right). For HeV-α1.2-F, WT is plotted on the right axis and mutant on the left, with axes scaled to 1L expression. **(B)** DSF unfolding analysis profile, shown with the center line as an average of three replicates and lines above and below with shading in the middle to represent standard deviation. Left and right species as in **A.** **(C)** Representative 2D classes from NSEM analysis of WT and mutant constructs, left and right species as in **A.** **(D)** BLI analysis of binding between 4B8 anti-NiV-F antibody and a series of HNV-F ectodomain constructs.

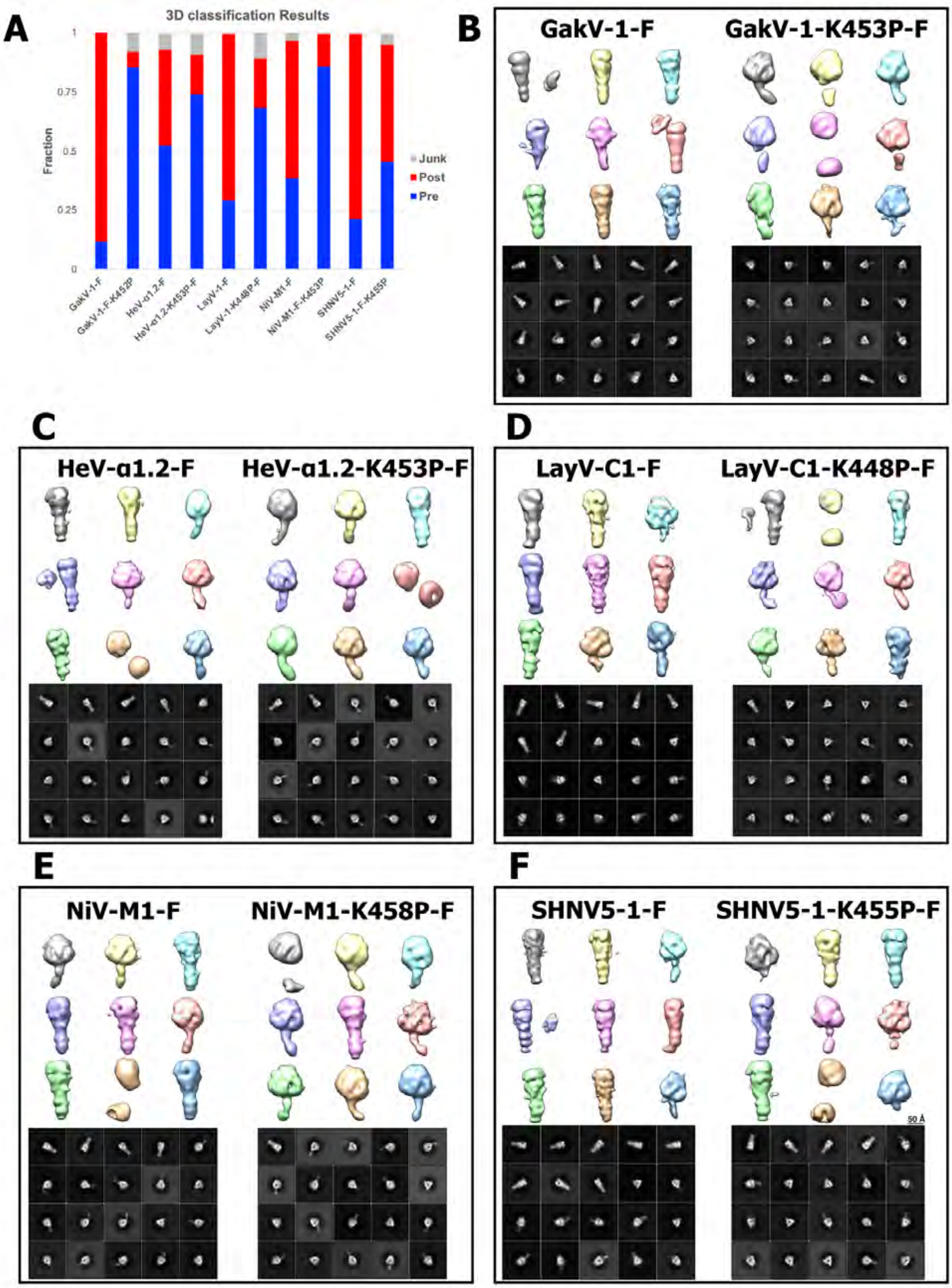


**Supplemental Figure 21: NSEM 3D Reconstruction of HRB Start Proline Mutants. (A)** Bar graph representation of pre-fusion, post-fusion, and junk particles in each sample's dataset. **(B-F)** Comparison of wild type (left) and mutant (right) classes' representative 3D classes and 2D classes. **(B)** GakV-1-F. **(C)** HeV-α1.2-F. **(D)** LayV-C1-F. **(E)** NiV-M1-F. **(F)** SHNV5-1-F.

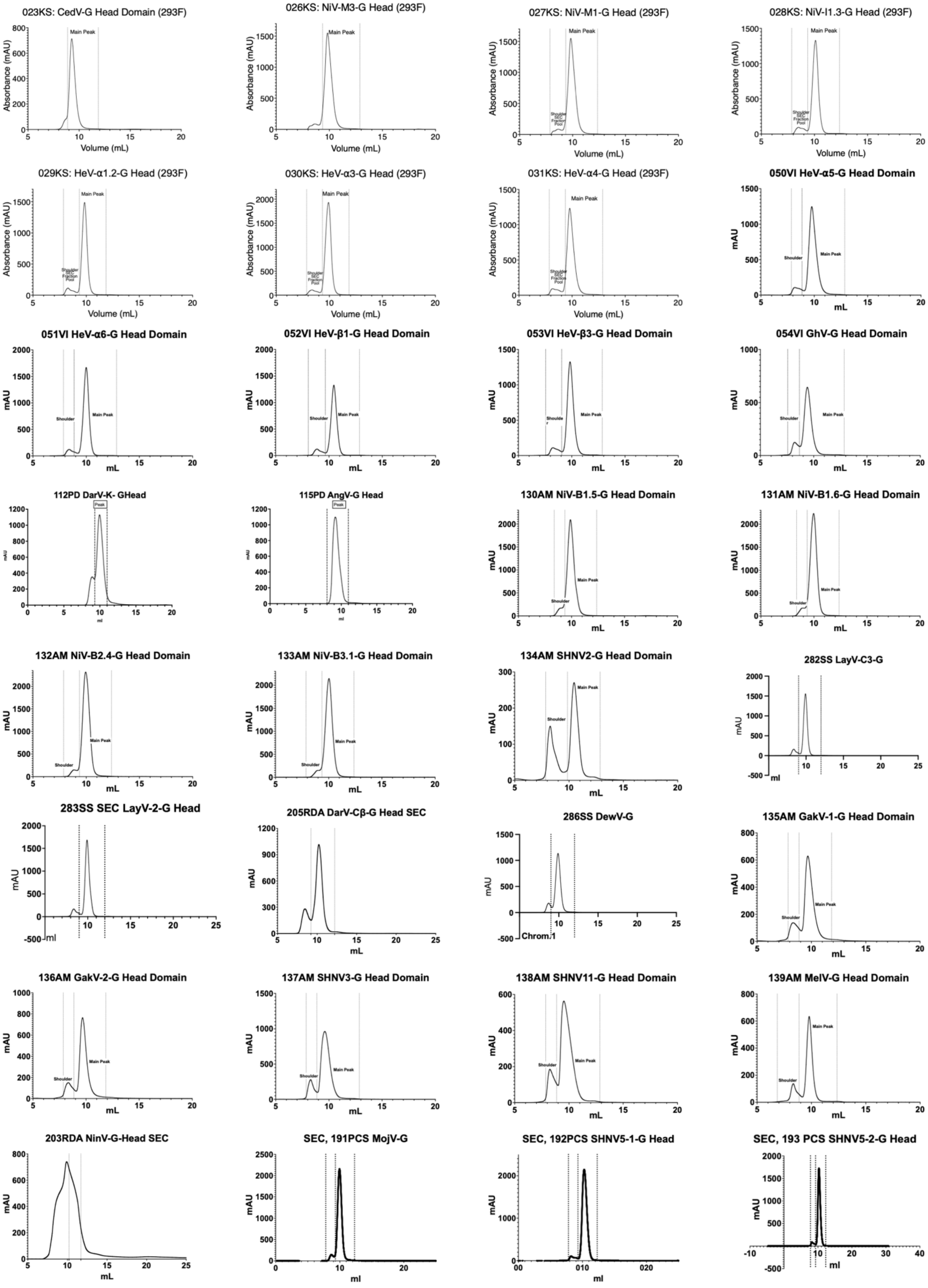


**Supplemental Figure 22: Size Exclusion Chromatography (SEC) Profiles of HNV-G Head Domains.** Fraction boundaries of collected peaks are labelled. Internal production lot numbers included with strain names.

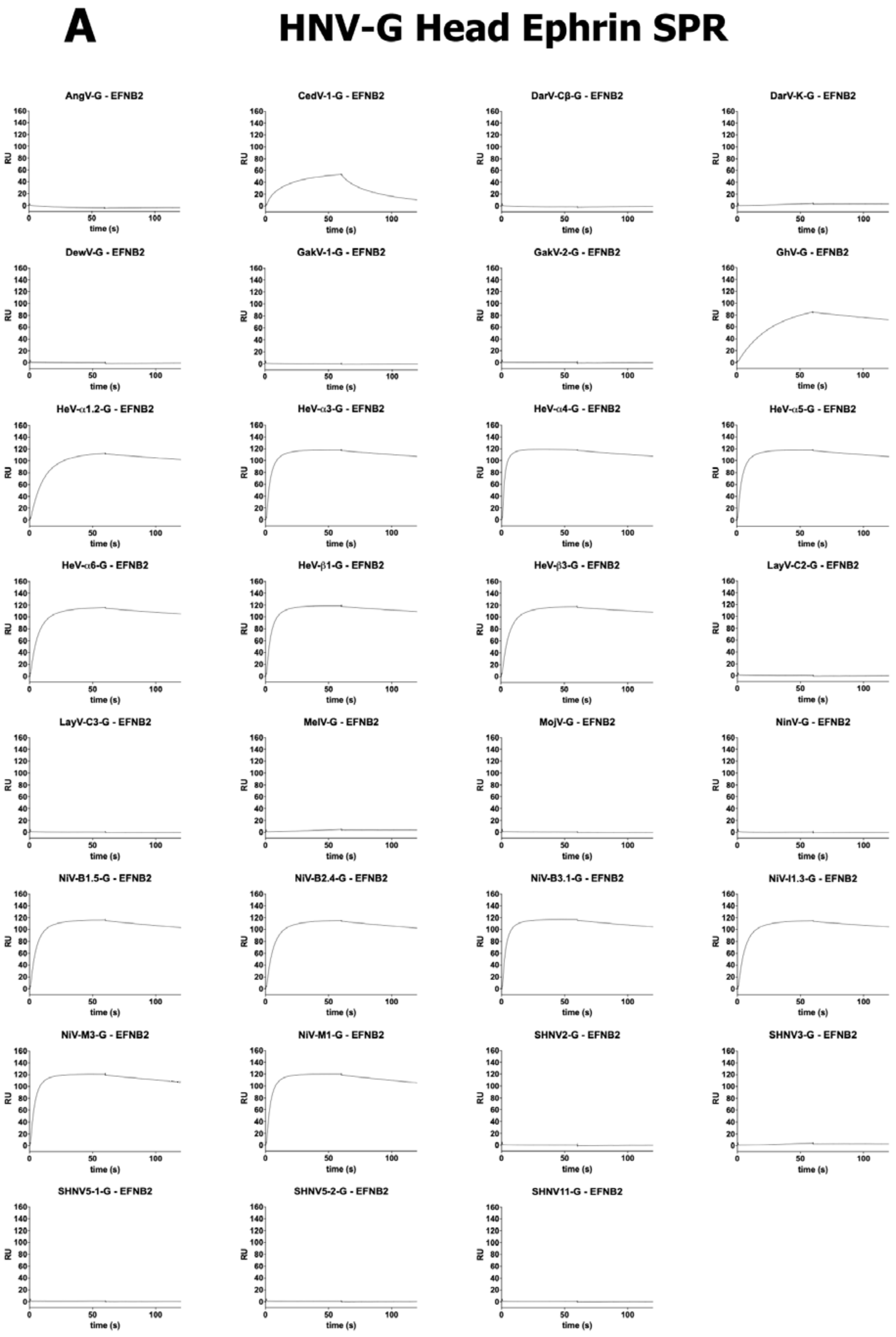


**Supplemental Figure 23: HNV-G Binding to Ephrin and Antibodies. (A)** Binding curves generated through SPR for recombinant Fc-Ephrin-B2 and B3 binding to members of the G head domain panel

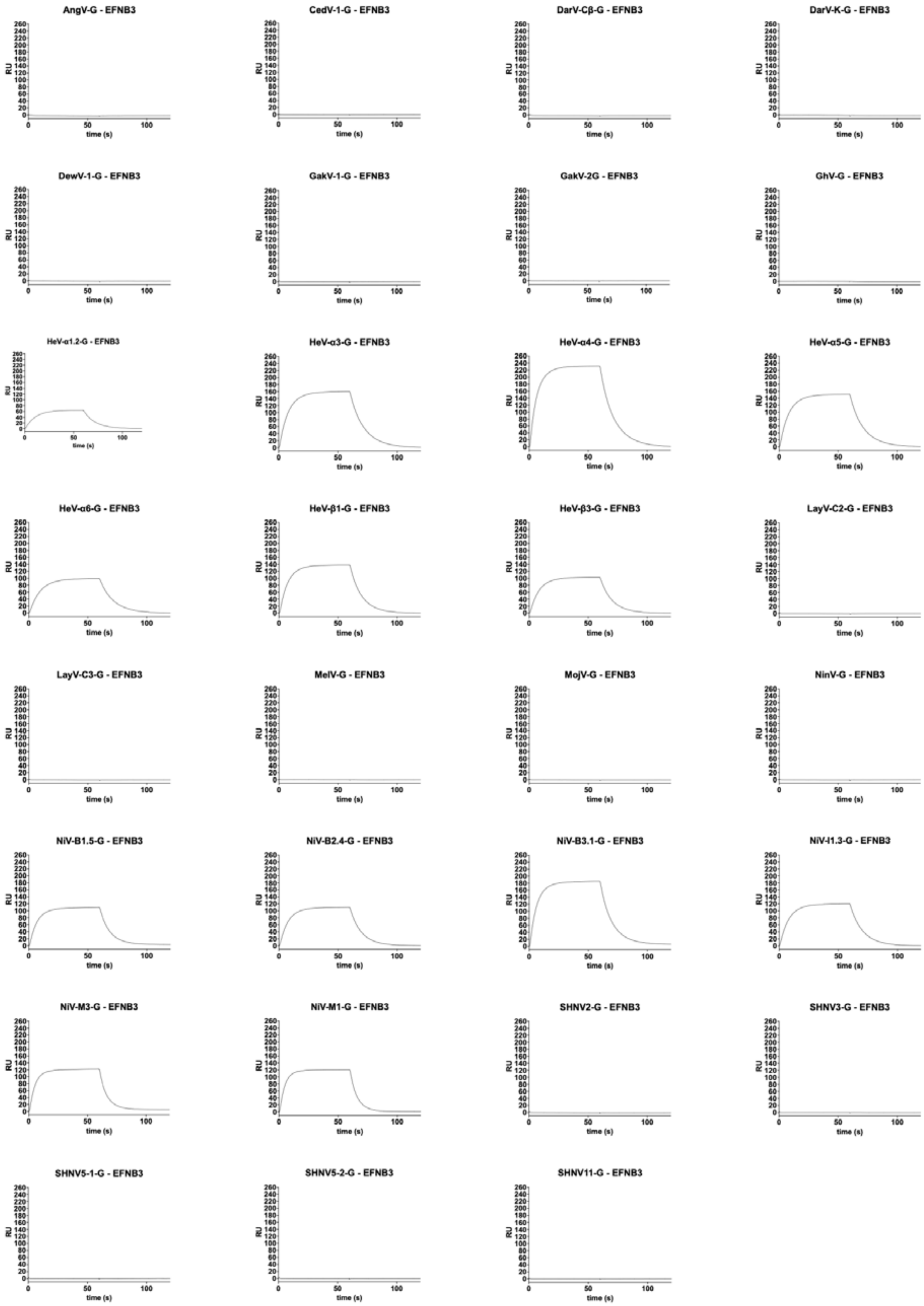


**Supplemental Figure 23 (continued): HNV-G Binding to Ephrin and Antibodies.**

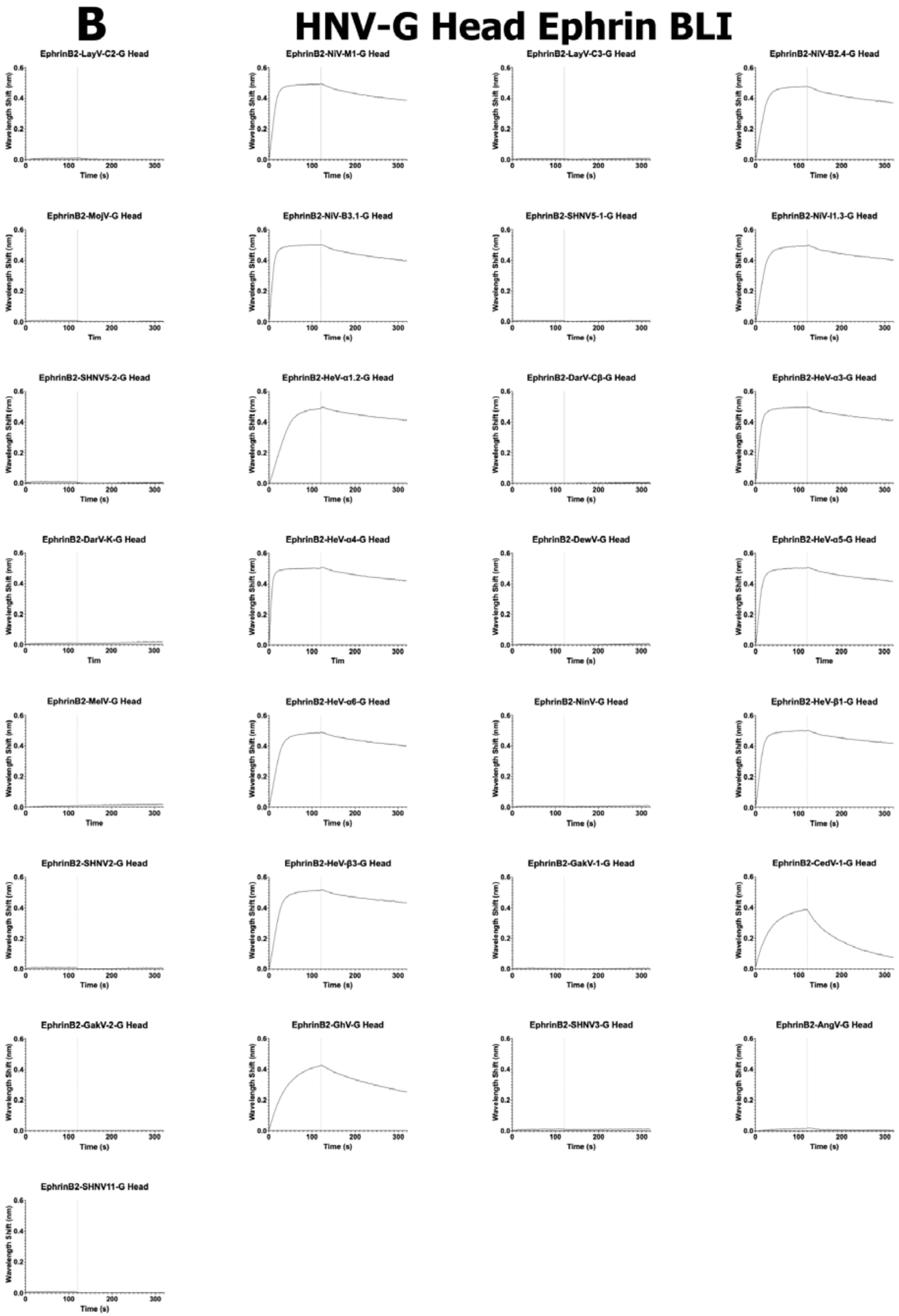


**Supplemental Figure 23 (continued): HNV-G Binding to Ephrin. (B)** Binding curves generated through BLI for recombinant Fc-Ephrin-B2 binding to members of the G head domain panel.

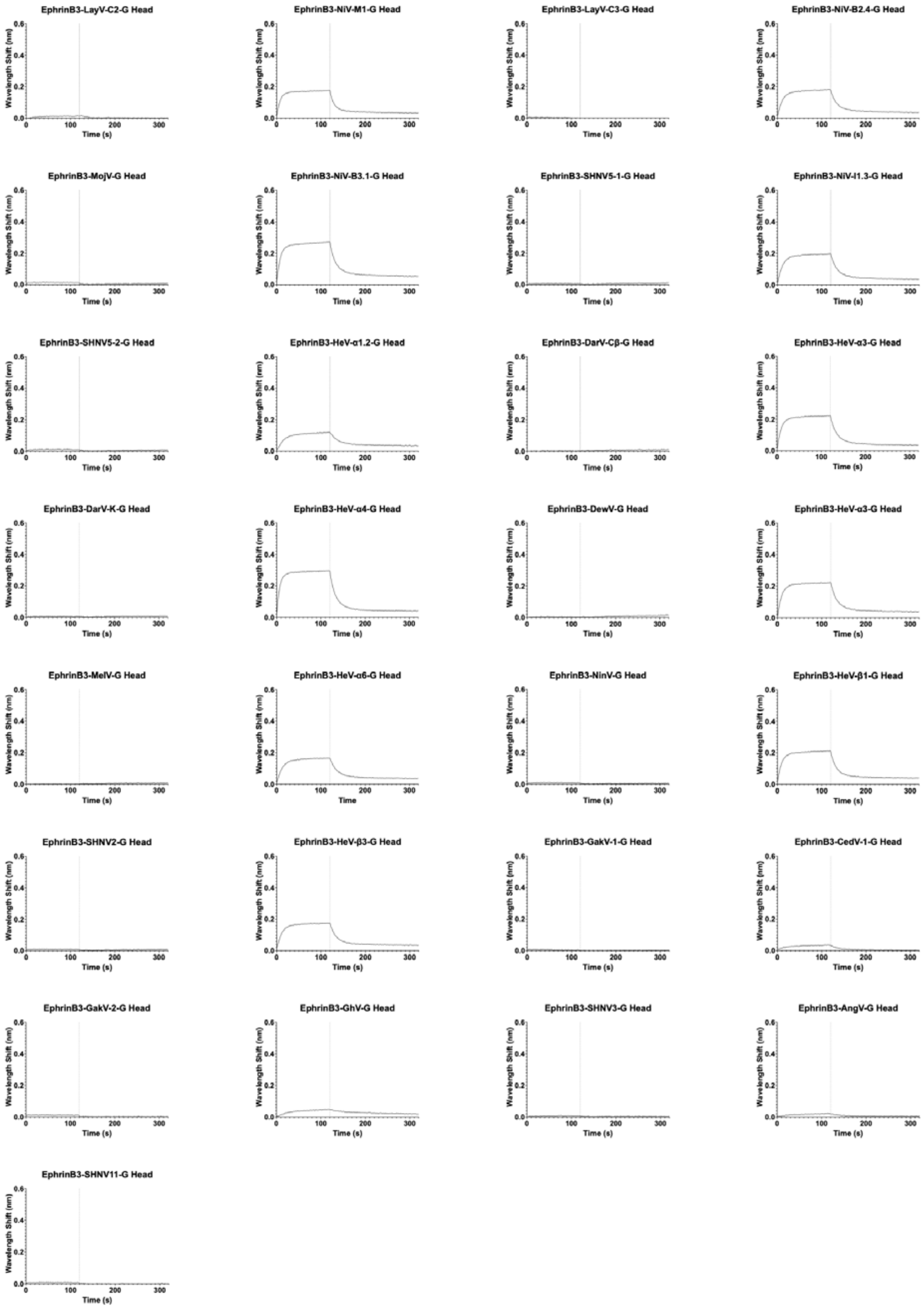


**Supplemental Figure 23 (continued): HNV-G Binding to Ephrin. (B)** Binding curves generated through BLI for recombinant Fc-Ephrin-B3 binding to members of the G head domain panel.

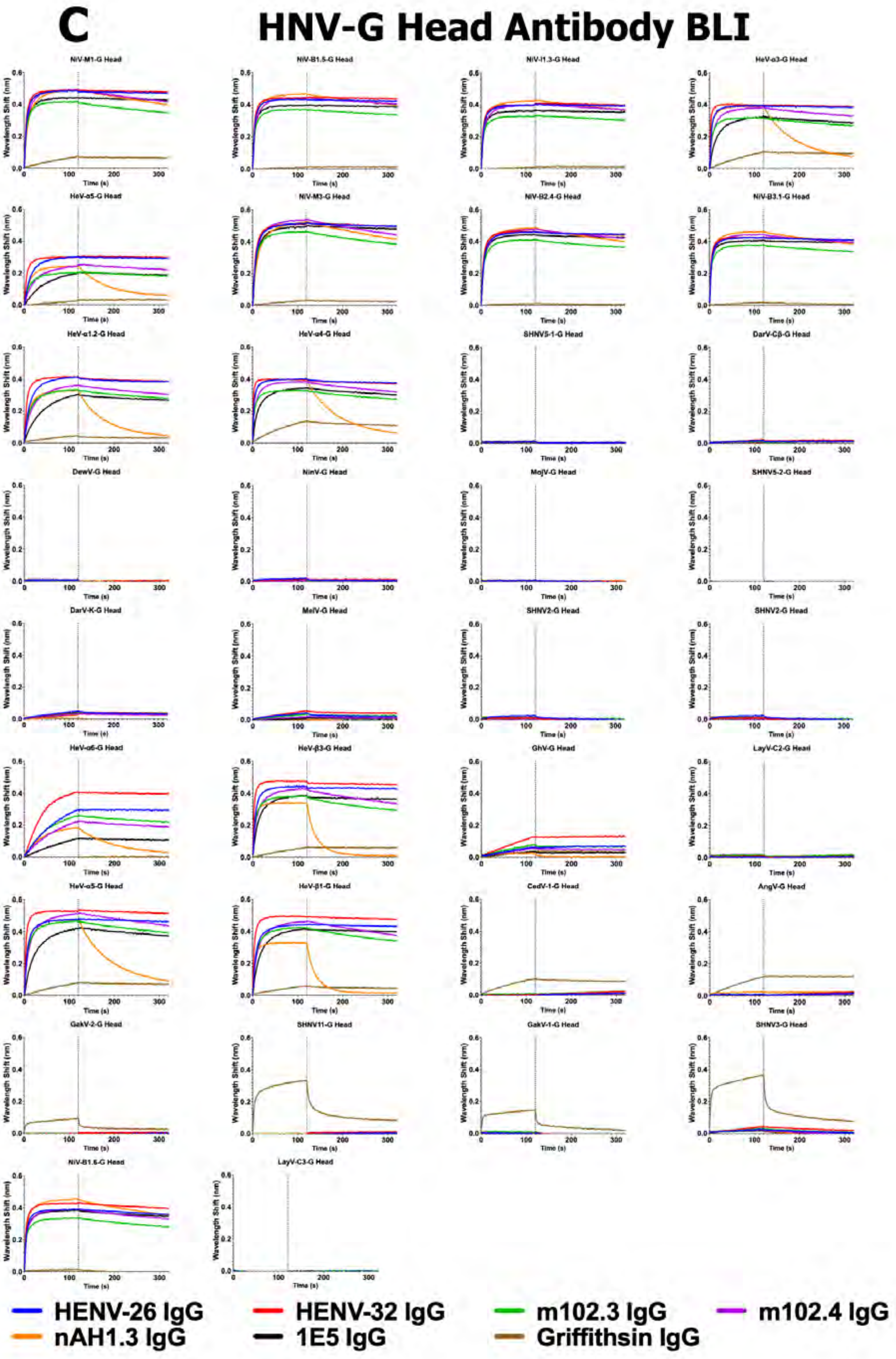


**Supplemental Figure 23 (continued): HNV-G Binding to Ephrin. (C)** Binding curves generated through BLI for antibody binding to members of the G head domain panel, grouped by G head strain.

### D HNV-G Head Antibody ELISA

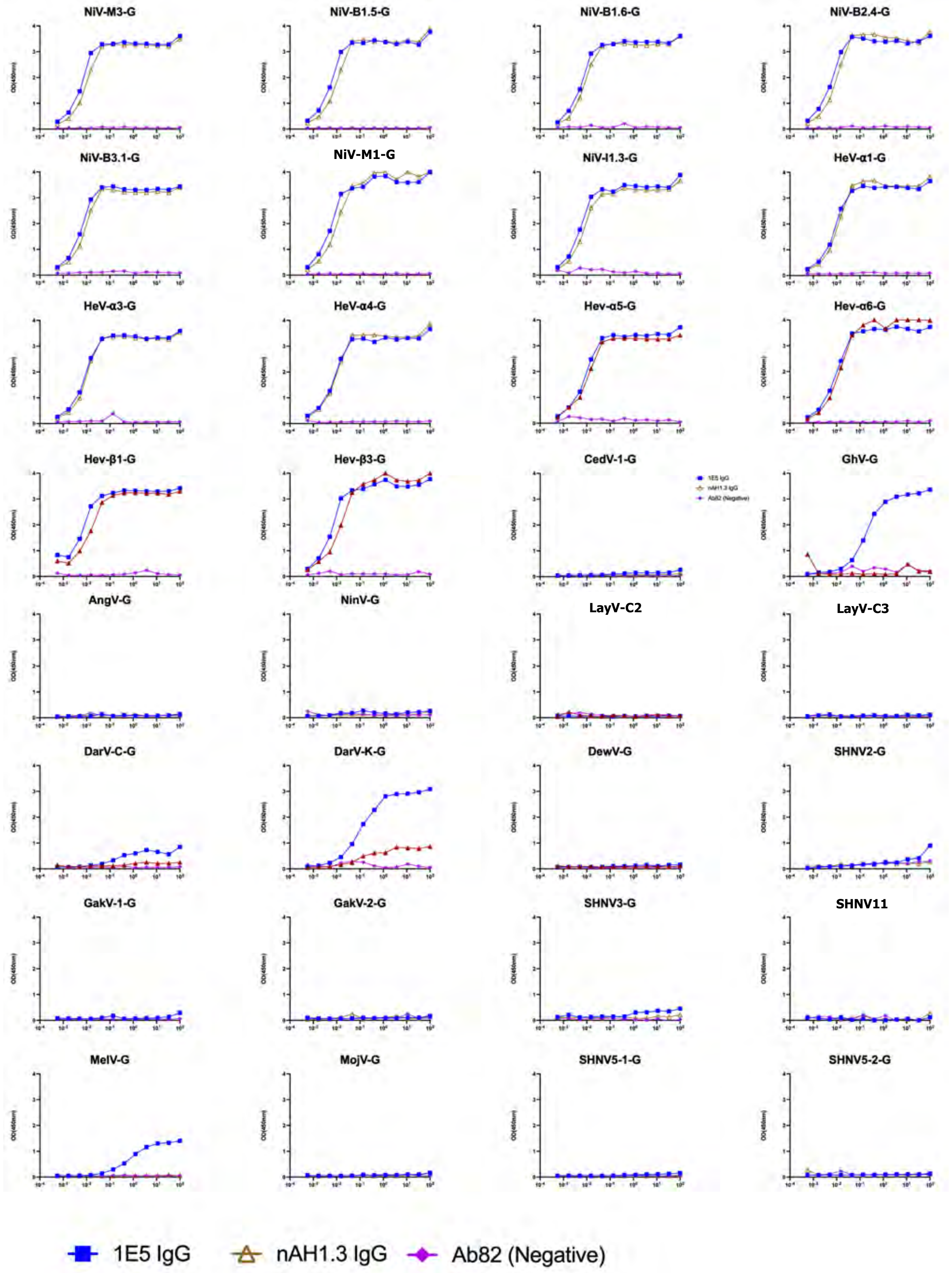


**Supplemental Figure 23 (continued): HNV-G Binding to Ephrin and Antibodies. (D)** Binding curves generated through ELISA for several anti-G antibodies binding to members of the G head domain panel, grouped by G head strain.

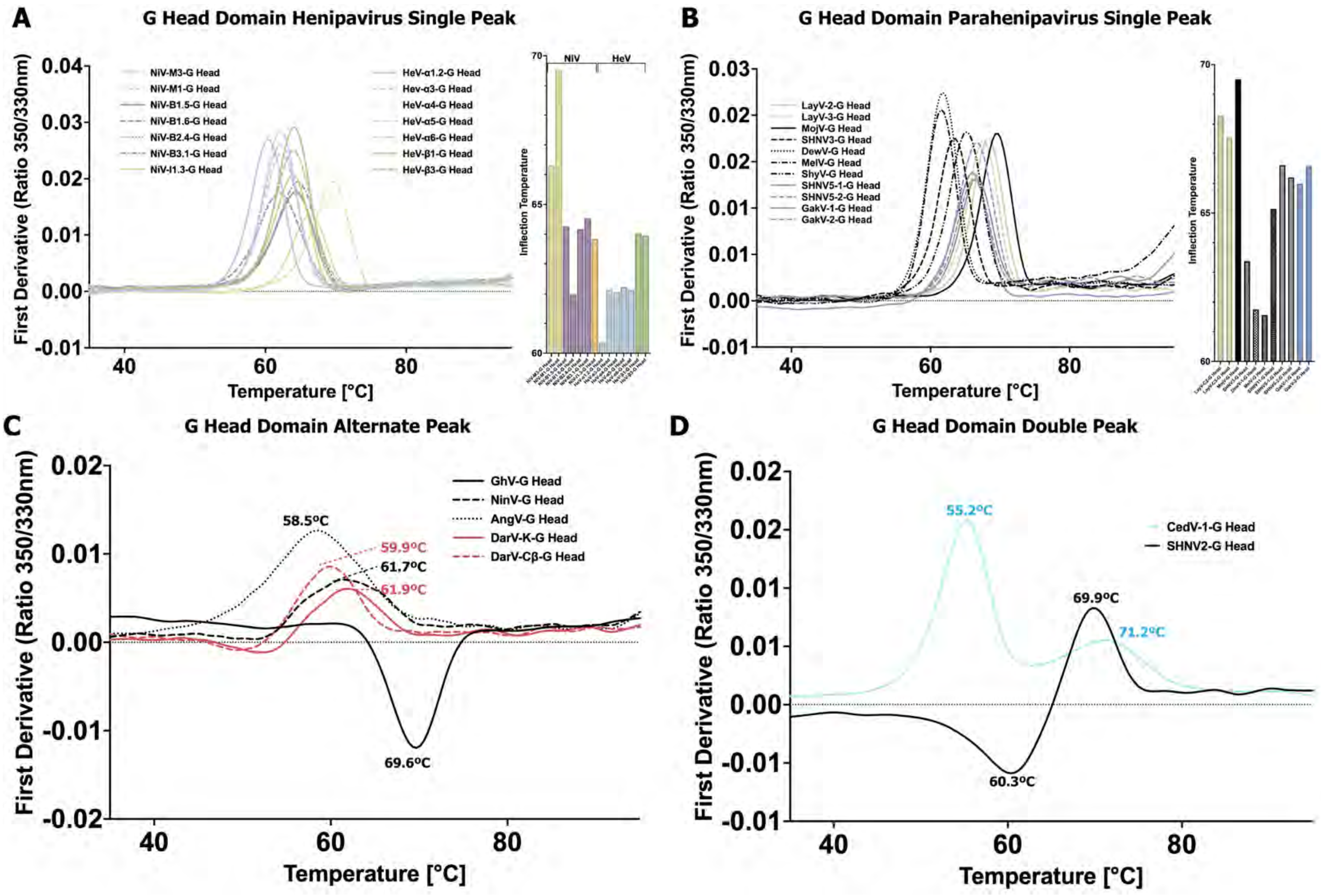


**Supplemental Figure 24: Thermostability of HNV G Head Domains. (A-D)** DSF analysis of G head domains, reporting the first derivative of 350/330nm ratio as a function of temperature. The panel's results are split based on their phylogeny and general DSF profile: **(A)** *Henipavirus* proteins with a single DSF peak; **(B)** *Parahenipavirus* proteins with a single DSF peak; **(C)** proteins with alternative peak shapes (broad or negative); **(D)** proteins with double peaks. In **(A-B)**, a side plot of inflection temperatures is shown for each species in the associated graph. In **(C-D)**, inflection temperatures are drawn directly on the plot.

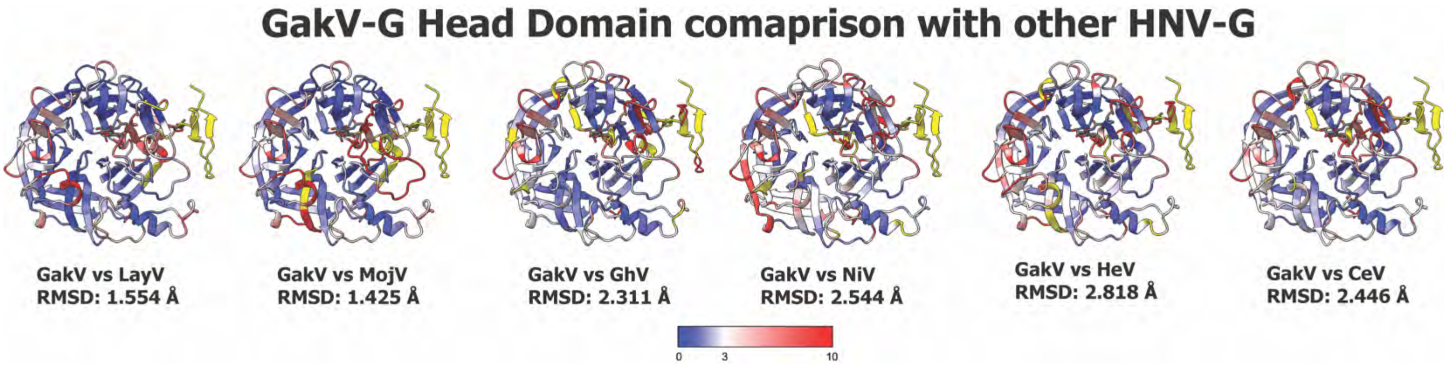


**Supplemental Figure 25: Head Domain Comparison.** Overlay of GakV-G head domain on the G head domains from other HNV species. PDB Codes: LayV-G, 8K80 (Unpublished); MojV-G, 5NOP [52]; GhV-G, 4UF7 [80]; NiV-G, 3D11 [81]; HeV-G, 6VY4 [16]; CedV-G, 6P72 [82].

**Supplemental Table 2. Cryo-Em Data Collection and Refinement Statistics.**

| | **AngV-F Trimer** | **AngV-F Dimer-of-Trimer** | **LayV-1-F with 22F5** | **LayV-F-DS with 22F5** |
|---|---|---|---|---|
| **PDB ID** | 9MNH | 9MQN | 9NZ0 | 9NZ2 |
| **EMDB ID** | EMD-48423 | EMD-48535 | EMD-48715 | EMD-49948 |
| **Data collection and processing** | | | | |
| Microscope | FEI Titan Krios | FEI Titan Krios | FEI Titan Krios | Tundra TEM |
| Detector | Gatan K3 | Gatan K3 | Gatan K3 | Falcon 4i |
| Magnification | 81000 | 81000 | 81000 | 180000 |
| Voltage (kv) | 300 | 300 | 300 | 100 |
| Electron exposure (e-/å^2) | 65 | 65 | 59.6 | 30.72 |
| DEFOCUS RANGE (µm) | -2.8 to -1.5 | -2.8 to -1.5 | -2.8 to -1.2 | -2.5 to -0.7 |
| Pixel size (å) | 1.1 | 1.1 | 1.1 | 1.5 |
| Reconstruction software | cryoSPARC | cryoSPARC | cryoSPARC | cryoSPARC |
| Symmetry imposed | C1 | C1 | C1 | C3 |
| Initial particle images (no.) | 5,754,367 | 906,924 | 3,421,147 | 3,150,357 |
| Final particle images (no.) | 1,106,722 | 240,865 | 895,868 | 245,019 |
| Map resolution (å) | 4.03 | 5.9 | 3.49 | 4.46 |
| Fsc threshold | 0.143 | 0.143 | 0.143 | 0.143 |
| **Model composition** | | | | |
| Nonhydrogen atoms | 10875 | 21278 | | |
| Protein residues | 491 | 916 | | |
| Carbohydrates | 30 | 48 | | |
| **Validation** | | | | |
| Molprobity score | 1.63 | 1.78 | | |
| Clashscore | 3.73 | 6.05 | | |
| Poor rotamers (%) | 0 | 0 | | |
| **Ramachandran plot** | | | | |
| Favored regions (%) | 92.45 | 93.09 | | |
| Allowed (%) | 7.41 | 6.65 | | |
| Disallowed regions (%) | 0.15 | 0.26 | | |

**Supplemental Table 3. Data Collection and Refinement Statistics (Molecular Replacement)**

| | **GakV-2-F Head Domain** |
|---|---|
| **PDB ID** | 9EHU |
| **Data collection** | |
| Space group | P 31 2 1 |
| Cell dimensions | |
| *A*, *b*, *c* (Å) | 85.48, 85.48, 162.62 |
| α, β, γ (°) | 90, 90, 120 |
| Resolution (Å) | 67.38 – 1.42 |
| $R_{merge}$ | 0.167 (7.383) |
| $I / \sigma I$ | 10.4 (0.5) |
| Completeness (%) | 97.5 (91.6) |
| Redundancy | 16.8 (18.7) |
| **Refinement** | |
| Resolution (Å) | 67.38 – 1.42 |
| No. Unique reflections | 128347 (3998) |
| $R_{work}$ / $r_{free}$ | 0.2102/0.2271 |
| No. Atoms | 3756 |
| Protein | 3462 |
| Water | 224 |
| Ligand | 70 |
| *B*-factors | |
| Protein | 27.36 |
| Water | 30.74 |
| Ligand | 36.31 |
| R.M.S. Deviations | |
| Bond lengths (Å) | 0.0006 |
| Bond angles (°) | 0.86 |
| Ramachandran | |
| Favored (%) | 97.59 |
| Allowed (%) | 2.19 |
| Outlier (%) | 0.22 |

**Supplemental Data 1: Sequence Alignment of HNV-F Proteins**

```
NiV-M1   1   ------------MVVI-----LDKRCYCNL-LILILMISECSVGILHYEKLSKIGLVKGVTRKYKIKSNPLTKDIVIKMIPNV------SNMSQCTGSVMENYKTRLNGILTPIKGALEIYKNNTHDLVG
NiV-M2   1   ------------MVVI-----LDKRCYCNL-LILILMISECSVGILHYEKLSKIGLVKGVTRKYKIKSNPLTKDIVIKMIPNV------SNMSQCTGSVMENYKTRLNGILTPIKGALEIYKNNTHDLVG
NiV-M3   1   ------------MVVI-----LDKRCYCNL-LILILMISECSVGILHYEKLSKIGLVKGVTRKYKIKSNPLTKDIVIKMIPNV------SNMSQCTGSVMENYKTRLNGILTPIKGALEIYKNNTHDLVG
NiV-M4   1   ------------MVVI-----LDKRCYSNL-LILILMISECSVGILHYEKLSKIGLVKGVTRKYKIKSNPLTKDIVIKMIANV------SNMSQCTGSVMENYKTRLNGILTPIKGALEIYKNNTHDLVG
NiV-C    1   ------------MVVV-----LDKRCYSNL-LMLILMISECSVGILHYEKLSKIGLVKGITRKYKIKSNPLTKDIVIKMIPNV------SNMSQCTGSVMENYKTRLNGILTPIKGALEIYKNNTHDLVG
NiV-B1.1 1   ------------MAVI-----LNKRYYSNL-LILILMISECSVGILHYEKLSKIGLVKGITRKYKIKSNPLTKDIVIKMIPNV------SNMSQCTGSVMENYKTRLNGILTPIKGALEIYKNNTHDLVG
NiV-B1.2 1   ------------MAVI-----LNKRYYSNL-LILILMISECSVGILHYEKLSKIGLVKGITRKYKIKSNPLTKDIVIKMIPNV------SNMSQCTGSVMENYKTRLNGILTPIKGALEIYKNNTHDLVG
NiV-B1.3 1   ------------MAVI-----LNKRYYSNL-LILILMISECSVGILHYEKLSKIGLVKGITRKYKIKSNPLTKDIVIKMIPNV------SNMSQCTGSVMENYKTRLNGILTPIKGALEIYKNNTHDLVG
NiV-B1.4 1   ------------MAVI-----LNKRYYSNL-LILILMISECSVGILHYEKLSKIGLVKGITRKYKIKSNPLTKDIVIKMIPNV------SNMSQCTGSVMENYKTRLNGILTPIKGALEIYKNNTHDLVG
NiV-B2.3 1   ------------MAVI-----LNKRYYSNL-LILILMISECSVGILHYEKLSKVGLVKGITRKYKIKSNPLTKDIVIKMIPNV------SNMSQCTGSVMENYKTRLNGILTPIKGALEIYKNNTHDLVG
NiV-B3.1 1   ------------MAVI-----LNMRYYSNL-LILILMISECSVGILHYEKLSKIGLVKGITRKYKIKSNPLTKDIVIKMIPNV------SNMSQCTGSVMENYKTRLNGILMPIKGALEIYKNNTHDLVG
HeV-α1.1 1   ------------MAT------QEVRLKCLLCGIIVLVLSLEGLGILHYEKLSKIGLVKGITRKYKIKSNPLTKDIVIKMIPNV------SNVSKCTGTVMENYKSRLTGILSPIKGAIELYNNNTHDLVG
HeV-α1.2 1   ------------MAT------QEVRLKCLLCGIIVLVLSLEGLGILHYEKLSKIGLVKGITRKYKIKSNPLTKDIVIKMIPNV------SNVSKCTGTVMENYKSRLTGILSPIKGAIELYNNNTHDLVG
HeV-β1   1   ------------MATQ-----RVKVSYLIC-GIVISALSLEGLGILHYEKLSKIGLVKGVTRKYKIKSNPLTKDIVIKMIPNV------SNVSKCTGTVMENYKNRLTGILSPIKGAIELYNNNTHDLIG
HeV-β2   1   ------------MATQ-----GVKVSYLIC-GIVISALSLEGLGILHYEKLSKIGLVKGVTRKYKIKSNPLTKDIVIKMIPNV------SNVSKCTGTVMENYKNRLTGILSPIRGAIELYNNNTHDLIG
CedV-1   1   ------------MS-------NKRTTVLIIISYTLFYLNNAAIVGFDFDKLNKIGVVQGRVLNYKIKGDPMTKDLVLKFIPNI------VNITECVREPLSRYNETVRRLLLPIHNMLGLYLNNTNAKMT
GhV(+A)  1   ------------MV-------INGNIITIILLLILILKTQMSEGAIHYETLSKIGLIKGITREYKVKGTPSSKDIVIKLIPNV------TGLNKCTNISMENYKEQLDKILIPINNIIELYANSTKSAPG
AngV     1   ------------MGFR-----KNLRLLGIF-WLIILS------DTVDFERLASIGGIKGHSKLYKIKGHPTTKDIVIKLVPNL------NNLTVCSEMSIDGYLNLINAVIIPISQSLELMRNNVKDGTP
SHNV1    1   ---------------------MNYTIVGIM-IITFVHESQC----INYEQLASIGVIKGHTYNYKIRGPPNTKLMVVKLIPNINIDNLGGGLSNCSSKQMESHKELVEKVLSPVAQALETMRNRVTDYSG
NinV     1   --MSENSHWYNIIYII-----MKSRCSRLIGLIIVLTALMSAVVGIDFEGLSSIGVVKGRTLNYKIRGNSAYKILVIKLTPTIRSTSGAVNFENCTKDQISAHKVLVRNVLTPVKDALTSMKSKVTDYTP
SHNV2    1   ------------MSLKFRQSTNNRTNIRLILIIVLIKHMLC----IHYENLSSVGIIKGNTYNYKIKGDPSTKMLVVKLIPNI------DGLGNCTDKQMKDYKTLVKSTLQPVKDSLAQMLNNVETYNG
GakV-1   1   -------------MEL-----IKLFNILYL-VGYSGLFFTDVNAGLDYEGLSSIGVIKGPSYNYKIRGTPSTKLLVIKLIPNV------ESIDNCTQKQMSDYRALVKNVLTPVSESLSTMLNYIEQQSN
GakV-2   1   -------------MEL-----IKLFYILYL-VGYSGLFFTDVNAGLDYEGLSSIGVIKGPSYNYKIRGTPSTKLLVIKLIPNV------ESIDNCTQKQMSDYRALVKNVLTPVSESLSTMLNYIEQQSN
SHNV3    1   -------------MEL-----IKNYCLIYL-IFYSSTLIRPLQAGLDYEALASIGVIKGPSYNYKIRGTPSTKLLVIKLIPNV------GSLDNCTQKQMSDYRALVKNVLTPVSESLKTMLNYIEQQSN
SHNV11   1   MKLNNTRRDNKINKQHTTMACIKNCCLVYL-ILYSSTLIIPLEAGLDYESLASIGVIKGPSYNYKIRGTPSTKLLVIKLIPNV------GSLDNCTQKQMADYKALVKNVLTPVSESLKTMLNYIEQQSN
SHNV4    1   -------------MGL-----TKSTILVYL-LVYYHAHIIAINAGLDYEGLASIGVVKGPSYNYKIRGTPTTKLLVIKLIPNV------GSLDNCTQKQMADYKSLVKNVLTPVSDALSTMLNYIEQQSN
MelV     1   ----------MAKFVF-----LKTMCIGLL-----IIISDRVDSSMDFHSLSKIGIIKGKTYNYKIRGEPNTKLMVIKLIPNI------DVVENCSTTQVNNYKKLVRNVLTPVRIISRYYVKNVIEQNN
DewV-1   1   ---------------------MARLQVVIL-YLYLLVASDVVKASLDFDNMSKIGIIKGNTYNYKIRGEPTTKLMVVKLIPNI------DVVENCSATQLANYKKLVTNVLTPVKLSLDNMLKNVQDQNN
DarV-K   1   ------------MSS------SNKIRITVI-IINILISNYLIHCSMDFVELSRVGIIKGSTYNYKIRGEPSTKLMVVKLIPNV------GDVANCSATQVSNYKKLVKNVLSPVSNALNTMLENVQTENN
DarV-Cβ  1   ------------MSS------SNKIRITVI-IINILISNYLINCSMDFVELSRVGIIKGSTYNYKIRGEPSTKLMVVKLVPNV------GDVANCSTTQVSNYKKLVKNVLSPVSNALNTMLENVQTENN
LayV-C1  1   -------------MAF-----LKSAIICYL-LFYPHI----VKSSLHYDSLSKVGIIKGLTYNYKIKGSPSTKLMVVKLIPNI------DGVRNCTQKQFDEYKNLVKNVLEPVKLALNAMLDNVKSGNN
LayV-C2  1   -------------MAF-----LKSAIICYL-LFYPHI----VKSSLHYDSLSKVGIIKGLTYNYKIKGSPSTKLMVVKLIPNI------DRVRNCTQKQFDEYKNLVKNVLEPVKLALNAMLDNVKSGNN
MojV     1   ------------MA-------LNKNMFSSLFLGYLLVYATTVQSSIHYDSLSKVGVIKGLTYNYKIKGSPSTKLMVVKLIPNI------DSVKNCTQKQYDEYKNLVRKALEPVKMAIDTMLNNVKSGNN
SHNV5-1  1   ------------MINMD----SKRLKVKVVFPIYLIIYIQMSRASLDYDQLSKIGIVKGPSYNYKIKGSPSTKLMVVKLVPNI------KEVANCTHKQVENYKTLVRNVLDPVKMSLQAMLEKVKSGNN
SHNV5-2  1   ------------MINME----SKRLKVKVVFSIYFIIYIQMSRASLDYDQLSKIGIVKGPSYNYKIKGSPSTKLMVVKLVPNI------KEVANCTHKQIENYKTLVRNVLDPVKMSLQAMLEKVKSGNN

NiV-M1   107 DVRLAGVIMAGVAIGIATAAQITAGVALYEAMKNADNINKLKSSIESTNEAVVKLQETAEKTVYVLTALQDYINTNLVPTIDKISCKQTELSLDLALSKYLSDLLFVFGPNLQDPVSNSMTIQAISQAFG
NiV-M2   107 DVRLAGVIMAGVAIGIATAAQITAGVALYEAMKNADNINKLKSSIESTNEAVVKLQETAEKTVYVLTALQDYINTNLVPTIDKISCKQTELSLDLALSKYLSDLLFVFGPNLQDPVSNSMTIQAISQAFG
NiV-M3   107 DVRLAGVIMAGVAIGIATAAQITAGVALYEAMKNADNINKLKSSIESTNEAVVKLQETAEKTVYVLTALQDYINTNLVPTIDKISCKQTELSLDLALSKYLSDLLFVFGPNLQDPVSNSMTIQAISQAFG
NiV-M4   107 DVRLAGVIMAGVAIGIATAAQITAGVALYEAMKNADNINKLKSSIESTNEAVVKLQETAEKTVYVLTALQDYINTNLVPTIDKISCKQTELSLDLALSKYLSDLLFVFGPNLQDPVSNSMTIQAISQAFG
NiV-C    107 DVRLAGVIMAGVAIGIATAAQITAGVALYEAMKNADNINKLKSSIESTNEAVIKLQETAEKTVYVLTALQDYINTNLVPTIDKISCKQTELSLDLALSKYLSDLLFVFGPNLQDPVSNSMTIQAISQAFG
NiV-B1.1 107 DVRLAGVIMAGVAIGIATAAQITAGVALYEAMKNADNINKLKSSIESTNEAVVKLQETAEKTVYVLTALQDYINTNLVPTIDKISCKQTELSLDLALSKYSSDLLFVFGPNLQDPVSNSMTIQAISQAFG
NiV-B1.2 107 DVRLAGVIMAGVAIGIATAAQITAGVALYEAMKNADNINKLKSSIESTNEAVVKLQETAEKTVYVLTALQDYINTNLVPTIDKISCKQTELSLDLALSKYLSDLLFVFGPNLQDPVSNSMTIQAISQAFG
NiV-B1.3 107 DVRLAGVIMAGVAIGIATAAQITAGVALYEAMKNADNINKLKSSIESTNEAVVKLQETAEKTVYVLTALQDYINTNLVPTIDKISCKQTELSLDLALSKYLSDLLFVFGPNLQDPVSNSMTIQAISQAFG
NiV-B1.4 107 DVRLAGVIMAGVAIGIATAAQITAGVALYEAMKNADNINKLKSSIESTNEAVVKLQETAEKTVYVLTALQDYINTNLVPTIDKISCKQTELSLDLALSKYLSDLLFVFGPNLQDPVSNSMTIQAISQAFG
NiV-B2.3 107 DVRLAGVIMAGVAIGIATAAQITAGVALYEAMKNADNINKLKSSIESTNEAVVKLQETAEKTVYVLTALQDYINTNLVPTIDKISCKQTELSLDLALSKYLSDLLFVFGPNLQDPVSNSMTIQAISQAFG
NiV-B3.1 107 DVRLAGVIMAGVAIGIATAAQITAGVALYEAMKNADNINKLKSSIESTNEAVVKLQETAEKTVYVLTALQDYINTNLVPTIDKISCKQMELSLDLALSKYLSDLLFVFGPNLQDPVSNSMTIQAISQAFG
HeV-α1.1 107 DVKLAGVVMAGIAIGIATAAQITAGVALYEAMKNADNINKLKSSIESTNEAVVKLQETAEKTVYVLTALQDYINTNLVPTIDQISCKQTELALDLALSKYLSDLLFVFGPNLQDPVSNSMTIQAISQAFG
HeV-α1.2 107 DVKLAGVVMAGIAIGIATAAQITAGVALYEAMKNADNINKLKSSIESTNEAVVKLQETAEKTVYVLTALQDYINTNLVPTIDQISCKQTELALDLALSKYLSDLLFVFGPNLQDPVSNSMTIQAISQAFG
HeV-β1   107 DVKLAGVVMAGIAIGIATAAQITAGVALYEAMKNADNINKLKSSIESTNEAVVKLQETAEKTVYVLTALQDYINTNLVPTIDQISCKQTELALDLALSKYLSDLLFVFGPNLQDPVSNSMTIQAISQAFG
HeV-β2   107 DVKLAGVVMAGIAIGIATAAQITAGVALYEAMKNADNINKLKSSIESTNEAVVKLQETAEKTVYVLTALQDYINTNLVPTIDQISCKQTELALDLALSKYLSDLLFVFGPNLQDPVSNSMTIQAISQAFG
CedV-1   106 GLMIAGVIMGGIAIGIATAAQITAGFALYEAKKNTENIQKLTDSIMKTQDSIDKLTDSVGTSILILNKLQTYINNQLVPNLELLSCRQNKIEFDLMLTKYLVDLMTVIGPNINNPVNKDMTIQSLSLLFD
GhV(+A)  106 NARFAGVIIAGVALGVAAAAQITAGIALHEARQNAERINLLKDSISATNNAVAELQEATGGIVNVITGMQDYINTNLVPQIDKLQCSQIKTALDISLSQYYSEILTVFGPNLQNPVTTSMSIQAISQSFG
AngV     101 NENIFGAIVAGAALGIATAAQVTSVVALHKSNQNAAKINSLRDSITKTNQAVEQLSLGVQETVSVLMGLQDQINTNLVPKINILSCKQLSNTLNIMLLQYYSQILTVFGPNLRDPATVPVSIQALSQLFE
SHNV1    105 NYRFVGAVMAGAALGVATAATVTAGIALHQSNQNAKAIDQMKEAIRTTNKAVQELTLSTRQTLLVIDSLQNQINTQIVPAMNRLSCEVLGLTVGIQLTQYYSEILTYFGPALQDPIDSTLTIQAISHAFG
NinV     124 NSRLFGAIVAGAALVTATAATITAGVALHQSNQNAQAIANMKDAISKTNQAVSELKQGTQKLATVVDSLQNQINSQIIPVLNQLGCQTTGLTIGLYLTRYYSEILTVFGPAIQDPVNSALTIQAISKAFE
SHNV2    109 YVKFFGAVMAGAALGVATAATVTAGIALHQSNQNAKDIANMKDAIMKTNQAITTLSSASQKMLTVIDSIRGEINQQIIPVLNQLSCETVGLNLGIKLTQYYSQISTFFGPALQNPVQSILTIQAISHAFG
GakV-1   106 GVRLIGAVLAGAALGVATGAAITAGIALHKSNQNAQAIAQLKDAIKNTNQAVQTLKMANQELLGVVDSLRGQINTQIIPVLNKLSCDTVGLTLGIKLTQYYSEILTAFGPAIQDPVNSKLTIQAISGAFS
GakV-2   106 GVRLIGAVLAGAALGVATGAAITAGIALHKSNQNAQAIAQLKDAIKNTNQAVQTLKMANQELLGVVDSLRGQINTQIIPVLNKLSCDTVGLTLGIKLTQYYSEILTAFGPAIQDPVNSKLTIQAISGAFS
SHNV3    106 GVRLIGAVLAGAALGVATGAAITAGIALHKSNQNAQAIAQLKDSIKNTNMAVQTLKLANQEILGVVDSLRGQINTQIIPVLNQLSCDTVGLTLGIKLTQYYSEILTAFGPAIQDPVNSKLTIQAISGAFS
SHNV11   124 GVRLIGAVLAGAALGVATGAAITAGIALHKSNQNAQAIAQLKDSIKNTNMAVQTLKLANQEILGVVDSLRGQINTQIIPVLNQLSCDTVGLTLGIKLTQYYSEILTAFGPAIQDPVNSKLTIQAISGAFS
SHNV4    106 GVRLIGAVLAGAALGVATGAAITAGIALHKSNQNAQAIAQLKDAIKNTNQAVQTLKLANQEILGVVDSLRGQINTQIIPVINQLSCDTIGLTLGIKLTQYYSEILTAFGPAIQDPVNSKLTIQAISGAFS
MelV     105 RVRLFGAIMAGAALGVATAATVTAGIALHRSNENARNIALMKEAIKNTNQAVTKLQLAGQQTLAVIDNIRGEINNQIIPVINKLTCENIGLNVGIKLTQYYSEVLTAFGPAIQDPVNARITIQAISKVFN
DewV-1   103 RVRLFGAIMAGAALGVATAATVTAGIALHRSNENAKNIAKLKNAIQNTNLAITKLQMAGQQTLAVIDNIRGEINNQIIPVLNQLSCETVGLNVGIKLTQYYSEVLTAFGPAIQDPVNSKITIQAISKAFN
DarV-K   106 RYRLFGAIMAGAALGVATAATVTAGIALHRSNENARNIAAMKNAIQNTNQAITKLQMAGQQTLAVIDNIRGEINNQIIPLLNKLSCETVGLNVGIKLTQYYSEILTVFGPAIQDPINSKITIQAISKAFG
DarV-Cβ  106 RYRLFGAIMAGAALGVATAATVTAGIALHRSNENARNIAAMKNAIQNTNQAITKLQMAGQQTLAVIDNIRGEINNQIIPLLNKLSCETVGLNVGIKLTQYYSEILTVFGPAIQDPINSKITIQAISKAFG
LayV-C1  102 KYRFAGAIMAGVALGVATAATVTAGIALHRSNENAQAIANMKNAIQNTNEAVKQLQLANKQTLAVIDTIRGEINNNIIPVINQLSCDTIGLSVGIKLTQYYSEILTAFGPALQNPVNTRITIQAISSVFN
LayV-C2  102 KYRFAGAIMAGVALGVATAATVTAGIALHRSNENAQAIANMKNAIQNTNEAVKQLQLANKQTLAVIDTIRGEINNNIIPVINQLSCDTIGLSVGIKLTQYYSEILTAFGPALQNPVNTRITIQAISSVFN
MojV     106 KYRFAGAIMAGVALGVATAATVTAGIALHRSNENAQAIANMKSAIQNTNEAVKQLQLANKQTLAVIDTIRGEINNNIIPVINQLSCDTIGLSVGIRLTQYYSEIITAFGPALQNPVNTRITIQAISSVFN
SHNV5-1  109 KYRFAGAIMAGVALGVATAATVTAGIALHRSSENAQAIARIKGAIQNTNEAVKQLQLANKQMLAVIDTIRGEINENIIPIMNELSCETIGLNVGIKLTQYYSEIITAFGPALQNPIGSKITIQAISSAFN
SHNV5-2  109 KYRFAGAIMAGVALGVATAATVTAGIALHRSSENAQAIARIKGAIQNTNEAVKQLQLGNKQMLAVIDTIRGEINENIIPIMNELSCETIGLNVGIKLTQYYSEIITAFGPALQNPIGSKITIQAISSAFN

NiV-M1   237 GNYETLLRTLGYATEDFDDLLESDSITGQIIYVDLSSYYIIVRVYFPILTEIQQAYIQELLPVSFNNDNSEWISIVPNFILVRNTLISNIEIGFCLITKRSVICNQDYATPMTNNMRECLTGSTEKCPRE
NiV-M2   237 GNYETLLRTLGYATEDFDDLLESDSITGQIIYVDLSSYYIIVRVYFPILTEIQQAYIQELLPVSFNNDNSEWISIVPNFILVRNTLISNIEIGFCLITKRSVICNQDYATPTTNNMRECLTGSTEKCPRE
NiV-M3   237 GNYETLLRTLGYAIEDFDDLLESDSITGQIIYVDLSSYYIIVRVYFPILTEIQQAYIQELLPVSFNNDNSEWISIVPNFILVRNTLISNIEIGFCLITKRSVICNQDYATPMTNNMRECLTGSTEKCPRE
NiV-M4   237 GNYETLLRTLGYATEDFDDLLESDSITGQIIYVDLSSYYIIVRVYFPILTEIQQAYIQELLPVSFNNDNSEWISIVPNFILVRNTLISNIEIGFCLITKRSVICNQDYATPMTNNMRECLTGSTEKCPRE
NiV-C    237 GNYETLLRTLGYATEDFDDLLESDSITGQIIYVDLSSYYIIVRVYFPILTEIQQAYIQELLPVSFNNDNSEWISIVPNFILVRNTLISNIEIGFCLITKRSVICNQDYATPMTNNMRECLTGSTEKCPRE
NiV-B1.1 237 GNYETLLRTLGYATEGFDDLLESDSITGQIIYVDLSGYYIIVRVYFPILTEIQQAYIQELLPVSFNNDNSEWISIVPNFILVRNTLISNIEIGFCLITKRSVICNQDYATPMTNNMRECLTGSTEKCPRE
NiV-B1.2 237 GNYETLLRTLGYATEDFDDLLESDSITGQIIYVDLSSYYIIVRVYFPILTEIQQAYIQELLPVSFNNDNSEWISIVPNFILVRNTLISNIEIGFCLITKRSVICNQDYATPMTNNMRECLTGSTEKCPRE
NiV-B1.3 237 GNYETLLRTLGYATEDFDDLLESDSITGQIIYVDLSGYYIIVRVYFPILTEIQQAYIQELLPVSFNNDNSEWISIVPNFILVRNTLISNIEIGFCLITKRSVICNQDYATPMTNNMRECLTGSTEKCPRE
NiV-B1.4 237 GNYETLLRTLGYATEDFDDLLESDSITGQIIYVDLSGYYIIVRVYFPILTEIQQAYIQELLPVSFNNDNSEWISIVPNFILVRNTLISNIEIGFCLITKRSVICNQDYATPMTNNMRECLTGSTEKCPRE
NiV-B2.3 237 GNYETLLRTLGYTTEDFDDLLESDSITGQIIYVDLSGYYIIVRVYFPILTEIQQAYIQELLPVSFNNDNSEWISIVPNFILVRNTLISNIEIGFCLITKRSVICNQDYATPMTNNMRECLTGSTEKCPRE
NiV-B3.1 237 GNYETLLRTLGYATEDFDDLLESDSITGQIIYVDLSGYYIIVRVYFPILTEIQQAYIQELLPVSFNNDNSEWISIVPNFILVRNTLISNIEIGFCLITKRSVICNQDYATPMTNNMRECLTGSTEKCPRE
HeV-α1.1 237 GNYETLLRTLGYATEDFDDLLESDSIAGQIVYVDLSSYYIIVRVYFPILTEIQQAYVQELLPVSFNNDNSEWISIVPNFVLIRNTLISNIEVKYCLITKKSVICNQDYATPMTASVRECLTGSTDKCPRE
HeV-α1.2 237 GNYETLLRTLGYATEDFDDLLESDSITGQIVYVDLSSYYIIVRVYFPILTEIQQAYVQELLPVSFNNDNSEWISIVPNFVLIRNTLISNIEVKYCLITKKSVICNQDYATPMTASVRECLTGSTDKCPRE
HeV-β1   237 GNYETLLRTLGYATEDFDDLLESDSITGQIVYVDLSSYYIIVRVYFPILTEIQQAYVQELLPVSFNNDNSEWISIVPNFVLIRNTLISNIEVKYCLTTKKSVICNQDYATPMTSTVRECLTGSTDKCPRE
HeV-β2   237 GNYETLLRTLGYATEDFDDLLESDSITGQIVYVDLSSYYIIVRVYFPILTEIQQAYVQELLPVSFNNDNSEWISIVPNFVLIRNTLISNIEVKYCLTTKKSVICNQDYATPMTSTVRECLTGSTDKCPRE
CedV-1   236 GNYDIMMSELGYTPQDFLDLIESKSITGQIIYVDMENLYVVIRTYLPTLIEVPDAQIYEFNKITMSSNGGEYLSTIPNFILIRGNYMSNIDVATCYMTKASVICNQDYSLPMSQNLRSCYQGETEYCPVE
GhV(+A)  236 GNIDLLLNLLGYTANDLLDLLESKSITGQITYINLEHYFMVIRVYYPIMTTISNAYVQELIKISFNVDGSEWVSLVPSYILIRNSYLSNIDISECLITKNSVICRHDFAMPMSYTLKECLTGDTEKCPRE
AngV     231 GNLELLTSSLGISSTDFNDLLTGKLITGSIIWADTQGHYLILRVNIPDLIEVPGAVIQEFIKIGYNFGGSLWMPTIPNKIIIRGYHLSLIDSTNCIITDNSYVCDRDYSLPMNPILRECFEGNTSSCGRE
SHNV1    235 GNFDILMKTMGYTVGDLYDVLKGDLITGKIISVNPKEGFIALEVRFPTLTQVNNAIVQELMPISFNDKGDEWISTVPRYVLERVLYLSNIDISLCSVGETSVVCDNDYASPMSHQLRECLQTNTSYCPRE
NinV     254 GNFDKLMSAMGYSVSDLKDVLESDLVRGKIIDVDPDVGYMALQIEFPQITPVTGAHIQELLPISFNYKGDEWITILPSFVLNRMTYLSNIDIAHCLVTETSVICDNDLAMPMSTQMRDCLLGNTSKCSRE
SHNV2    239 GNFNELMTVMGYTGSDFQDIIQGNLITGSVIGVDPEVGYIALEIRIPSLTVVPNAYVQELLPVSFNIDGDEWMTIVPNYVLTRTTYLSNIDINRCLITDKSVICGNDYATPMSNQLINCLNGNTQHCARE
GakV-1   236 GNFDEMMKVMGYSGSDLHDILQGDLITGNIIGVDPDVGYIALEINFPTLTEIPNAVIQELMPISFNDKGDEWMALVPRYVLLRTTYISNIDISKCLLTERSVICYNDYATPMSFDIIRCLTGNLTYCPRE
GakV-2   236 GNFDEMMKVMGYSGSDLHDILQGDLITGNIIGVDPDVGYIALEINFPTLTEIPNAVIQELMPISFNDKGDEWMALVPRYVLLRTTYISNIDISKCLLTERSVICYNDYATPMSFDIIRCLTGNLTYCPRE
SHNV3    236 GNFDEMMKVMGYTGSDLHDILQGDLITGNIIGVDPDVGYIALEIHFPTLTEIPNAVIQELMPISFNDKGDEWMALVPRYVLLRTTYVSNIDISKCLLTERSVICYNDYATPMSFDVLRCLTGNLTFCPRE
SHNV11   254 GNFDEMMKVMGYTGSDLHDILQGDLITGNIIGVDPDVGYIALEIHFPTLTEIPNAVIQELMPISFNDKGDEWMALVPRYVLLRTTYVSNIDISKCLLTERSVICYNDYATPMSFDVLRCLTGNLTFCPRE
SHNV4    236 GNFDEMMKVMGYTGSDLHDILQGDLITGNIIGVDPDIGYIALEIHFPTLTEIPNAVIQELMPISFNDKGDEWMTLVPRYVLLRTTYVSNIDISKCLITERSVICYNDYATPMSFDVIRCLTGNLTYCPRE
MelV     235 NNFDELLNVMGYSTQDLYEVLHGGLIRGNVISADPEVGYLALEIEFPNLSVVPNAYIQEIMPISFNVDGDEWVTVVPRHTLIRTTLLSNIDITTCSIIESSIICNNDYALPMSNELINCLQGSTDLCARE
DewV-1   233 NNFDELLKVMGYTSQDLYEILHGDLITGNVISADPEAGYIALEIEFPNLSTVPNAYVQELMPISFNVDGDEWVTLVPRYVLIRTTFLSNIDISLCSIMETSIVCNNDYALPMSSELINCLQGDTGVCARE
DarV-K   236 GNFDELLKVMGYTSQDLYEILHGGLITGNIIGVDPDTGYIALEIEFPNLSIVQNAYIQELMPISFNVDGDEWVTLAPRFVLIRTTLLSNIDTSMCTIIDSSVICNNDYALPMSTELINCLQGMTESCARE
DarV-Cβ  236 GNFDELLNVMGYTSQDLYEILHGGLITGNIIGVDPDTGYIALEIEFPNLSIVQNAYIQELMPISFNVDGDEWVTLAPRFVLTRTTLLSNIDTSMCTIIDSSVICNNDYALPMSTELINCLQGMTESCARE
LayV-C1  232 RNFDELLKIMGYTSGDLYEILHSGLIRGNIIDVDVEAGYIALEIEFPNLTLVPNAVVQELMPISYNVDGDEWVTLVPRFVLTRTTLLSNIDTSRCTVTESSVICDNDYALPMSYELIGCLQGDTSKCARE
LayV-C2  232 RNFDELLKIMGYTSGDLYEILHSGLIRGNIIDVDVEAGYIALEIEFPNLTLVPNAVVQELMPISYNVDGDEWVTLVPRFVLTRTTLLSNIDTSRCTVTESSVICDNDYALPMSYELIGCLQGDTSKCARE
MojV     236 GNFDELLKIMGYTSGDLYEILHSELIRGNIIDVDVDAGYIALEIEFPNLTLVPNAVVQELMPISYNIDGDEWVTLVPRFVLTRTTLLSNIDTSRCTITDSSVICDNDYALPMSHELIGCLQGDTSKCARE
SHNV5-1  239 GNFDELLKSMGYTSNDLYEVLQSGLIRGNIIDVDPEVGYIALEIEFPNLTLVPNAIIQELMPISFNSEGDEWVTLMPRFILTRTTLLSNIDITKCTVTDRSVICDNDYALPMSNQLIECLRGDTMKCTRE
SHNV5-2  239 GNFDELLKSMGYTSNDLYEVLQSGLIRGNIIDVDPEAGYIALEIEFPNLTLVPNAIIQELMPISFNSEGDEWVTLMPRFILTRTTLLSNIDITKCTVTDRSVICDNDYALPMSNQLIECLRGDTMKCTRE

NiV-M1   367 LVVSSHVPRFALSNGVLFANCISVTCQCQTTGRAISQSGEQTLLMIDNTTCPTAVLGNVIISLGKYLGSVNYNSEGIAIGPPVFTDKVDISSQISSMNQSLQQSKDYIKEAQRLLDTVNPSLISMLSMII
NiV-M2   367 LVVSSHVPRFALSNGVLFANCISVTCQCQTTGRAISQSGEQTLLMIDNTTCPTAVLGNVIISLGKYLGSVNYNSEGIAIGPPVFTDKVDISSQISSMNQSLQQSKDYIKEAQRLLDTVNPSLISMLSMII
NiV-M3   367 LVVSSHVPRFALSNGVLFANCISVTCQCQTTGRAISQSGEQTLLMIDNTTCPTAVLGNVIISLGKYLGSVNYNSEGIAIGPPVFTDKVDISSQISSMNQSLQQSKDYIKEAQRLLDTVNPSLISMLSMII
```

```
NiV-M4   367 LVVSSHVPRFALSNGVLFANCISVTCQCQTTGRAISQSGEQTLLMIDNTTCPTAVLGNVIISLGKYLGSVNYNSEGIAIGPPVFTDKVDISSQKSSMNQSLQQSKDYIKEAQRLLDTVNPSLISMLSMII
NiV-C    367 LVVSSHVPRFALSNGVLFANCISVTCQCQTTGRAISQSGEQTLLMIDNTTCPTAVLGNVIISLGKYLGSVNYNSEGIAVGPPVFTDKVDISSQISSMNQSLQQSKDYIKEAQRLLDTVNPSLISMLSMII
NiV-B1.1 367 LVVSSHVPRFALSNGVLFANCISVTCQCQTTGRAISQSGEQTLLMIDNTTCPTAVLGNVIISLGKYLGSVNYNSEGIAIGPPVFTDKVDISSQISSMNQSLQQSKDYIKEAQRLLDTVNPSLISMLSMII
NiV-B1.2 367 LVVSSHVPRFALSNGVLFANCISVTCQCQTTGRAISQSGEQTLLMIDNTTCPTAVLGNVIISLGKYLGSVNYNSEGIAIGPPVFTDKVDISSQISSMNQSLQQSKDYIKEAQRLLDTVNPSLISMLSMII
NiV-B1.3 367 LVVSSHVPRFALSNGVLFANCISVTCQCQTTGRAISQSGEQTLLMIDNTTCPTAVLGNVIISLGKYLGSVNYNSEGIAIGPPVFTDKVDISSQISSMNQSLQQSKDYIKEAQRLLDTVNPSLISMLSMII
NiV-B1.4 367 LVVSSHVPRFALSNGVLFANCISVTCQCQTTGRAISQSGEQTLLMIDNTTCPTAVLGNVIISLGKYLGSINYNSEGIAIGPPVFTDKVDISSQISSMNQSLQQSKDYIKEAQRLLDTVNPSLISMLSMII
NiV-B2.3 367 LVVSSHVPRFALSNGVLFANCISVTCQCQTTGRAISQSGEQTLLMIDNTTCPTAVLGNVIISLGKYLGSVNYNSEGIAIGPPVFTDKVDISSQISSMNQSLQQSKDYIKEAQRLLDTVNPSLISMLSMII
NiV-B3.1 367 LVVSSHVPRFALSNGVLFANCISVTCQCQTTGRAISQSGEQTLLMIDNTTCPTAVLGNVIISLGKYLGSVNYNSEGIAIGPPVFTDKVDISSQISSMNQSLQQSKDYIKEAQRLLDTVNPSLISMLSMII
HeV-α1.1 367 LVVSSHVPRFALSGGVLFANCISVTCQCQTTGRAISQSGEQTLLMIDNTTCTTVVLGNIIISLGKYLGSINYNSESIAVGPPVYTDKVDISSQISSMNQSLQQSKDYIKEAQKILDTVNPSLISMLSMII
HeV-α1.2 367 LVVSSHVPRFALSGGVLFANCISVTCQCQTTGRAISQSGEQTLLMIDNTTCTTVVLGNIIISLGKYLGSINYNSESIAVGPPVYTDKVDISSQISSMNQSLQQSKDYIKEAQKILDTVNPSLISMLSMII
HeV-β1   367 LVVSSHVPRFALSGGVLFANCISVTCQCQTTGRAISQSGEQTLLMIDNTTCTTVVLGNIIISLGKYLGSTNYNSESIAVGPPVYTDRVDISSQISSMNQSLQQSKDYIKEAQRILDTVNPSLISMLSMIV
HeV-β2   367 LVVSSHVPRFALSGGVLFANCISVTCQCQTTGRAISQSGEQTLLMIDNTTCTTVVLGNIIISLGKYLGSTNYNSESIAVGPPVYTDRVDISSQISSMNQSLQQSKDYIKEAQRILDTVNPSLISMLSMIV
CedV-1   366 AVIASHSPRFALTNGVIFANCINTICRCQDNGKTITQNINQFVSMIDNSTCNDVMVDKFTIKVGKYMGRKDINNINIQIGPQIIIDKVDLSNEINKMNQSLKDSIFYLREAKRILDSVNISLISPSVQLF
GhV(+A)  366 AVVTSYVPRFAISGGVIYANCLSTTCQCYQTGKVIAQDGSQTLMMIDNQTCSIVRIEEILISTGKYLGSQEYNTMHVSVGNPVFTDKLDITSQISNINQSIEQSKFYLDKSKAILDKINLNLIGSVPISI
AngV     361 MVLNSYIPLYALSEGVIYANCLATSCKCATTNKPIVQSTSTVITMIDNSKCPVVEVGKQMISVGSYLGQVSPNNLTIEIGPPVYTEPVDITNQLGNINETLSKTLDLIKQSNSILDMITGGLNPQIGSMI
SHNV1    365 RVLASYVPKFALSQGVIFANCIATTCRCADDGRAISQSSSQTVLLLTSKDCKVYEVQSMMISTGEYLGESIFENTDIPLGPSIVIDKIDISGQLAEINKTLDHVDNTIKDSNDILDKIDVSTVSAISMVI
NinV     384 QVITSYVPRFALSGGVIYANCLNTQCACATNGRSISQSSRQTIMMLDDKDCKIYEVGGIFISVSTYMGIGKFENENISIGPPVVVDKIDVSGQLAEVNQSLSKTEELLEESNEYLSQVHVSLVSLKSMII
SHNV2    369 AVVTSYVPKFALSGGVIYANCLSTVCRCIDKDQPISQSLSQTLMMLDNQHCNVYQISNVLISTGRYLGDAEFRNEGIDLGPPIVVDKIDLGGQIADINQTISDAEEFIEESNKILSKINPKIISVKSMVV
GakV-1   366 QIMASHVPKFALSGGVIYANCLSAVCRCAVDGVPIVQSLKATVMMLDNKSCRVYQIGELLISTGAYLGSIEFKNENIELGPPIVIDKIDLGGQIAGINQTLQGVEDYIDKSNEILDQVNPSVTSLGAMII
GakV-2   366 QIMASHVPKFALSGGVIYANCLSAVCRCAVDGVPIVQSLKATVMMLDNKNCRVYQIGELLISTGAYLGSIEFKNENIELGPPIVIDKIDLGGQIAGINQTLQGVEDYIDKSNDILDQVNPSVTSLGAMII
SHNV3    366 QIIASHVPKFALSGGVIYANCLSAVCRCAVDGVPIVQSLKTTIMMLDNKNCRVYQIGELLISTGAYLGSVEFKNENIELGPPIVVDKIDLGGQIAGINQTLQGVEDYIDKSNEILDQVNPSVTSLGAMII
SHNV11   384 QIIASHVPKFALSGGVIYANCLSAVCRCAVDGVPIVQSLKTTIMMLDNKNCRVYQIGELLISTGSYLGSVEFKNENIELGPPIVVDKIDLGGQIAGINQTLQGVEDYIDKSNEILDQVNPSVTSLGAMII
SHNV4    366 QIIASYVPRFALSGGVIYANCLSTVCRCAVDGVPIVQSLKATIMMLDNKNCRVYQIGELLISTGAYLGSVEFKNENIDLGPPIVIDKVDLGGQIAGINQTLQGVEDYIDKSNEILDQVNPSVTSLGAMII
MelV     365 KVISNYVPKFALSDGVVYANCLSTVCRCMDNGVPISQSLKSTVMMLDDKKCTIYQIGDILISVGKYMGHIDYNPENVVLGPPIVIDKIDIGNQLAGINQTLQEAGDFIEKSEEILNSINPSVLTLSSMIT
DewV-1   363 KVISSYVPKFALSGGVVYANCLSTVCRCMDNGTPISQGIRHTVALLDNKKCSVYQVGDILISVGKYLGHLDYNTEDIVLGPPIVIDKIDIGNQLAGINSSLQQAENYIDKSNEILKSINPTIISTNSMIT
DarV-K   366 KVISSYVPKFALSGGVIYANCLSTVCRCMDNQKPISQSLRSTVVMLDNKMCKVYQIGDILISVGEYKGAVEYNPEDVHLGPPIVLDKIDIGNQLAGINQTLSEAGDFISKSEEILKDINPAILSVSSMIV
DarV-Cβ  366 KVISSYVPKFALSGGVIYANCLSTVCRCMDNQKPISQSLRSTVVMLDNKMCKVYQIGDILISVGEYKGAVEYNPEDVHLGPPIVLDKIDIGNQLAGINQTLSEAGDFISKSEEILKDINPAILSVSSMIV
LayV-C1  362 KVVSSYVPRFALSDGLVYANCLNTICRCMDTDTPISQSLGTTVSLLDNKKCLVYQVGDILISVGSYLGEGEYSADNVELGPPVVIDKIDIGNQLAGINQTLQNAEDYIEKSEEFLKGINPSIITLGSMAV
LayV-C2  362 KVVSSYVPRFALSDGLVYANCLNTICRCMDTDTPISQSLGTTVSLLDNKKCLVYQVGDILISVGSYLGEGEYSADNVELGPPVVIDKIDIGNQLAGINQTLQNAEDYIEKSEEFLKGINPSIITLGSMAV
MojV     366 KVVSSYVPKFALSDGLVYANCLNTICRCMDTDTPISQSLGATVSLLDNKRCSVYQVGDVLISVGSYLGDGEYNADNVELGPPIVIDKIDIGNQLAGINQTLQEAEDYIEKSEEFLKGVNPSIITLGSMVV
SHNV5-1  369 KVMSSYIPKFALSDGVVYANCLNTICRCMDTDTPITQSLRSTVTLLDNKACLVYQIGDILISVGSYLGNTEYNTQNITLGPPIVIDKIDIGNQLAGINQSLQNAEDYIEKSDEFLKGINPSVITLGSMVV
SHNV5-2  369 KVMSSYIPKFALSDGVVYANCLNTICRCMDTDTPITQSLRSTVTLLDNKACLVYQIGDILISVGSYLGNTEYNTQNITLGPPIVIDKIDIGNQLAGINQSLQNAEDYIEKSEEFLKGINPSVITLGSMVV

NiV-M1   497 LYVLSIASLCIGLITFISFIIVEKKRNTYSRL-------------EDRRVR----PTSSGDLYYIGT---------------------------------
NiV-M2   497 LYVLSIASLCIGLITFISFIIVEKKRNTYSRL-------------EDRRVR----PTSSGDLYYIGT---------------------------------
NiV-M3   497 LYVLSIASLCIGLITFISFIIVEKKRNTYSRL-------------EDRRVR----PTSSGDLYYIGT---------------------------------
NiV-M4   497 LYVLSIASLCIGLITFISFIIVEKKRNTYSRL-------------EDRRVR----PTSSGDLYYIGT---------------------------------
NiV-C    497 LYVLSIASLCIGLITFISFIIVEKKRNTYSRL-------------EDRRVR----PTSSGDLYYIGT---------------------------------
NiV-B1.1 497 LYVLSIASLCIGLITFISFIIVEKKRNTYSRL-------------EDRRVR----PTSSGDLYYIGT---------------------------------
NiV-B1.2 497 LYVLSIASLCIGLITFISFIIVEKKRNTYSRL-------------EDRRVR----PTSSGDLYYIGT---------------------------------
NiV-B1.3 497 LYVLSIASLCIGLITFISFIIVEKKRNTYSRL-------------EDRRVR----PTSSGDLYYIGT---------------------------------
NiV-B1.4 497 LYVLSIASLCIGLITFISFIIVEKKRNTYSRL-------------EDRRVR----PTSSGDLYYIGT---------------------------------
NiV-B2.3 497 LYVLSIASLCIGLITFISFIIVEKKRNTYSRL-------------EDRRVR----PTSSGDLYYIGT---------------------------------
NiV-B3.1 497 LYVLSIASLCIGLITFISFIIVEKKRNTYSRL-------------EDRRVR----PTSNGDLYYIGT---------------------------------
HeV-α1.1 497 LYVLSIAALCIGLITFISFVIVEKKRGNYSRL-------------DDRQVR----PVSNGDLYYIGT---------------------------------
HeV-α1.2 497 LYVLSIAALCIGLITFISFVIVEKKRGNYSRL-------------DDRQVR----PVSNGDLYYIGT---------------------------------
HeV-β1   497 LYVLSTAALCIGLITLISFIIVEKKRGNYSRL-------------DDRRVR----PVSNGDLYYIGT---------------------------------
HeV-β2   497 LYVLSTAALCIGLITLISFIIVEKKRGNYSRL-------------DDRRVR----PVSNGDLYYIGT---------------------------------
CedV-1   496 LIIISVLSFIILLIIIVYLYCKSKHSYKYNKFIDDPDYYN-----DYKRERINGKASKSNNIYYVGD---------------------------------
GhV(+A)  496 LFIIAILSLILSIITFVIVMIIVRRYNKYTPLIN-----------SDPSSR----RSTIQDVYIIPNPGEHSIRSAARSIDRDRD---------------
AngV     491 SLILAAIAILLSGITCLFSTKSYVKCNKLQMNC----------FKSYERMD----P-----IYHSQQN--------------------------------
SHNV1    495 LYVVIALIGFMSALSLLLSVRTFSRCTTLGSQF------------AYQRQD----P-TLGDVHYAFTSNIPKNQRSKNSNNLSNDLGSTNGSFSSNE---
NinV     514 LYVFIGLIGLLAAAALLVSVRSLSMLKTMGPAT------------LMNNIQ----P-TMHNLQYSGHSTINSKSSLINTHQSSHNHKYGSVNGSYTSDES
SHNV2    499 LYVVVALIAILAMVSLVLSIRLTMQVGTKINQF------------AYTKHV----P-SMENVQYIGTR--------------------------------
GakV-1   496 VYIFIALAILLGLIALIMDVKLNSQVKILINQSMINQMRAGLENPGYSRSL----P-SFGSIASRGSSQELVNTN-------------------------
GakV-2   496 VYIFIALAILLGLIALIMDVKLNSQVKILINQSMINQMRAGLENPGYSRSL----P-SFGSVASRGSSQELVNTN-------------------------
SHNV3    496 VYIFIALAILIGLIALMIDVKLNSQIKILINQSIINQMRAGLENPGYSKSL----P-SFSSTVSKGSSQELINTH-------------------------
SHNV11   514 VYIFIALAILIGLIALMIDVKLNSQIKILINQSIINQMRAGLENPGYSRSL----P-SFSSTVSKGSSQELINTH-------------------------
SHNV4    496 VYIFIALAILIGLIALMIDVKLNSQIKILINQSIVNQMRAGLENPGYSRSL----P-SFSSIASKGSSQELVNTN-------------------------
MelV     495 LYIFLIITVIIAVAALVLSIRLTIKARILTNQF------------AYGRHS----P-SMDNVSYVSK---------------------------------
DewV-1   493 LYIFIIITVIIAITSLVLSLRLTMKAKILTNQF------------AYGRHS----P-SMDNVSYVSR---------------------------------
DarV-K   496 LYIFIIVITIISLTSLILSIRLTIKAKIFNNQF------------AYGRHS----P-SMDNVSYVTH---------------------------------
DarV-Cβ  496 LYIFIIVITIISLTSLILSIRLTIKAKIFNNQF------------AYGRHS----P-SMDNVSYVTH---------------------------------
LayV-C1  492 LYIFMIVIAVISIIALVLSIKLTVKGNVVRQQF------------AYTQHV----P-SMENVNYVSH---------------------------------
LayV-C2  492 LYIFMIVIAVISIIALVLSIKLTVKGNVVRQQF------------AYTQHV----P-SMENVNYVSH---------------------------------
MojV     496 LYIFMILIAIVSVIALVLSIKLTVKGNVVRQQF------------TYTQHV----P-SMENINYVSH---------------------------------
SHNV5-1  499 LYIFMILIAIISIIALVLSIRLTTKSNMQKAQF------------SYTRQA----P-SMDNINYVSR---------------------------------
SHNV5-2  499 LYIFMILIAIISIIALVLSIRLTTKSNMQKAQF------------SYTRQA----P-SMDNINYVSR---------------------------------
```

**Supplemental Data 2: Sequence Alignment of HNV-G Proteins**

```
NiV-M1   1   ---------------------------MPTENKKVRFENTTSDKGKIPSKVIK-SYYGTMDI----KK----INEGLLDSKI--LSAFNTVIALLGSIV-IIVMNIMIIQNYTRSTDNQ--AVIKDALQG
NiV-M3   1   ---------------------------MPAENKKVRFENTTSDKGKNPSKVIK-SYYGTMDI----KK----INEGLLDSKI--LSAFNTVIALLGSIV-IIVMNIMIIQNYTRSTDNQ--AVIKDALQG
NiV-B1.5 1   ---------------------------MPTESKKVRFENTASDKGKNPSKVIK-SYYGTMDI----KK----INEGLLDSKI--LSAFNTVIALLGSIV-IIVMNIMIIQNYTRSTDNQ--AMIKDALQS
NiV-B1.6 1   ---------------------------MPTESKKVRFENTASDKGKNPSKVIK-SYYGTMDI----KK----INEGLLDSKI--LSAFNTVIALLGSIV-IIVMNIMIIQNYTRSTDNQ--AMIKDALQS
NiV-B2.4 1   ---------------------------MPTESKKVRFENTASDKGKNPSKVIK-SYYGTMDI----KK----INEGLLDSKI--LSAFNTVIALLGSIV-IIVMNIMIIQNYTRSTDNQ--AMIKDALQS
NiV-B3.1 1   ---------------------------MPTESKKVRFENTASDKGKNPSKVIK-SYYGTMDI----KK----INEGLLDSKI--LSAFNTVIALLGSIV-IIVMNVMIIQNYTRSTDNQ--AMIKDALQS
NIV-I1.3 1   ---------------------------MPTESKKVRFENTASDKGKNPSKVIK-SYYGTMDI----KK----INEGLLDSKI--LSAFNTVIALLGSIV-IIVMNIMIIQNYTRSTDNQ--AMIKDALQS
HeV-α1.2 1   ---------------------------MMADSKLVSLNNNLSGKIKDQGKVIK-NYYGTMDI----KK----INDGLLDSKI--LGAFNTVIALLGSII-IIVMNIMIIQNYTRTTDNQ--ALIKESLQS
HeV-α3   1   ---------------------------MMADSKLVSLNNNLSGKIKDQGKVIK-NYYGTMDI----KK----INDGLLDSKI--LGAFNTVIALLGSII-IIVMNIMIIQNYTRTTDNQ--ALIKESLQS
HeV-α4   1   ---------------------------MMADSKLVSLNNNLSGKIKDQGKVIK-NYYGTMDI----KK----INDGLLDSKI--LGAFNTVIALLGSII-IIVMNIMIIQNYTRTTDNQ--ALIKESLQS
HeV-α5   1   ---------------------------MMADSKLVSPNNNLSGKIKDQGKVIK-NYYGTMDI----KK----INDGLLDSKI--LGAFNTVIALLGSII-IIVMNIMIIQNYTRTTDNQ--ALIKESLQS
HeV-α6   1   ---------------------------MMADSKLVSLNNNLSGKIKDQGKVIK-NYYGTMDI----KK----INDGLLDSKI--LGAFNTVIALLGSII-IIVMNIMIIQNYTRTTDNQ--ALIKESLQS
HeV-β1   1   ----------------------------MAESKVVRPSSSLSSKIKDQGKIIK-NYYGTMDI----KK----INDGLLDSKI--LGAFNTVIALLGSII-IIVMNVMIIQNYARTTDNQ--ALIKDSLQS
HeV-β3   1   ----------------------------MAESKVVRPSSSLSSKIKDQGKIIK-NYYGTMDI----KK----INDGLLDSKI--LGAFNTVIALLGSII-IIVMNVMIIQNYTRTTDNQ--ALIKDSLQS
CedV-1   1   MLSQLQ----KNYLD----NSNQQGDKMNNPDKKLSVNFNPLELDKGQKDLNK-SYYVKNKN----YNVSNLLNESLHDIKFCIYCIFSLL-IIITIIN-IITI-SIVITRLKVHEENN--GMESPNLQS
GhV      1   -MPQKT----VEFIN------------MNSPLER-GVSTLSDKKTLNQSKITKQGYFGLGSHSERNWKKQKNQNDHYMTVSTMILE----ILVVLGIMFNLIVL-TMV--YYQNDNINQRMAELTSNITV
AngV     1   -MSQKK--------------------------SLIIKGHDYSPKDKNQSS----KYYNSYESD------QDTINPN--EFKCMLTLIFLVL--LL-TIASVIVLGICN--VVKLDHESKQISINTMSINK
LayV-C2  1   ---------------------------MATNKKD----VI-KTTESTRED-KVKKFYGVETA----EK----VADSISSNKVFIL--MNTLLILTGAII-TITLNVTNLTTIKNQQA--MLKIIQDEVNS
LayV-C3  1   ---------------------------MATNKRD----VI-KTTESTRED-KVKKFYGVETA----EK----VADSISSNKVFIL--MNTLLILTGAII-TITLNVTNLTTIKNQQA--MLKIIQDEVNS
MojV     1   ---------------------------MATNRDN----TI-TSAEVSQED-KVKKYYGVETA----EK----VADSISGNKVFIL--MNTLLILTGAII-TITLNITNLTAAKSQQN--MLKIIQDDVNA
SHNV5-1  1   MQNRRTSSDPVNTQNSNNRPRIENGGPMNPT-MG----NQ-PRIQTARSD-PARKYYGVETA----ER----VADSISSNKVFIL--MNTLLILTGAII-TITLNITNLNNAQNQQT--MLKMIQDEVDL
SHNV5-2  1   MQNRRTSSDLANTQNSNNRPRTENGGPMNST-MG----NQ-PRIQTARSD-PVRKYYGVETA----ER----VADSISSNKVFIL--MNTLLILTGAII-TITLNITNLNNAQNQQT--MLKMIQDEVDL
DarV-K   1   ------------------------MDKIMANKN-----NN-NNKVNTRET-TVKKFYGVDTA----EK----VADSIISNKIFIL--INTLLIITSSVI-TITLNITNLEGIKNQNA--IIKTLQMEIMN
DarV-Cβ  1   ------------------------MDKIMANKN-----NN-NNKVNTRET-TVKKFYGVDTA----EK----VADSIISNKIFIL--INTLLIITSSVI-TITLNITNLEGIKNQNA--IIKTLQMEIMN
DewV-1   1   -----------------------------MANT-----NT-SNKVQTRES-TVKKFYGVDTA----EK----VADSISNNKLFIL--ANSLLILTSTVI-IITLNITNLEGIKHQNQ--VIKTLHDEIQA
MelV     1   ---------------------------MSGPAK-----QN-NPKINTRET-NVKKFYGVETA----EK----VADSISNNKLFIL--ANTLLILTSTVI-IITLNITNLEGIKHQNE--VIRNLHDEIKS
NinV     1   ------------------------MTQISTGSER----TM-STEKQEFEN-AARRYYGVEKA----EN----LAGTILNNKMFIL--MNTLLIITGSLI-TIGLNATNLTNISYQKT--LVNNLDKEMRH
SHNV2    1   ------------------------------MTDD----NK-MVKVGTETT-TPHKYYGVEKV----DN----FAGKVVSNRIFIL--TNMILIITSSIV-AITLNVTNLESLYNQRE--TLAMVNQEVHN
GakV-1   1   -------------------------------MSP----KI-DTNKTAPIS-DSRKYYGVDSV----ER----FADGVTNNKIFIL--ANMLLIIMSSIV-MISLNITNLENLNNQKT--SLKILNEDVTN
GakV-2   1   -------------------------------MSP----KI-DTNKTAPIS-DSRKYYGVDSV----ER----FADGVTNNKIFIL--ANMLLIIMSSIV-MISLNITNLENLNNQKT--SLKILNEDVTN
SHNV3    1   --------------------------------MS----KV-DTSKTTSTT-NSRKYYGVESV----EK----FADGVTNNKIFIL--ANMLLIIMSSIV-MISLNITNLENLNNQKT--SLKILNDDVNN
SHNV11   1   --------------------------------MS----KV-DTSKTTSTT-NSRKYYGVESV----EK----FADGVTNNKIFIL--ANMLLIIMSSIV-MISLNITNLENLNNQKT--SLKILNEDVNN

NiV-M1   90  IQQQIKGLADKIGTEIGPKVSLIDTSSTITIPANIGLLGSKISQSTASINENVNEKCKF---TLPPLKIHEC-----NIS-CPNPLPFREYRPQTEGVSNLVGLPNNICLQKTSNQI-------------
NiV-M3   90  IQQQIKGLADKIGTEIGPKVSLIDTSSTITIPANIGLLGSKISQSTASINENVNEKCKF---TLPPLKIHEC-----NIS-CPNPLPFREYRPQTEGVSNLVGLPNNICLQKTSNQI-------------
NiV-B1.5 90  IQQQIKGLADKIGTEIGPKVSLIDTSSTITIPANIGLLGSKISQSTASINENVNEKCKF---TLPPLKIHEC-----NIS-CPNPLPFREYKPQTEGVSNLVGLPNNICLQKTSNQI-------------
NiV-B1.6 90  IQQQIKGLADKIGTEIGPKVSLIDTSSTITIPANIGLLGSKISQSTASINENVNEKCKF---TLPPLKIHEC-----NIS-CPNPLPFREYKPQTEGVSNLVGLPNNICLQKTSNQI-------------
NiV-B2.4 90  IQQQIKGLADKIGTEIGPKVSLIDTSSTITIPANIGLLGSKISQSTASINENVNEKCKF---TLPPLKIHEC-----NIS-CPNPLPFREYKPQTEGVSNLVGLPNNICLQKTSNQI-------------
NiV-B3.1 90  IQQQIKGLADKIGTEIGPKVSLIDTSSTITIPANIGLLGSKISQSTASINENVNEKCKF---TLPPLKIHEC-----NIS-CPNPLPFREYKPQTEGVSNLVGLPNNICLQKTSNQI-------------
NIV-I1.3 90  IQQQIKGLADKIGTEIGPKVSLIDTSSTITIPANIGLLGSKISQSAASINENVNEKCKF---TLPPLKIHEC-----NIS-CPNPLPFREYKPQTEGVSNLVGLPNNICLQKTSNQI-------------
HeV-α1.2 90  VQQQIKALTDKIGTEIGPKVSLIDTSSTITIPANIGLLGSKISQSTSSINENVNDKCKF---TLPPLKIHEC-----NIS-CPNPLPFREYRPISQGVSDLVGLPNQICLQKTTSTI-------------
HeV-α3   90  VQQQIKALTDKIGTEIGPKVSLIDTSSTITIPANIGLLGSKISQSTSSINENVNDKCKF---TLPPLKIHEC-----NIS-CPNPLPFREYRPISQGVSDLVGLPNQICLQKTTSTI-------------
HeV-α4   90  VQQQIKALTDKIGTEIGPKVSLIDTSSTITIPANIGLLGSKISQSTSSINENVNDKCKF---TLPPLKIHEC-----NIS-CPNPLPFREYRPISQGVSDLVGLPNQICLQKTTSTI-------------
HeV-α5   90  VQQQIKALTDKIGTEIGPKVSLIDTSSTITIPANIGLLGSKISQSTSSINENVNDKCKF---TLPPLKIHEC-----NIS-CPNPLPFREYRPISQGVSDLVGLPNQICLQKTTSTI-------------
HeV-α6   90  VQQQIKALTDKIGTEIGPKVSLIDTSSTITIPANIGLLGSKISQSTSSINENVNDKCKF---TLPPLKIHEC-----NIS-CPNPLPFREYRPISQGVSDLVGLPNQICLQKTTSTI-------------
HeV-β1   89  VQQQIKALTDKIGTEIGPKVSLIDTSSTITIPANIGLLGSKISQSTSSINENVNDKCKF---TLPPLKIHEC-----NIS-CPNPLPFREYRPISQGVSDLVGLPNQICLQRTTSTI-------------
HeV-β3   89  VQQQIKALTDKIGTEIGPKVSLIDTSSTITIPANIGLLGSKISQSTSSINENVNDKCKF---TLPPLKIHEC-----NIS-CPNPLPFREYRPISQGVSDLVGLPNQICLQRTTSTI-------------
CedV-1   113 IQDSLSSLTNMINTEITPRIGILVTATSVTLSSSINYVGTKTNQLVNELKDYITKSCGF---KVPELKLHEC-----NIS-CADPKISKSAMYSTNAYAELAGPPKIFCKSVSKDPD-------------
GhV      106 LNLNLNQLTNKIQREIIPRITLIDTATTITIPSAITYILATLTTRISELLPSINQKCEF---KTPTLVLNDC-----RIN-CTPPLNPSDGVKMSSLATNLVAHGPSPCRNFSSVPT-------------
AngV     87  AIGKINELYSLINNELKIRMMVMDTAVNVDIPADISALSMKLTSAITSIEELIQSQTSSVPSNRPSMNQSGGSSGSSQLSNSSRPIPD-QGNPNY----CNYDKPNRPSMSFTSLNIITPTESAYTEGCH
LayV-C2  85  KLEMFVSLDQLVKGEIKPKVSLINTAVSVSIPGQISNLQTKFLQKYVYLEESITKQCTCNPLSG--------------IFPTTKP--P--PPPTDKPDDDT---TDD----DKVDTTIKPVEYYKPDGCN
LayV-C3  85  KLEMFVSLDQLVKGEIKPKVSLINTAVSVSIPGQISNLQTKFLQKYVYLEESITKQCTCNPLSG--------------IFPTTKP--P--PPPTDKPDDDT---TDD----DKVDTTIKPVEYYKPDGCN
MojV     85  KLEMFVNLDQLVKGEIKPKVSLINTAVSVSIPGQISNLQTKFLQKYVYLEESITKQCTCNPLSG--------------IFPTSGP--T--YPPTDKPDDDT---TDD----DKVDTTIKPIEYPKPDGCN
SHNV5-1  111 KLSAFDNLEQLVKGDLKPKVTLINSAVSVSIPSQISNLQNKLTQRISSLEDMVGRKCNFNPLAT--------------IFPTVEP--S--KPPTTEEDEDT---SDD----DKVDISIKPIDYKSPINCS
SHNV5-2  111 KLSVFDNLEQLVKGDLKPKVTLINSAVSVSIPSQISNLQNKLTQRISNLEDMVGRKCNFNPLAT--------------IFPTVEP--S--KPPTTEEDEDT---SDD----DKVDISIRPIDYKSPMNCS
DarV-K   87  KVETISKLEQIVKGDLKPKVTLINSAVSVSIPGQISNLQTKMSQKLKFIEDSVIDQCTCNPLSG--------------IFPSKEP--PKEPHPGTEDDDDT---SDD----DKVDDSIQSFDYPSFTVCN
DarV-Cβ  87  KVETISKLEQIVKGDLKPKVTLINSAVSVSIPSQISNLQTKMSQKLKFIEDSVIDQCTCNPLSG--------------IFPSKEP--PKEPHPGTEDDDDT---SDD----DKVDDSIQSFDYPGFTVCN
DewV-1   82  KIDVISQLETIIKGDLKPKISLINSAVSVSLPGQITNLQTKLNQKIKLLEDQVQDQCTCNPLSG--------------IFPSRPP--QPTPSPDDESDDDM---SDD----DKVDDSIQGIDYPNFMICN
MelV     84  KIETINNLEQIIKGDLKPKISLINSAVSVSLPGQISNLQNKMLQKFKTLEDSVIDQCTCNPLSG--------------IFPSKPP--HTSKPDDEEDDNDF---TDD----DKVDDTIQGIDYPTFTTCN
NinV     88  KFDVVSQLDEFLRGEIRPKINIISSATTVSLPTQLSQLSQRLMNQIAKLESMIVAQCSGATSTG--------------APANDPN-----KKPGEVDVDAPTLQPPE----DDDQDSIQMINLSEYPMCS
SHNV2    82  KLGDVSTIGNTIKGDIKPKVNLINSAVSVSLPAQMSSMLQKMLNKMNGIEDAIDQRCSCSTLGS--------------LLPGSSQRPPTVSTPGAGDDDES---TDT----D--SGSVTMIDLSAFTKCT
GakV-1   81  KLEQIDTLDQIVKGEIKPKLTLVNSAVSVTIPSQISNLATKLNNQIKTLEVAVNKQCSGNILGG--------------LLTPKPPTGSGTTSPDDGNPDDN-----V----DDLEGDMRPLNLAELNDCS
GakV-2   81  KLEQIDTLDQIVKGEIKPKLTLVNSAVSVTIPSQISNLATKLNNQIKTLEVAVNKQCSGNILGG--------------LLTPKPPTGSGTTSPDDGNSDDN-----V----DDLEGDMRPLNLAELNDCS
SHNV3    80  KLEQIDTLDQIVKGEIKPKLTLVNSAVSVTIPSQLSNLATKLNNQIKTLEVAVNKQCSGNILGG--------------LLTPKPPEGGDVINPDDNSTGDN-----T----DDLEGDMRPLNLAQLNDCS
SHNV11   80  KLEQIDTLDQIVKGEIKPKLTLVNSAVSVTIPSQLSNLATKLNNQIKTLEVAVNKQCSGNILGG--------------LLTPKPPEGGDVINPDDNSTGDN-----T----DDLEGDMRPLNLAQLNDCS

NiV-M1   197 ---------------LKPKLISYTLPVVGQSGTCITDPLLAMDEGYFAYSHLERIGSCSRGVSKQRIIGVGEVLDRGDEVPSLFMTNVWTPPNPNTVYHCSAVYNNEFYYVLCAVSTVGDPILNSTYWSG
NiV-M3   197 ---------------LKPKLISYTLPVVGQSGTCITDPLLAMDEGYFAYSHLERIGSCSRGVSKQRIIGVGEVLDRGDEVPSLFMTNVWAPPNPNTVYHCSAVYNNEFYYVLCAVSTVGDPILNSTYWSG
NiV-B1.5 197 ---------------LKPKLISYTLPVVGQSGTCITDPLLAMDEGYFAYSHLEKIGSCSRGVSKQRIIGVGEVLDRGDEVPSLFMTNVWTPSNPNTVYHCSAVYNNEFYYVLCAVSVVGDPILNSTYWSG
NiV-B1.6 197 ---------------LKPKLISYTLPVVGQSGTCITDPLLAMDEGYFAYSHLEKIGSCSRGVSKQRIIGVGEVLDRGDEVPSLFMTNVWTPSNPNTVYHCSAVYNNEFYYVLCAVSVVGDPILNSTYWSG
NiV-B2.4 197 ---------------LKPKLISYTLPIVGQSGTCITDPLLAMDEGYFAYSHLEKIGSCSRGVSKQRIIGVGEVLDRGDEVPSLFMTNVWTPSNPNTVYHCSAVYNNEFYYVLCAVSVVGDPILNSTYWSG
NiV-B3.1 197 ---------------LKPKLISYTLPVVGQSGTCITDPLLAMDEGYFAYSHLEKIGSCSRGVSKQRIIGVGEVLDRGDEVPSLFMTNVWTPSNPNTVYHCSAVYNNEFYYVLCAVSVVGDPILNSTYWSG
NIV-I1.3 197 ---------------LKPKLISYTLPVVGQSGTCITDPLLAMDEGYFAYSHLEKIGSCSRGVSKQRIIGVGEVLDRGDEVPSLFMTNVWTPSNPNTVYHCSAVYNSEFYYVLCAVSVVGDPILNSTYWSG
HeV-α1.2 197 ---------------LKPRLISYTLPINTREGVCITDPLLAVDNGFFAYSHLEKIGSCTRGIAKQRIIGVGEVLDRGDKVPSMFMTNVWTPPNPSTIHHCSSTYHEDFYYTLCAVSHVGDPILNSTSWTE
HeV-α3   197 ---------------LKPRLISYTLPINTREGVCITDPLLAVDNGFFAYSHLEKIGSCTRGIAKQRIIGVGEVLDRGDKVPSMFMTNVWTPPNPSTIHHCSSTYHEDFYYTLCAVSHVGDPILNSTSWTE
HeV-α4   197 ---------------LKPRLISYTLPINTREGVCITDPLLAVDNGFFAYSHLEKIGSCTRGIAKQRIIGVGEVLDRGDKVPSMFMTNVWTPPNPSTIHHCSSTYHEDFYYTLCAVSHVGDPILNSTSWTE
HeV-α5   197 ---------------LKPRLISYTLPINTREGVCITDPLLAVDNGFFAYSHLEKIGSCTRGIAKQRIIGVGEVLDRGDKVPSMFMTNVWTPPNPSTIHHCSSTYHEDFYYTLCAVSHVGDPILNSTSWTE
HeV-α6   197 ---------------LKPRLISYTLPINTREGVCITDPLLAVDNGFFAYSHLEKIGSCTRGIAKQRIIGVGEVLDRGDKVPSMFMTNVWTPPNPSTIHHCSSTYHEDFYYTLCAVSHVGDPILNSTSWTE
HeV-β1   196 ---------------LKPKLISYTLPINTREGVCITDPLLTIDNGFFAYSHLEKIGSCTRGIAKQRIIGVGEVLDRGDKVPSMFMTNVWTPPNPSTVHHCSSTYHDDFYYTLCAVSHVGDPILNGTSWIE
HeV-β3   196 ---------------LKPKLISYTLPINTREGVCITDPLLTIDNGFFAYSHLEKIGSCTRGIAKQRIIGVGEVLDRGDKVPSMFMTNVWTPPNPSTVHHCSSTYHDDFYYTLCAVSHVGDPILNGTSWIE
CedV-1   220 ---------------FRLKQIDYVIPVQQDRSICMNNPLLDISDGFFTYIHYEGINSCKKSDSFKVLLSHGEIVDRGDYRPSLYLLSSHYHPYSMQVINCVPVTCNQSSFVFCHISNNTKTLDNSDYSSD
GhV      213 ---------------IYYYRIPGLYNRTALDERCILNPRLTISSTKFAYVHSEYDKNCTRGFKYYELMTFGEILEGPEKEPRMFSRSFYSPTNAVNYHSCTPIVTVNEGYFLCLECTSSDPLYKANLSNS
AngV     212 TETVLTSE------QFQLVPLNQLLAGDYGEDYCSRYPSLSLRGRRFTYTEERYRGSCSTRTKEAQIIVYGYITDFGYRHPTLMILDTFNPPNYDMIYRCTSVTKLSTGLVICLYTNMTEFELYESPKNT
LayV-C2  190 KTNDHFTM----QPGVNFYTVPNLGPSSSSADECYTNPSFSIGSSIYMFSQEIRKTDCTTGEILSIQIVLGRIVDKGQQGPQASPLLVWSVPNPKIINSCAVAAGDETGWVLCSVTLTAASGEPIPHMFD
LayV-C3  190 KTNDHFTM----QPGVNFYTVPNLGPSSSSADECYTNPSFSIGSSIYMFSQEIRKTDCTTGEILSIQIVLGRIVDKGQQGPQASPLLVWSVPNPKIINSCAVAAGDETGWVLCSVTLTAASGEPIPHMFD
MojV     190 RTGDHFTM----EPGANFYTVPNLGPASSNSDECYTNPSFSIGSSIYMFSQEIRKTDCTAGEILSIQIVLGRIVDKGQQGPQASPLLVWAVPNPKIINSCAVAAGDEMGWVLCSVTLTAASGEPIPHMFD
SHNV5-1  216 KPNDHPTM----EPGPNFYAVPNLGPSSSNADECYTNPSFSIGSSIYMFSQEIRKTDCIAGELQSVQIILGRIVDKGQQGPQASPLLVWSVPNPRTINSCAVAAGDEMGWVLCSVTSTAASGGPNPHTFD
SHNV5-2  216 KPNDHPTM----EPGPNFYAVPNLGPSSSNADECYTNPSFSIGSSIYMFSQEIRKTDCIAGELQSVQIILGRIVDKGQQGPQASPLLVWSVPNPRTINSCAVAAGDEMGWVLCSVTLTAASGGPSPHTFD
DarV-K   194 NSDETTSI----IPGPNLYAVPNLSIKEDEEDECVTNPSFSVGTSIYMYSQEIRKTDCKSGPLTGIKIILGRIVDKGQFGPQVSPLLVWDVPKPETINSCAVVAEDEVGWALCSITSIANSGEPIPYLLS
DarV-Cβ  194 NSDETTSI----IPGPNLYAVPNLSIKEDEEDECVTNPSFSVGTSIYMYSQEIRKTDCKSGPLTGIKIILGRIVDKGQFGPQVSPLLVWDVPKPETINSCAVVAEDEVGWALCSITSIANSGEPIPYLLS
DewV-1   189 GTRNELSI----TPGQNLYSVPSLTMREDHDDECVVNPSFSVGTSIYMYSQEIRKTDCKQGPLIGIKISLGRVVDKGQSGPQVSPLLVWDVPKPETISSCAVVAGDEIGWALCSTIMTAKSGEPIPYMFE
MelV     191 KSDHSMSI----TPGPNLYTVPSLTYREDNDDECVVNPSFSIGTSIYMYSQEIRKTDCKQGPLVGIKITLGRIVDKGLSGPQVSPLLVWDVPKPELTGSCAVAAGNEVGWVLCSTTLAASSGEPIPNMFE
NinV     195 NQQQYKEPHFNVTVGPIEYT--ELLGLNNSSDYCISSPSLAIEGSTYIYAQQVRKTNCQFGEIVGIQVVMGRLVDKGRTVPSSSPLLRWEVPAPESDQSCAAAASRGTGWLFCVSGNTVSGEFLSDGRLL
SHNV2    189 LDRSKPYAA--YSLTGVAHNIPGLSVNIADNGDCVSNVAFAISEGIYVYTQEIRKTDCLTGQKISQITTLGRIVDKGQSGPRVSPLSTYLEPDPSSKISCSVAASSGSGWVFCVLTTGEGVNYSKFGPFI
GakV-1   188 RYPQTPNPTF--AISPDIHPMPELTEPIQLKNHCAVFPTVALGESIYVYSHQIRKTPCKSEDTIHQRVSLGRIVDRGFSGPRASPLSTWDLQETEFLSSCSVAASGEIGWVLCVTTDKFTRDTVYAGPYT
GakV-2   188 RYPQTPNPTF--AISPDIHPMPELTEPIQLKNHCAVFPTVALGESIYVYSHQIRKTPCKSEDTTHQRVSLGRIVDRGFSGPRASPLSTWDLQETEFLSSCSVAASGEIGWVLCVTTDKFTRDTVYAGPYT
SHNV3    187 NYPQAPQPTF--AIDPDLHPMPELTEPIQLKNHCAVFPTVALGESIYVYSHQIRKTPCKSEDTIHQRVSIGRIVDRGFSGPRASPLSTWDLQETGYLSSCSVAASGETGWVLCVTTDKFTGATVYLGPYT
SHNV11   187 NYPQAPQPTF--AIDPDLHPMPELTEPIQLKNHCAVFPTVALGESIYVYSHQIRKTPCKSEDTIHQRVSIGRIVDRGFSGPRASPLSTWDLQETGYLSSCSVAASGETGWVLCVTTDKFTGTTVYLGPYT

NiV-M1   313 SLMMTRLA--VKPKSNGGGYNQHQLALR-SIEKGRYDKVMPYGPSGIKQGDTLYFPAVGFLVRTEFKYNDSNCPITKCQYS----KPENCRLSMG--IRPNSHYILRSGLLKYNLSDGENPKVVFIEISD
NiV-M3   313 SLMMTRLA--VKPKSNGGGYNQHQLALR-SIEKGRYDKVMPYGPSGIKQGDTLYFPAVGFLVRTEFKYNDSNCPITKCQYS----KPENCRLSMG--IRPNSHYILRSGLLKYNLSDGENPKVVFIEISD
NiV-B1.5 313 SLMMTRLA--VKPKNNGESYNQHQFALR-NIEKGKYDKVMPYGPSGIKQGDTLYFPAVGFLVRTEFKYNDSNCPIAECQYS----KPENCRLSMG--IRPNSHYILRSGLLKYNLSDEENSKIVFIEISD
NiV-B1.6 313 SLMMTRLA--VKPKNNGESYNQHQFALR-NIEKGKYDKVMPYGPSGIKQGDTLYFPAVGFLVRTEFTYNDSNCPIAECQYS----KPENCRLSMG--IRPSSHYILRSGLLKYNLSDEENSKIVFIEISD
NiV-B2.4 313 SLMMTRLA--VKPKNNGESYNQHQFALR-NIEKGKYDKVMPYGPSGIKQGDTLYFPAVGFLVRTEFKYNDSNCPIAECQYS----RPENCRLSMG--IRPNSHYILRSGLLKYNLSDEENSKIVFIEISD
NiV-B3.1 313 SLMMTRLA--VKPKNNGESYNQHQFALR-NIEKGKYDKVMPYGPSGIKQGDTLYFPAVGFLVRTEFKYNDSNCPIAECQYS----KPENCRLSMG--IRPNSHYILRSGLLKYNLSDEENSKIVFIEISD
NIV-I1.3 313 SLMMTRLA--VKPKNNGESYNQHQFALR-NIEKGMYDKVMPYGPSGIKQGDTLYFPAVGFLVRTEFKYNDSNCPVAKCQYS----KPENCRLSMG--IRPNSHYILRSGLLKYNLSDEEDSKIIFIEISD
HeV-α1.2 313 SLSLIRLA--VRPKSDSGNYNQKYIAIT-KVERGKYDKVMPYGPSGIKQGDTLYFPAVGFLPRTEFQYNDSNCPIIHCKYS----KAENCRLSMG--VNSKSHYILRSGLLKYNLSLGGDIILQFIEIAD
HeV-α3   313 SLSLIRLA--VRPKSDNGDYNQKYIAIT-KVERGKYDKVMPYGPSGIKQGDTLYFPAVGFLPRTEFQYNDSNCPIIHCKYS----KAENCRLSMG--VNSKSHYILRSGLLKYNLSLGGDIILQFIEIAD
HeV-α4   313 SLSLIRLA--VRPKSDSGDYNQKYITIT-KVERGKYDKVMPYGPSGIKQGNTLYFPAVGFLPRTEFQYNDSNCPIIHCKYS----KAENCRLSMG--VNSKSHYILRSGLLKYNLSLGGDIILQFIEIAD
HeV-α5   313 SLSLIRLA--VRPKSDSGDYNQKYIAIN-KVERGKYDKVMPYGPSGIKQGDTLYFPAVGFLPRTEFQYNDSNCPIIHCKYS----KAENCRLSMG--VNSKSHYILRSGLLKYNLSLGGDIILQFIEIAD
HeV-α6   313 SLSLIRLA--VRPKSDSGDYNQKYIAIT-KVERGKYDKVMPYGPSGIKQGDTLYFPAVGFLPRTEFQYNDSNCPIIHCKYS----KAENCRLSMG--VNSKSHYILRSGLLKYNLSLGGDITLQFIEIAD
```

```
HeV-β1   312 SLFVIRLA--VRPKTDGGGYNQKYITMT-KIERGKYDKVMPYGPSGIKQGDTLYFPAVGFLPRTEFQYNDSNCPIINCKYS----KAENCRLSMG--INPRSHYVLRSGLLKYNLSLGEDTRLQFIEIAD
HeV-β3   312 SLFMIRLA--VRPKTDGGGYNQKYITMT-KIERGKYDKVMPYGPSGIKQGDTLYFPAVGFLPRTEFQYNDSNCPIINCKYS----KAENCRLSMG--INPRSHYVLRSGLLKYNLSLGEDTRLQFIEIAD
CedV-1   336 EYYITYFNGIDRPKTK-------KIPINNMTADNRYIHFTFSGGGGVCLGEEFIIPVTTVINTDVFT--HDYCESFNCSVQTGKSLKEICSESLR--SPTNSSRYNLNGIMIISQNNMTDFKIQLNGITY
GhV      329 TFHLVILRHNKDE-------KIVSMPSFNLSTDQEYVQIIPAEGGGTAESGNLYFPCIGRLLHKRVT--HPLCKKSNCSRT----DDESCLKSYY--NQGSPQHQVVNCLIRIRNAQRDNPTWDVITVDL
AngV     335 --DLIIFEAFDGSQHKGTILN-----SSKFIYLNDIDMFIPGTGGGIYLNKTIYLPGIVFTTNPPDG--NAGCPNSACTNK----DPKLCRKGLK--YIFNNSYYMVNGFALISQKGRKEYEVTVMIET
LayV-C2  316 GFWLYKFE-------PDTEVVAYRITGFAYLLDKVYDSVFIGKGGGIQRGNDLYFQMFGLSRNRQSI--KALCEHGSCLGTGGGGYQVLCDRAVM--SFGSEESLISNAYLKVNDVASGKPTIISQTFPP
LayV-C3  316 GFWLYKFE-------PDTEVVAYRITGFAYLLDKVYDSVFIGKGGGIQRGNDLYFQMFGLSRNRQGI--KALCEHGSCLGTGGGGYQVLCDRAVM--SFGSEESLISNAYLKVNDVASGKPTIISQTFPP
MojV     316 GFWLYKLE-------PDTEVVSYRITGYAYLLDKQYDSVFIGKGGGIQKGNDLYFQMYGLSRNRQSF--KALCEHGSCLGTGGGGYQVLCDRAVM--SFGSEESLITNAYLKVNDLASGKPVIIGQTFPP
SHNV5-1  342 GFWLYRLT-------PDTDVVSYRISGFAYLLDKVYDSVYIGKGGGVQKGSDLYFQMYGISRNRASF--TALCDHGSCPGVGGGGYQTLCDKSMT--VLGSDESLVTSAYLRVGDIYSGKPVIVSQTFPP
SHNV5-2  342 GFWLYRLT-------PDTDVVSYRISGFAYLLDKVYDSVYIGKGGGVQKGSDLYFQMYGISRNRASF--TALCDHGSCPGVGGGGYQTLCDKSMT--VLGSDESLVTSAYLRVGDIYSGKPVIVSQTFPP
DarV-K   320 GFKLFKFE-------PDSEVVQYVLGSKSFRLDRVYHTLYVGKGGGAARYDGLYFEGFGVITNIGKD--QPLCNHGKCSGS--GSYIPICLSAMS--MMGDTDFLVVSNIIKVVDANIGKPVITVQTFKI
DarV-Cβ  320 GFKLFKFE-------PDSEVVQYVLGSKSFRLDRVYHTLYVGKGGGAVRYDGLYFEGFGVITNIGKD--QPLCNHEKCSGS--GSYTPICLSAMS--MMGDTDFLVVSNIIKVVDTNIGKPVITVQTFKI
DewV-1   315 GFKLFKFE-------PDKDVVQYTLAPRAFRLDEKYHTLNIGKGGGIIRGEELYFTGFGIITNYKKV--DPLCNHGICPGS--GSYLPVCKSAIS--FMGDEDLIVVDNIIKVVNVDIGTPVIDLKTFKA
MelV     317 GFKLYKFE-------PEKEVVQYIITKRAFRLDNPYHTMNIGKGGGIIKEDGLYFQGFGVIYNSGKV--DPLCDHGKCPGV--GSYMPTCLSAMS--MMGDVDFVVVDNIIKVVDYDIGKPVIDLKTFKI
NinV     323 GVRLFTFK-------GGSLYHAVYVPQTSIRFSLPIKTFMVGTGGGVYVDGKLYFVGYGTVTAPHTG--GYLCSFDDCPGITDA--GTKCGDALFPPISYDGKAMLVNCLVKVSLDNPEVPSIEVDFFHP
SHNV2    317 GFTLSHIT-------EDGDVTTYNISLNVVNSNYAFKSIFIGPGNGIVRGNNMYFVGYGALRDNTNY--PSWCIYDGCPAD--PSYNKYCEWSMR--PDYFGRLPGLNCIITITNYKESNRGILIRAIPN
GakV-1   316 GLKLYKLS-------IRGQKEEYSITANNITTDAVIIALTLTRGSGVPKNNKLIFLGLAAVRDVDTT--GVLCPTWKCENINN--NVGSCVHSYR--LTADMNNYFMNVVVAVDVTPTGKMTASVSLLPM
GakV-2   316 GLKLYKLS-------IRGQKEEYSITANNITTDAVIIALTLTRGSGVPKNNKLIFLGLAAVRDVDTT--GVLCPTWKCDNINN--NVGSCVHSYR--LTADMNNYFMNVVVAVDVTPTGKMTASVSLLPM
SHNV3    315 GLKLYKLS-------VRGQKEEYAITANDITSDATLIALTLTRGSGVSKNNKLIFLGIAAVRDADRT--GVLCPDWKCDNINN--NINSCVHSYR--LTSDNNNYFMNVVVTVDVSPAGKMIASVSLLPM
SHNV11   315 GLKLYKLS-------VRGQKEEYAITASDITADTTVIALTLTRGSGVSKNNKLIFLGLAAVRDADRT--GVLCPDWKCDNINN--NINSCVHSYR--LTSDNNNYFMNVVVTVDVSPAGKMIASISLLPM

NiV-M1   434 QRLSIGSPSKIYDSLGQPVFYQASFSWDTMIKFGDVL--TVNPLVVNWRNNTVISRP--GQSQCPRFNTCPEICWEGVYNDAFLIDRINWIS--AGVFLDSNQTAENPVFTVFKDNEILYRAQLASEDTN
NiV-M3   434 QRLSIGSPSKIYDSLGQPVFYQASFSWDTMIKFGDVL--TVNPLVVNWRNNTVISRP--GQSQCPRFNTCPEICWEGVYNDAFLIDRINWIS--AGVFLDSNQTAENPVFTVFKDNEILYRAQLASEDTN
NiV-B1.5 434 QRLSIGSPSKIYDSLGQPVFYQASFSWDTMIKFGDVQ--TVNPLVVNWRDNTVISRP--GQSQCPRFNKCPEICWEGVYNDAFLIDRINWIS--AGVFLDSNQTAENPVFTVFKDNEVLYRAQLASEDTN
NiV-B1.6 434 QRLSIGSPSKIYDSLGQPVFYQASFSWDTMIKFGDVQ--TVNPLVVNWRDNTVISRP--GQSQCPRFNKCPEVCWEGVYNDAFLIDRINWIS--AGVFLDSNQTAENPVFTVFKDNEVLYRAQLASEDTN
NiV-B2.4 434 QRLSIGSPSKIYDSLGQPVFYQASFSWDTMIKFGDVQ--TVNPLVVNWRDNTVISRP--GQSQCPRFNKCPEVCWEGVYNDAFLIDRINWIS--AGVFLDSNQTAENPVFTVFKDNEVLYRAQLASEDTN
NiV-B3.1 434 QRLSIGSPSKIYDSLGQPVFYQASFSWDTMIKFGDVQ--TVNPLVVNWRDNTVISRP--GQSQCPRFNKCPEVCWEGVYNDAFLIDRINWIS--AGVFLDSNQTAENPVFTVFKDNEVLYRAQLASEDTN
NIV-I1.3 434 QRLSIGSPSKIYDSLGQPVFYQASFSWDTMIKFGDVQ--TVNPLVVNWRDNTVISRP--GQSQCPRFNTCPEVCWEGVYNDAFLIDRINWIS--AGVFLDSNQTAENPVFTVFKDNEVLYRAQLASEDTN
HeV-α1.2 434 NRLTIGSPSKIYNSLGQPVFYQASYSWDTMIKLGDID--TVDPLRVQWRNNSVISRP--GQSQCPRFNVCPEVCWEGTYNDAFLIDRLNWVS--AGVYLNSNQTAENPVFAVFKDNEILYQVPLAEDDTN
HeV-α3   434 NRLTIGSPSKIYNSLGQPVFYQASYSWDTMIKLGDVD--TVDPLRVQWRNNSVISRP--GQSQCPRFNVCPEVCWEGTYNDAFLIDRLNWVS--AGVYLNSNQTAENPVFAVFKDNEILYQVPLAEDDTN
HeV-α4   434 NRLTIGSPSKIYNSLGQPVFYQASYSWDTMIKLGDVD--TVDPLRVQWRNNSVISRP--GQSQCPRFNVCPEVCWEGTYNDAFLIDRLNWVS--AGVYLNSNQTAENPVFAVFKDNEILYQVPLAEDDTN
HeV-α5   434 NRLTIGSPSKIYNSLGQPVFYQASYSWDTMIKLGDVD--TVDPLRVQWRNNSVISRP--GQSQCPRFNVCPEVCWEGTYNDAFLIDRLNWVS--AGVYLNSNQTAENPVFAVFKDNEILYQVPLAEDDTN
HeV-α6   434 NRLTIGSPSKIYNSLGQPVFYQASYSWDTMIKLGDVD--TVDPLRVQWRNNSVISRP--GQSQCPRFNVCPEVCWEGTYNDAFLIDRLNWVS--AGVYLNSNQTAENPVFAVFKDNEILYQVPLAEDDTN
HeV-β1   433 NRLTIGSPSKIYNSLGQPVFYQASYSWDTMIKLGDVD--TVDPLKVQWRNNSVISRP--GQSQCPRFNVCPEVCWEGTYNDAFLIDRLNWIS--AGVYLNSNQTAENPVFAVFKDNEILYQVQLADEDTN
HeV-β3   433 NRLTIGSPSKIYNSLGQPVFYQASYSWDTMIKLGDVD--TVDPLKVQWRNNSVISRP--GQSQCPRFNVCPEVCWEGTYNDAFLIDRLNWIS--AGVYLNSNQTAENPVFAVFKDNEILYQVQLADEDTN
CedV-1   455 NKLSFGSPGRLSKTLGQVLYYQSSMSWDTYLKAGFVE--KWKPFTPNWMNNTVISRP--NQGNCPRYHKCPEICYGGTYNDIAPLDLGKDMY--VSVILDSDQLAENPEITVFNSTTILYKERVSKDELN
GhV      444 TNTYPGSRSRIFGSFSKPMLYQSSVSWHTLLQVAEIT--DLDKYQLDWLDTPYISRP--GGSECPFGNYCPTVCWEGTYNDVYSLTPNNDLF--VTVYLKSEQVAENPYFAIFSRDQILKEFPLDAWISS
AngV     451 DVYHFGSRSRVQYFNSRAKYYQAPNGWFSYPIYGNIYFEKNQKITFDEANYTVYERY--TEGSCPINSSCPAFCHSSGYNDIWANDIDSNIT--FGIYNSDSHSYGRPVFFMANQSGIMYEFVIPGSIIA
LayV-C2  435 SDSYKGSNGRIYTIGERYGIYLAPSSWNRYLRFGLTP--DISVRSTTWLKEKDPIMKVLTTCTNTDKDMCPEICNTRGYQDIFPLSEDSSFYTYIGITPSN--EGTKSFVAVKDDAGHVASITILPNYYS
LayV-C3  435 SDSYKGSNGRIYTIGERYGIYLAPSSWNRYLRFGLTP--DISVRSTTWLKEKDPIMKVLTTCTNTDKDMCPEICNTRGYQDIFPLSEDSSFYTYIGITPSN--EGTKSFVAVKDDAGHVASITILPNYYS
MojV     435 SDSYKGSNGRMYTIGDKYGLYLAPSSWNRYLRFGITP--DISVRSTTWLKSQDPIMKILSTCTNTDRDMCPEICNTRGYQDIFPLSEDSEYYTYIGITPNN--GGTKNFVAVRDSDGHIASIDILQNYYS
SHNV5-1  461 SDSYRGSNGRIYLIGDSYGIYLAPSSWNRYLRFGVTP--DISVKSTRWLDAKYPVMYIRSTCTNKVSNMCPEICSTRGYQDIFPLSADSESYTYIGISPKG--EGTSSFVAVRDRDGHHASKEILSSYFS
SHNV5-2  461 SDSYRGSNGRIYLIGDSYGIYLAPSSWNRYLRFGVTP--DISVKSTRWLDAKYPVMYIRSTCTNKVSNMCPEICSTRGYQDIFPLSADSESYTYIGISPKG--EGTSSFVAVRDRDGHHASKEILSSYFS
DarV-K   437 QDTYKGSHGRIYQMDNNYGIYLASSSWNRYLKFGTTN--TLIPTEVEWSKLKDPIMDIQNNCHNTNQDMCPAVCSSYGYEDIFPLNIEGNAMTYMSMSRNG--EGTSNFIVARSQDNYVNTMKALGEYFK
DarV-Cβ  437 QDTYKGSHGRIYQMDNNYGVYLASSSWNRYLKFGTTN--TLIPTEVEWSKLKDPIMDIQNNCHNTNQDMCPAVCSSYGYEDIFPLNIEGNAMTYMSMSRNG--EGTSNFIVARSQDNYVNTMKALGEYFK
DewV-1   432 SDTYRGSNGRLYAMGDNFGVYFSPSSWNRYLRFGTSS--SPVPGTFKWSPLKSPVLAVGTTCHNDNLSMCPQVCHSYGYEDVFPLDVDGTSMTYMSISTDG--QRSKAYVQARSKDGFVNSIPALTDYYN
MelV     434 SETYRGSNGRIYDMGSKFGLYLASSSWNRFLKFGTSN--DMNPDSFKWANFKTPIMNVQTSCRNTNENMCPQVCYGTGYEDIFPLNDDGSSRTYTTITTNG--SRTQVYIRARVNGNNVGSIQVLNDYYF
NinV     442 GDTYAGSNGRVYKMDRGIGIYISPASWNPHPRFGITR--RKSISDISWIGYASQSRV--STTNCPSNVACPAVCYYPNYFDIYPLDEEGELMTITNYNNPDQADRGDLYTCISTDQEDLNRVLDSNVKRE
SHNV2    434 TNSYPAAGGRLYDLGDRVGLYFTTASWQGKLQFALFE--DFPSYKPIYVNS--DIITVRAHGSCRDKNNCPQDCYSPLYADIYPLNLEATLFTSTPIFMGG--DNADYRTVIINSNKIESLVQMFVNYKS
GakV-1   433 SESYIGSEGGVIDKPGGYGLMISNKGWFARIRYGQTD--RASPQRYEWTDYMSFETP--YYLYCSGGRICPVSCKTWNFTVPTILNPSGSII--IGVKAKSKTGNSASMITINTPDEVIDQYEVFNDYQS
GakV-2   433 SESYIGSEGGVIDKPGGYGLMISNKGWFARIRYGQTD--RASPQRYEWTDYMSFETP--YYLYCSGGRICPVSCKTWNFTVPTILNPSGSII--IGVKAKSKTGNSASMITINTPDEVIDQYEVFNDYQS
SHNV3    432 SESYIGSEGSVIDKPGGYGLMISNKGWFARIRYGQTE--RTSPQRYDWTDYMSFETP--YYLYCSGGRICPSSCRTWNFTVPTILNPSGEII--IGVKAKSRTGNSMSMISINTPNEVIDQYEVFTDYLS
SHNV11   432 SESYIGSEGSVIDKPGGYGLMISNKGWFARIRYGQTE--RTSPQRYEWTDYMSFETP--YYLYCSGGRMCPSSCRTWNFTVPTILNPSGEII--IGVKAKSKTGNSMSMISINTPNEVIDQYEVFKDYLS

NiV-M1   558 AQKTITNCFLLKNKIWCISLVEIYDTGDNVIRPKLFAVKI--PEQCT-------------------------------------------------------------------
NiV-M3   558 AQKTITNCFLLKNKIWCISLVEIYDTGDNVIRPKLFAVKI--PEQCT-------------------------------------------------------------------
NiV-B1.5 558 AQKTITNCFLLRNKIWCISLVEIYDTGDNVIRPKLFAVKI--PEQCT-------------------------------------------------------------------
NiV-B1.6 558 AQKTITNCFLLKNKIWCISLVEIYDTGDNVIRPKLFAVKI--PEQCT-------------------------------------------------------------------
NiV-B2.4 558 AQKTITNCFLLKNKIWCISLVEIYDTGDNVIRPKLFAVKI--PEQCT-------------------------------------------------------------------
NiV-B3.1 558 AQKTITNCFLLKNKIWCISLVEIYDTGDNVIRPKLFAVKI--PEQCT-------------------------------------------------------------------
NIV-I1.3 558 AQKTITNCFLLKNKIWCISLVEIYDTGDNVIRPKLFAVKI--PEQCT-------------------------------------------------------------------
HeV-α1.2 558 AQKTITDCFLLENVIWCISLVEIYDTGDSVIRPKLFAVKI--PAQCSES-----------------------------------------------------------------
HeV-α3   558 AQKTITDCFLLENVIWCISLVEIYDTGDSVIRPKLFAVKI--PAQCSES-----------------------------------------------------------------
HeV-α4   558 AQKTITDCFLLENVIWCISLVEIYDTGDSVIRPKLFAVKI--PAQCSES-----------------------------------------------------------------
HeV-α5   558 AQKTITDCFLLENVIWCISLVEIYDTGDSVIRPKLFAVKI--PAQCSES-----------------------------------------------------------------
HeV-α6   558 AQKTITDCFLLENVIWCISLVEIYDTGDSVIRPKLFAVKI--PAQCSES-----------------------------------------------------------------
HeV-β1   557 AQRTITDCFLLDNVIWCISLVEIYDTGDSVIRPKLFAVKI--PAQCSGN-----------------------------------------------------------------
HeV-β3   557 AQRTITDCFLLDNVIWCISLVEIYDTGDSVIRPKLFAVKI--PAQCSGN-----------------------------------------------------------------
CedV-1   579 TRSTTTSCFLFLDEPWCISVLETNRFNGKSIRPEIYSYKI--PKYC--------------------------------------------------------------------
GhV      568 ARTTTISCFMFNNEIWCIAALEITRLNDDIIRPIYYSFWL--PTDCRTPYP--------------HTGKMTRVPLRSTYNY---------------------------------
AngV     577 AGQSTTSCLLFYQEVWCMNVIEIRKTLNSS--QEIYGFWTKTPLVCPTKDQYDIAHAFPLPTPSPAPAPSQSVPVTEASTITLIQSVEPQGTPTDIPSTTNRDDIQTITNATSG
LayV-C2  561 ITSATISCFMYKEEIWCIAVTEGRKQKENPQRIYAHSYRV--QKMCFNIKPASVVTS--------LPSNVTI----RS------------------------------------
LayV-C3  561 ITSATISCFMYKEEIWCIAVTEGRKQKENPQRIYAHSYRV--QKMCFNIKPASVVTS--------LPSNVTI----RS------------------------------------
MojV     561 ITSATISCFMYKDEIWCIAITEGKKQKDNPQRIYAHSYKI--RQMCYNMKSATVTVG--------NAKNITI----RRY-----------------------------------
SHNV5-1  587 ILSSTISCFTYKEEIWCIAVSEGKRTKGSAQRIFAHSYRV--IRGCGHNLVYPNMTD--------IVPSFSK------------------------------------------
SHNV5-2  587 ILSSTISCFTYKEEIWCIAVSEGKRTKGSAQRIFAHSYRV--IRGCGHNLVYPNMTD--------IVSSFSK------------------------------------------
DarV-K   563 IITATTSCFTFKDEAWCIVVNEGQHKKGDKQRIFAHSYKL--KKSCIKRSKITDLATTSI-----TFNNRSI----R-------------------------------------
DarV-Cβ  563 IITATTSCFTFKDEAWCIVVNEGQHKKGDKQRIFAHSYKL--KKSCIKRSRITDLATTSI-----TINNRSI----R-------------------------------------
DewV-1   558 IYTSTISCFMYRNDVWCIVINEASLKKNDKQRIYAHSYKL--KKECVKSDMLMTDMV---T--LMQNNKSRT----R-------------------------------------
MelV     560 VYSATTSCFMFKNEGWCIVINEAKLNKNDKQRIYAHSYRL--KRLCMSGMRSINSTTGLTT--VVLPSNRRT----R-------------------------------------
NinV     568 VTYTTTSCFVYRRDVWCIVIIVEKSEVREYTEIKPYIYKI--KKSCSLHQGHDAGLRDEEVKVVPLPTEKPKPEARSSYSFWF-------------------------------
SHNV2    558 IMGTTTTCFMYKDEPWCIVTVRGVRKGYSGVEAISYSYKI--KKTCQNTQPKIGNISIPD-----FPSIKPV----R-------------------------------------
GakV-1   557 IGSTITKCFSYKRQPWCLVLLEGVLKSTGVTETSIQTFKI--FRSCVKHRTYLDSLGTRFYYTVSDNGNKTK----QTYIPGSDT-----------------------------
GakV-2   557 IGSTITKCFSYKRQPWCLVLLEGVLKSTGVTETSIQTFKI--FRSCVKHRTYLDSLGTRFYYTVSDNGNKTK----QTYIPGSDT-----------------------------
SHNV3    556 IGSTITKCFLYKRQPWCLVLLEGVPKSTGLTETSIQTFRI--YKSCVKSRTYLDSLGSRYYYVSDQGGNRTR----QAYKLTSDT-----------------------------
SHNV11   556 IGSTITKCFLYKRQPWCLVLLEGVPKSTGQTETSIQTFRI--YKSCVKSRTYLDSLGARYFYVPDQESNRTR----QAYKLTSDT-----------------------------
```
